# Intrinsic IL-6 expression reduces rhIL-6-induced JAK/STAT activation and promotes glucose and oleic acid oxidation in cultured human myoblasts

**DOI:** 10.64898/2026.05.06.722928

**Authors:** Anja Srpčič, Katarina Miš, Barbara Žvar Baškovič Gantar, Klemen Dolinar, Ulrik Nygaard Mjaaseth, Arild C. Rustan, Eili Tranheim Kase, Katja Lakota, Katja Perdan Pirkmajer, Sergej Pirkmajer

## Abstract

Interleukin-6 (IL-6), produced by skeletal muscle and extramuscular tissues, regulates skeletal muscle function through the Janus kinase/signal transducer and activator of transcription (JAK/STAT) pathway. However, the interaction between intrinsic (locally produced) IL-6 and extrinsic (circulating) IL-6 in skeletal muscle remains unclear. We investigated whether and how intrinsic expression of IL-6 in cultured primary human myoblasts influences their response to extrinsic stimulation with recombinant human IL-6 (rhIL-6). Using gene silencing, we found that suppression of intrinsic IL-6 enhanced rhIL-6-induced phosphorylation of STAT1 and STAT3. Silencing STAT3 also increased rhIL-6-induced STAT1 phosphorylation, but silencing STAT1 had no effect on STAT3 phosphorylation. Pretreatment of myoblasts with neutralising anti-IL-6 antibodies increased phosphorylation of STAT1 and STAT3 induced by 50 ng/mL rhIL-6, whereas pretreatment with 5 ng/mL rhIL-6 reduced this response. Despite increased JAK/STAT signalling, IL-6 silencing decreased glucose and oleic acid uptake and oxidation under both basal and rhIL-6-stimulated conditions. Collectively, our results imply that intrinsic IL-6 restrains activation of the JAK/STAT pathway by extrinsic IL-6, but acts synergistically with it to promote myoblast energy metabolism.

## INTRODUCTION

Skeletal muscle is a major endocrine tissue that secretes a variety of biologically active molecules, collectively known as myokines, which mediate interorgan communication and contribute to the beneficial effects of exercise (Pedersen et al., 2003; Zunner et al., 2022). Among these, interleukin-6 (IL-6) was first characterised as a pro-inflammatory cytokine that stimulates B lymphocytes (Gauldie et al., 1987; Helle et al., 1988; Hirano et al., 1986) and is recognized as an important driver of chronic inflammatory conditions (Garbers et al., 2018; Tanaka et al., 2014), including muscle loss in cancer cachexia (Belizario et al., 2016; Strassmann et al., 1992). In contrast, the exercise-induced increase in serum IL-6 levels, resulting from a transient upregulation of its secretion from contracting skeletal muscles (Bruunsgaard et al., 1997; Ostrowski et al., 1998), has beneficial anti-inflammatory and metabolic effects in various organs and tissues (Ellingsgaard et al., 2011; Wunderlich et al., 2010). Divergent IL-6 actions are likely dependent on the context, such as the site of IL-6 secretion, IL-6 concentrations, the duration and pattern of IL-6 exposure, the type of IL-6 signalling (i.e. classical signalling, trans-signalling, and trans-presentation), and the presence of other cytokines (Lin et al., 2023; Munoz-Canoves et al., 2013; Pelosi et al., 2021; Pelosi et al., 2014; Wei et al., 2026).

As a major source and target of IL-6, skeletal muscle is exposed to dynamic local and systemic fluctuations in IL-6 concentrations, necessitating mechanisms that regulate IL-6 actions (Munoz-Canoves et al., 2013; Pedersen & Febbraio, 2008). IL-6 elicits its effects by binding to its receptor, IL-6Rα, which forms a complex with the signal-transducing subunit IL-6Rβ (also known as gp130 or IL6ST) (Boulanger et al., 2003; Taga et al., 1989). This receptor complex activates the Janus kinases (JAK), which in turn phosphorylate and activate the signal transducer and activator of transcription (STAT) transcription factors (Kang et al., 2020). Physical exercise acutely increases muscle expression not only of IL-6, but also IL-6Rα (Keller et al., 2005), thereby enhancing IL-6 action. Trained individuals, who exhibit a blunted exercise-induced production of IL-6 (Fischer et al., 2004), express higher basal levels of IL-6Rα in skeletal muscle (Keller et al., 2005), indicating that sensitivity to IL-6 is increased and IL-6 actions are maintained despite lower IL-6 secretion. Clearly, regulation of IL-6 action may involve changes in the expression of IL-6 receptors.

Circulating IL-6, which acts in an endocrine manner, contributes to systemic adaptations to exercise (Bruce & Dyck, 2004; Glund et al., 2007; Holmes et al., 2008; Knudsen et al., 2017), and locally produced IL-6, which acts in an autocrine and paracrine manner, promotes skeletal muscle hypertrophy (McKay et al., 2009; Serrano et al., 2008) and regeneration (Becker et al., 2023; Toth et al., 2011). Muscle regeneration involves proliferation and differentiation of myoblasts (Charge & Rudnicki, 2004). The function of these myogenic progenitors is modulated by IL-6 in a concentration dependent manner: low IL-6 levels increase proliferation of myoblasts, whereas higher levels drive their differentiation (Steyn et al., 2019). Additionally, IL-6 stimulates cytoskeletal redistribution in myoblasts, facilitating their migration and fusion into multinucleated myotubes (Liu et al., 2020; Milewska et al., 2020). Although basal secretion of IL-6 is similar between myoblasts and myotubes, myoblasts exhibit greater induction of IL-6 secretion in response to inflammatory stimuli (Frost et al., 2003; Podbregar et al., 2013; Prelovsek et al., 2006). Notably, low concentrations of IL-6 promote IL-6Rα expression in myoblasts, while high concentrations suppress IL-6Rα expression (Steyn et al., 2019), suggesting that the secretory activity of myoblasts might modulate their responsiveness to IL-6.

Skeletal muscles are exposed to both intrinsic (locally produced) IL-6, which acts in an autocrine and paracrine manner, and extrinsic (circulating) IL-6, which acts in an endocrine manner. Local IL-6 production, which may be almost undetectable under resting conditions, can exceed 2 ng/mL during contractions (Rosendal et al., 2005). Circulating IL-6 concentrations fluctuate between 1–100 pg/mL in healthy individuals (Ostrowski et al., 1998), but may exceed 10 ng/mL under septic conditions (Friedland et al., 1992). Given the range of IL-6 concentrations to which skeletal muscle is exposed, intrinsic or extrinsic IL-6 signalling may predominate depending on the context. However, it remains unclear whether and how intrinsic and extrinsic IL-6 interact to modulate IL-6 actions in skeletal muscle.

In this study, we investigated whether intrinsic expression of IL-6 in cultured primary human myoblasts affects their signalling and metabolic responses to treatment with extrinsic (recombinant human) IL-6 (rhIL-6). We found that the reducing intrinsic IL-6 expression increased the responsiveness of the JAK/STAT pathway to extrinsic IL-6, but reduced glucose and oleic acid uptake and oxidation. Collectively, our results imply that intrinsic IL-6 restrains activation of the JAK/STAT pathway by extrinsic IL-6, but acts synergistically with it to promote myoblast energy metabolism.

## MATERIALS AND METHODS

Cell culture flasks and plates were obtained from Sarstedt (Nümbrecht, Germany) and TPP (Trasadingen, Switzerland). CD56 MicroBeads (#130-097-042) were purchased from Miltenyi Biotec (Bergisch Gladbach, Germany). Matrigel Matrix (#356231) and CellBIND® 96-well Clear Flat Bottom Polystyrene Microplates (#3300) were sourced from Corning (Corning, NY, USA). DMEM, low glucose (1 g/L), with GlutaMAX (#21885025), DMEM, high glucose (4.5 g/L), with GlutaMAX (#10566016), Advanced MEM (#12492013), Opti-MEM Reduced Serum Medium (#31985062), Fetal Bovine Serum (#A5256701 and #10500064), DPBS, calcium, magnesium (#14040133), HEPES (1 M, #15630080), Lipofectamine 2000 Transfection Reagent (#11668027), Human EGF Recombinant Protein (#PHG0311), Gentamicin (10 mg/mL, #15710049 and 50 mg/mL, #15750060), Amphotericin B 250 μg/mL (#15290018), Penicillin-Streptomycin 10,000 U/mL (#15140), MEM Vitamin Solution 100x (#11120037), GlutaMAX Supplement 100x (#35050061), 0.25% Trypsin-EDTA (#25200072), Bovine Collagen I (A10644-01), Human IL-6 Recombinant Protein (#200-06), High-Capacity cDNA Reverse Transcription Kit (#4368814), TaqMan Universal Master Mix II, with UNG (#4440038), and Invitrogen Human IL-6 ELISA Kit (#KHC0061) were from Thermo Fisher Scientific (Waltham, MA, USA). Skeletal Muscle Cell Growth Medium (C-23060) was obtained from PromoCell (Heidelberg, Germany). Anti-IL-6 antibody [B-E8] (ab11449) was from Abcam (Cambridge, UK). HumanKine recombinant human LIF protein (#HZ-1292) was from Proteintech (Rosemont, IL, USA). E.Z.N.A. HP Total RNA Kit (#R6812-02) was from Omega Bio-tek (Norcross, GA, USA). FastGene Adhesive PCR Foil (#FG-93AC2) and 96-well PCR plates (FG-03890-50) were from Nippon Genetics (Tokyo, Japan). 4-12% Criterion XT Bis-Tris Precast Gels (#3450123, # 3450124), XT MES Running Buffer (#1610789) and Bio-Rad Protein Assay Dye Reagent Concentrate (#5000006) were from Bio-Rad (Hercules, CA, USA). D-Glucose, [U-14C] (#NEC042V), Oleic Acid, [1-14C] (#NEC317), Ultima Gold XR scintillation liquid (#6013119), UniFilter-96 GF/C Microplates (#6055690), Isoplate 96-well Microplates (#6005040) and TopSeal-A PLUS (#6050185) were from Revvity (Waltham, MA, USA). Amersham ECL Full-Range Rainbow Molecular Weight Marker (#RPN800E) was from Cytiva (Marlborough, MA, USA). Immobilon-P Membrane, PVDF (#IPVH85R), Immobilon Crescendo Western HRP Substrate (#WBLUR0500), Bovine Serum Albumin (BSA, #A3059 and #A6003), Dexamethasone (#D8893), L-carnitine hydrochloride (#C0283), D-Glucose 6-phosphate sodium salt (#G7879), and Sodium oleate (O-7501) were from Merck Millipore (Burlington, MA, USA). ON-TARGETplus Human IL6 siRNA (L-007993-00), ON-TARGETplus Human IL6R siRNA (L-007994-00), ON-TARGETplus Human STAT1 siRNA (L-003543-00), ON-TARGETplus Human STAT3 siRNA (L-003544-00) and ON-TARGETplus Non-targeting Control Pool siRNA (D-001810-10-20) were from Dharmacon Horizon Discovery (Waterbeach, UK). Human IL-6R alpha ELISA Kit (KE00289) was from Proteintech (Rosemont, IL, USA). All other chemicals were from Merck (Darmstadt, Germany).

### Primary human skeletal muscle cell culture

The National Medical Ethics Committee of the Republic of Slovenia granted ethical approval for the preparation and use of primary human skeletal muscle cell cultures (ethical approval numbers 71/05/12 and 0120-698/2017/4). Written informed consent was obtained from all participants. Cell cultures were prepared from *musculus semitendinosus* samples collected as surgical waste during anterior cruciate ligament reconstruction in otherwise healthy individuals. Myogenic cells were isolated as described (Jan et al., 2021; Pirkmajer et al., 2020). First, adipose and connective tissue were manually removed from the samples. The purified muscle tissue was cut into smaller pieces and trypsinised at 37°C for 45 min to release muscle tissue cells, which were then grown in Advanced MEM growth medium supplemented with 10% (v/v) FBS, MEM Vitamin Solution, GlutaMAX, 0.3% (v/v) amphotericin B (250 μg/mL) and 0.15% (v/v) gentamicin (10 mg/mL). After the initial growth phase and before reaching confluence, cells were separated into CD56^+^ and CD56^-^fractions using MACS CD56 MicroBeads (Miltenyi Biotec). CD56, also known as NCAM (neural cell adhesion molecule), is expressed by satellite cells (Cashman et al., 1987), which allows separation of the mixed muscle tissue cell population into predominantly myogenic (CD56^+^) and non-myogenic (CD56^-^) cells (Jan et al., 2021). Cells were resuspended in growth medium with the same composition but supplemented with 10% (v/v) dimethyl sulfoxide (DMSO), and frozen in liquid nitrogen until use. For cell culture experiments, myogenic cells were thawed and cultured in Advanced MEM growth medium in humidified air at 37°C and 5% CO_2_ (v/v) and used for experiments in the proliferative (myoblast) phase at 40-70% confluence.

All experiments and the use of primary skeletal muscle cells for the substrate oxidation assay were approved by the Norwegian Regional Committee for Medical and Health Research Ethics (approval number REK 11959). Skeletal muscle cell cultures were prepared from needle biopsy samples of *musculus vastus lateralis* obtained from healthy individuals (Lund et al., 2018). To isolate satellite cells, biopsies were cut into small pieces and incubated for 30 min in DMEM with 0.1% trypsin-EDTA at room temperature with shaking. The supernatant containing the released satellite cells was collected and supplemented with FBS to a final concentration of 10% (v/v). The trypsinisation step was repeated three times in total. The cell suspension was centrifuged for 7 min at 1800 revolutions per min (RPM) at room temperature. Cells were resuspended in Skeletal Muscle Cell Growth Medium (PromoCell), seeded into a collagen-coated flask, cultured at 37°C and 5% CO_2_, and used for experiments in the proliferative phase.

### Gene silencing

Genes for interleukin-6 (*IL6*), interleukin-6 receptor α (*IL6R*), and two members of the signal transducer and activator of transcription protein family (*STAT1* and *STAT3*) were silenced individually or in combination in cultured human myoblasts. Each gene was silenced by transfection of four pooled gene-specific small interfering RNAs (siRNA), listed in Table 1. Cells were seeded on Matrigel-coated 6-well or 12-well plates and grown in Advanced MEM growth medium until 40-70% confluence. Six hours before siRNA transfection, the growth medium was replaced with Advanced MEM supplemented with 10% (v/v) FBS, MEM Vitamin Solution and GlutaMAX. Pools targeting *IL6* (siIL6), *IL6R* (siIL6R), *STAT1* (siSTAT1), or *STAT3* (siSTAT3) mRNA were diluted in OptiMEM reduced serum medium and transfected into myoblasts with Lipofectamine 2000 Transfection Reagent according to the manufacturer’s protocol at a final concentration of 20 nM siIL6 or 10 nM siIL6R, siSTAT1, and siSTAT3. Control myoblasts were transfected with 20 nM universal, non-targeting scrambled siRNA (siSCR). Twenty-four hours after transfection, the medium was replaced with Advanced MEM growth medium supplemented with 10% (v/v) FBS, MEM Vitamin Solution, GlutaMAX, 0.3% (v/v) Amphotericin B (250 μg/mL), and 0.15% (v/v) Gentamicin (10 mg/mL). Cells were harvested 48 h and 72 h after transfection for mRNA and protein isolation, respectively.

**Table 1:**
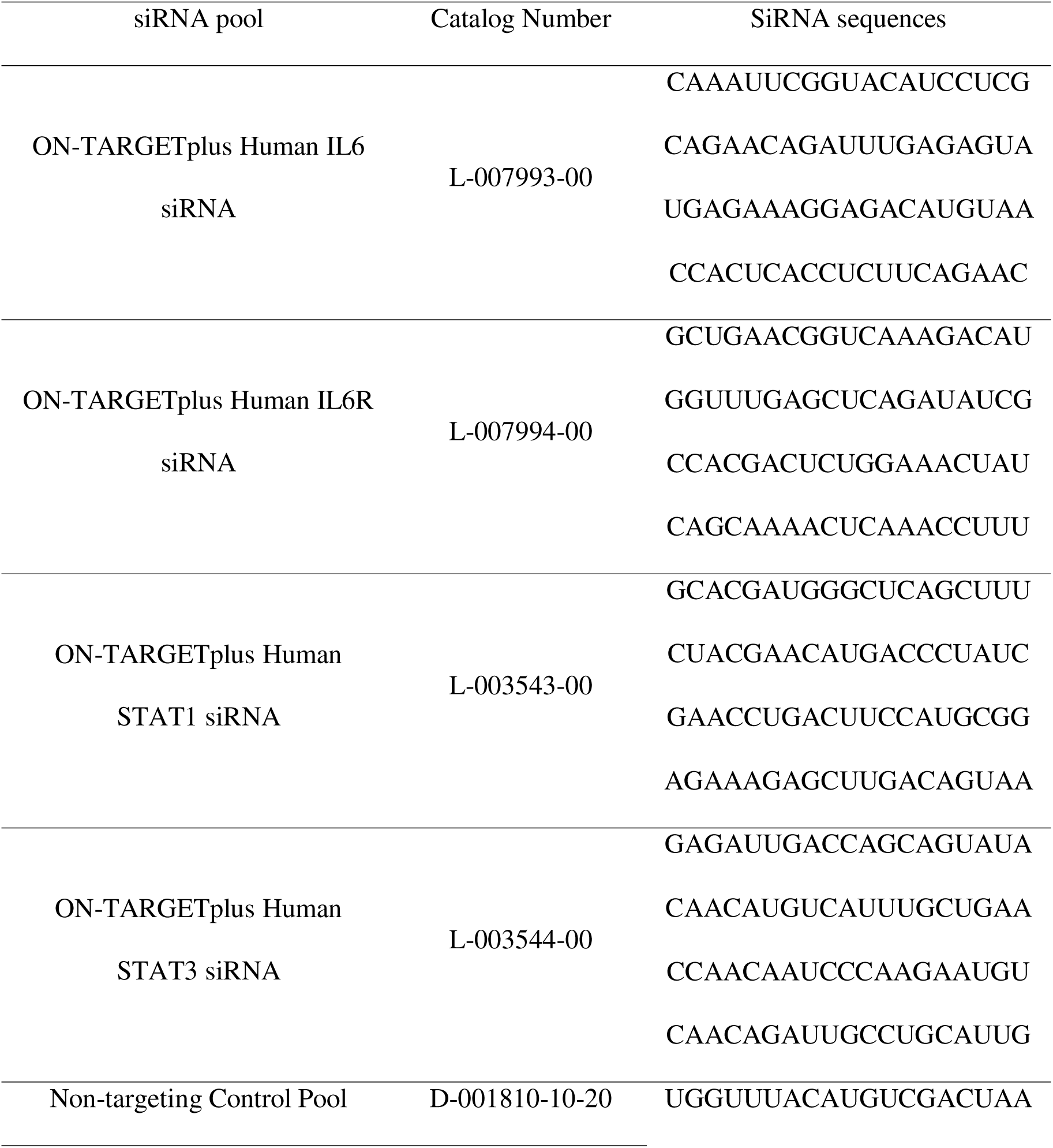
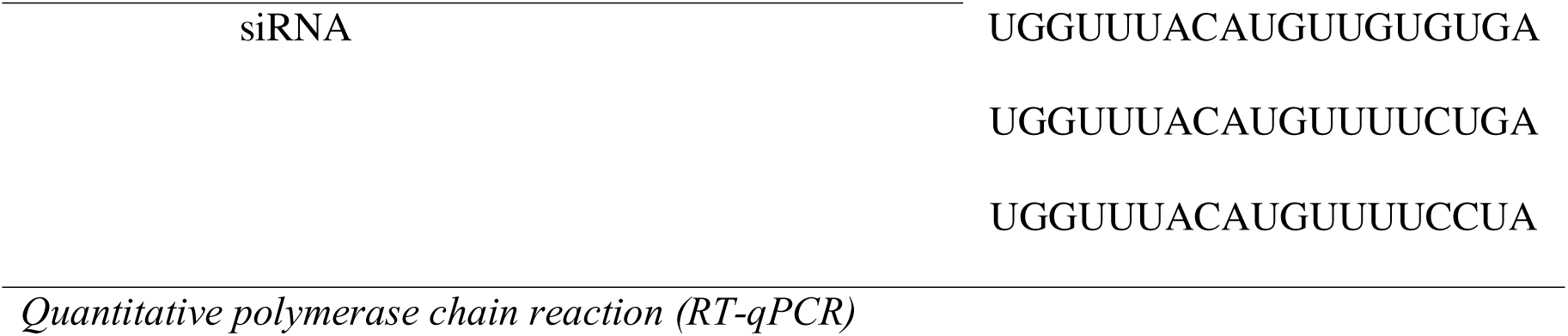
siRNA sequences (Dharmacon Horizon Discovery, Waterbeach, UK).

Forty-eight hours after transfection, cells were washed twice with phosphate-buffered saline (PBS, 137 mM NaCl, 2.7 mM KCl, 10 mM Na_2_HPO_4_, 1.8 mM KH_2_PO_4_, pH 7.4). Total RNA was isolated from the cells with the E.Z.N.A. HP Total RNA Kit. The quantity and quality of the isolated RNA were determined spectrophotometrically using the BioTek Epoch Microplate Spectrophotometer (Agilent Technologies, Santa Clara, CA, USA). mRNA was transcribed into cDNA using the High-Capacity cDNA Reverse Transcription Kit. RT-qPCR was performed with the QuantStudio 3 Real-Time PCR System (Thermo Fisher Scientific, Waltham, MA, USA) using the TaqMan Universal Master Mix II with uracil-N-glycosylase (UNG) and TaqMan Gene Expression Assays listed in Table 2. The expression of individual genes was normalised to the expression of the reference gene 18S rRNA using the following equation:

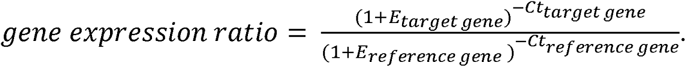

**Table 2:**
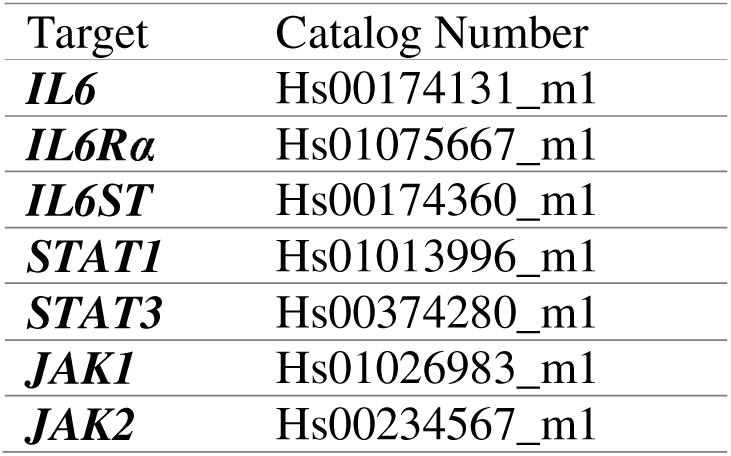
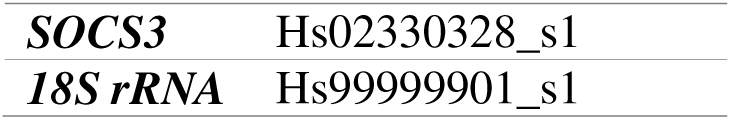
Gene expression assays (Thermo Fisher Scientific, Waltham, MA, USA).

| Target | Catalog Number |
| --- | --- |
| ***IL6*** | Hs00174131_m1 |
| ***IL6Rα*** | Hs01075667_m1 |
| ***IL6ST*** | Hs00174360_m1 |
| ***STAT1*** | Hs01013996_m1 |
| ***STAT3*** | Hs00374280_m1 |
| ***JAK1*** | Hs01026983_m1 |
| ***JAK2*** | Hs00234567_m1 |
| **SOCS3** | Hs02330328_s1 |
| **18S rRNA** | Hs99999901_s1 |

*E* is the PCR efficiency (with values between 0 and 1), calculated for each gene using the LinRegPCR program (Ramakers et al., 2003; Ruijter et al., 2009).

### Treatments with rhIL-6 and rhLIF

Four hours before stimulation of cells with rhIL-6 or recombinant human leukaemia inhibitory factor (rhLIF), the growth medium was replaced with Advanced MEM without FBS to initiate serum starvation. Serum starvation was used to synchronise the cells and reduce interfering effects of FBS (Pirkmajer & Chibalin, 2011). After removing the growth medium supplemented with FBS, the cells were washed twice with PBS and cultured in Advanced MEM without supplements until stimulation. During the last 30 or 15 min of serum starvation, cells were exposed to recombinant human IL-6 (rhIL-6) or recombinant human LIF (rhLIF), respectively. Both proteins were reconstituted in 0.1% (w/v) bovine serum albumin (BSA) in PBS. The final rhIL-6 concentration in the growth medium was 50 ng/mL unless otherwise specified, and the final rhLIF concentration was 10 ng/mL. An equivalent volume of vehicle (0.1% (w/v) BSA in PBS) was added to the control cells.

For the time course experiments, the medium was replaced with fresh Advanced MEM without supplements 3 h after the onset of serum starvation, followed by treatment with rhIL-6 at the final concentration of 50 ng/mL for 5, 10, 15, 30 or 60 min. The concentration-response experiments were performed by exposing myoblasts to 3, 10, 30, 100 or 300 ng/mL rhIL-6 during the last 12.5 min of serum starvation. IL-6 secreted by myoblasts was blocked by culturing them in the presence of a neutralising anti-IL-6 antibody, which was added to the growth medium 24 h after gene silencing and removed before the stimulation with rhIL-6. In some experiments, myoblasts were exposed to a low concentration of rhIL-6 for 24 h before stimulation with 50 ng/mL. In such experiments, myoblasts were washed twice with PBS and then pretreated with 5 ng/mL rhIL-6 or vehicle (0.1% (w/v) BSA/PBS) in serum-free Advanced MEM for 24 h. Myoblasts were ultimately treated with 50 ng/mL rhIL-6 or vehicle during the last 30 min. Stimulation with rhIL-6 or rhLIF was ended by removing the medium, washing the cells three times with ice-cold PBS and freezing the cells.

### Immunoblotting

Cells were lysed and proteins extracted in Laemmli buffer (62.5 mM Tris-HCl (pH 6.8), 10% (w/v) glycerol, 2% (w/v) sodium dodecyl sulfate (SDS), 5% (v/v) 2-mercaptoethanol, 0.002% (w/v) bromophenol blue), sonicated and denatured by heating the lysate at 56 °C for 20 min. Cell lysate proteins were resolved by sodium dodecyl sulfate-polyacrylamide gel electrophoresis (SDS-PAGE) at a constant voltage of 200 V in XT MES Running Buffer using Criterion XT 4-12% Bis-Tris Precast Gels and transferred to a polyvinylidene fluoride (PVDF) membrane in transfer buffer (31 mM Tris, 0.24 M glycine, 10% (v/v) methanol, and 0.01% (w/v) SDS) at a constant voltage of 100 V. Amersham ECL Full-Range Rainbow Molecular Weight Marker was used to estimate the molecular weight of resolved proteins. Sample loading and transfer efficiency were assessed by Ponceau S (0.1% (w/v) in 5% (v/v) acetic acid) staining. The membranes were blocked in 7.5% (w/v) skimmed milk in Tris-buffered saline with Tween 20 (TBST, 20 mM Tris, 150 mM NaCl, 0.02% (v/v) Tween 20, pH 7.5), followed by overnight incubation at 4°C with primary antibodies diluted in primary antibody buffer (20 mM Tris, 150 mM NaCl, 0.1% (w/v) BSA, pH 7.5 and 0.1% (w/v) NaN_3_). The next day, a 1-h incubation with horseradish peroxidase (HRP)-conjugated secondary antibodies diluted in 5% (w/v) skimmed milk in TBST followed. ECL Crescendo Western HRP substrate was added to the membranes, and the immunolabelled proteins were detected with Fusion FX (Vilber, Marne-la-Vallée, France) and quantified using Quantity One 1-D Analysis Software (Bio-Rad, Hercules, CA, USA). All antibodies used for immunoblotting and their dilutions are listed in Table 3.

**Table 3:**
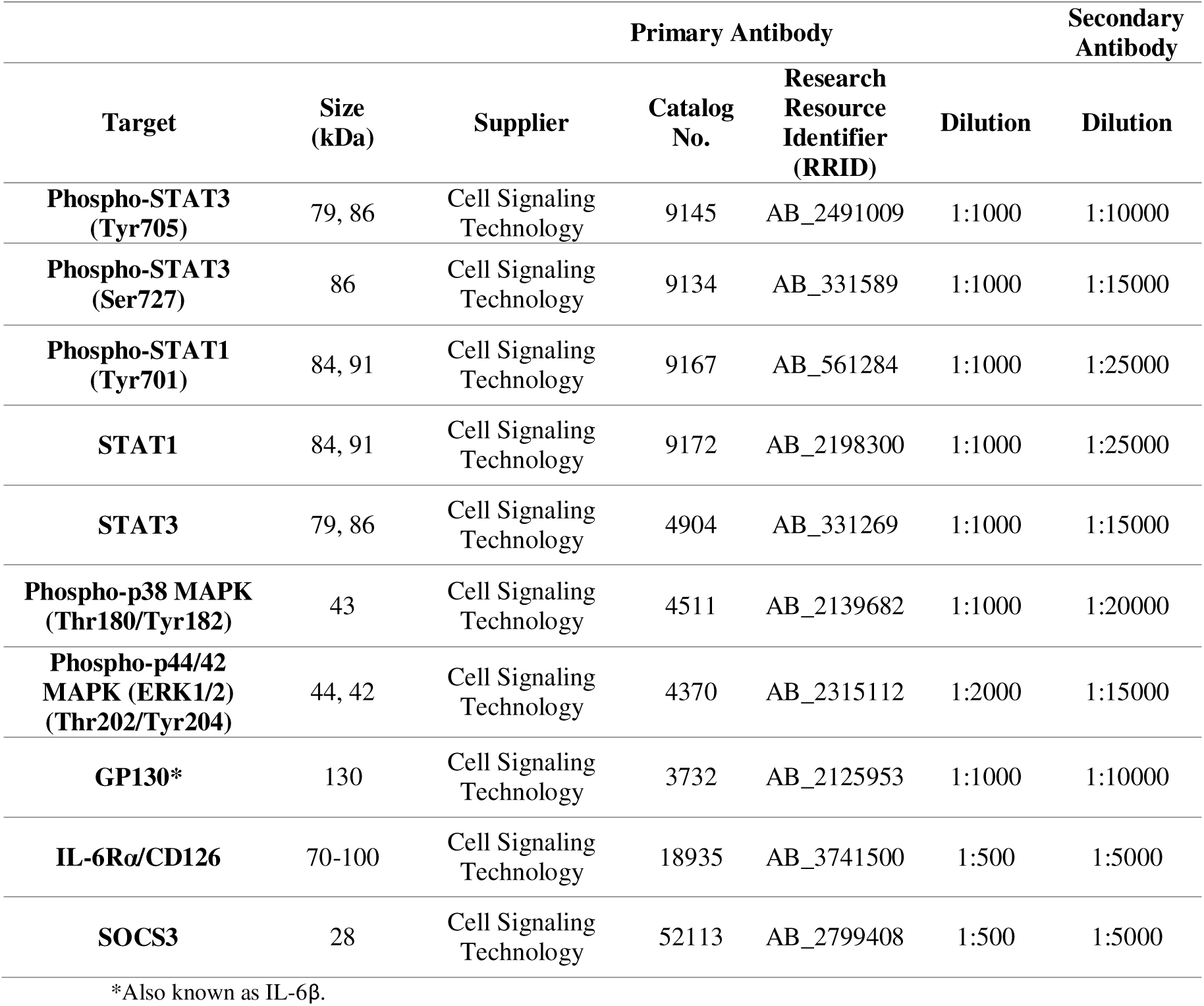
List of antibodies used for immunoblotting. The host species for all primary antibodies was rabbit. The secondary antibody was Goat Anti-Rabbit IgG (H + L)-HRP Conjugate (Bio-Rad #1706515).

### Measurement of IL-6 and sIL-6Rα with ELISA

On the third day after siRNA transfection, 44 h after the final medium exchange, the conditioned medium was collected from siSCR, siIL6, siIL6R and siIL6+siIL6R myoblasts to determine the concentrations of IL-6 and soluble interleukin-6 receptor (sIL-6Rα) secreted by the cells. IL-6 was quantified using the Invitrogen Human IL-6 ELISA (Thermo Fisher Scientific). sIL-6Rα was quantified using the Human IL-6R alpha ELISA Kit (Proteintech).

### Substrate uptake and oxidation assay

Myoblasts were cultured in DMEM-GlutaMAX (1 g/L glucose) supplemented with FBS (10%), streptomycin (25 µg/mL), penicillin (25 IU), gentamicin (50 ng/mL), HEPES (25 mM), BSA (0.05%), dexamethasone (0.39 µg/mL) and human epidermal growth factor (hEGF, 10 µg/mL). Reverse transfection was performed in 96-well CellBIND microplates using 20 nM siRNA targeting IL-6 or a non-targeting scrambled siRNA as described above. Silencing was followed with either 4- or 24-h treatment with rhIL-6 at two concentrations: 1 ng/mL (low) or 10 ng/mL (high). Control cells received vehicle (0.1% BSA in PBS) at a volume equivalent to that of the high rhIL-6 treatment. Metabolic activity of myoblasts was assessed using either ^14^C-labelled oleic acid or ^14^C-labelled glucose as previously described (Wensaas et al., 2007). The cell culture medium was replaced with Dulbecco’s Phosphate-Buffered Saline (DPBS (with Mg^2+^ and Ca^2+^)) containing 10 mM HEPES and 2.4 mM BSA. To assess glucose metabolism, DPBS was supplemented with 200 μM [^14^C]glucose (0.5 μCi/nmol). For analysis of fatty acid metabolism, the medium was supplemented with 100 μM [^14^C]oleic acid (0.5 μCi/nmol) and 1 mM L-carnitine. UniFilter microplates were presoaked with 1 M NaOH and placed on top of cell culture plates. A silicon gasket was placed between them, and a metal clasp was used to ensure airtight seal. After a 4-h incubation at 37°C, cells were washed twice with PBS and lysed in 0.1% NaOH. Captured ^14^CO_2_ in the filter plates and cell-associated radioactivity in the cell lysate were quantified using MicroBeta^2^ liquid scintillation microplate counter (Perkin Elmer, Massachusetts, USA) with Ultima Gold XR as scintillation liquid. Data were normalised to total protein concentration, measured in the cell lysate using Bio-Rad Protein Assay Dye Reagent Concentrate and Victor X4 Multilabel Microplate Reader (Perkin Elmer, Massachusetts, USA). Outliers, identified as values exceeding ± 2 standard deviations from the mean, were excluded from the analysis. Substrate uptake was calculated as the sum of complete oxidation (^14^CO_2_ production) and cell-associated radioactivity, while fractional oxidation was calculated as the ratio of ^14^CO_2_ to total substrate uptake.

### Statistical analysis

Statistical analysis and data visualisation were performed using GraphPad Prism 8 software. Data normality was assessed with the Shapiro-Wilk test. Results are presented as mean and standard deviation (SD) for normally distributed data, or as median, interquartile range, and minimum to maximum range for skewed data. Normally distributed data were analysed using repeated measures one-way ANOVA with Tukey’s *post hoc* multiple comparison test. Skewed data were analysed using the Friedman test and Dunn’s multiple comparison test. Time-course and concentration-response data were analysed using two-way ANOVA with multiple t-tests and Tukey’s multiple comparison test. Mixed-effects analysis and Dunnett’s *post hoc* test were used for substrate oxidation data. Statistical significance was defined as a p-value less than 0.05.

## RESULTS

### Gene silencing of IL-6 enhances rhIL-6-induced STAT1^Tyr701^ and STAT3^Tyr705^ phosphorylation

To determine whether and how intrinsic expression of IL-6 in human myoblasts affects activation of the JAK/STAT pathway by extrinsic (recombinant) human IL-6 (rhIL-6), IL-6 and/or its receptor (IL-6Rα) were knocked-down using siRNA. IL-6 mRNA levels were reduced by 44–47% (p<0.001) in siIL6 myoblasts (Fig. 1a), which was paralleled by a 49–59% reduction in concentrations of secreted IL-6 (p<0.0001) (Fig. 1b). IL-6Rα mRNA levels were reduced by 60–70% (p<0.05) in siIL6R and siIL6+siIL6R myoblasts (Fig. 1c). Interestingly, IL-6 mRNA levels were also reduced by approximately 35% (p=0.0006) in siIL6R myoblasts (Fig. 1a), while IL-6Rα mRNA levels showed a statistically non-significant trend towards upregulation in siIL6 myoblasts (Fig. 1c). Upregulation of IL-6Rα mRNA was significantly different between the siSCR and siIL6 group if analysed separately (p=0.03, Wilcoxon test). The protein abundance of IL-6Rα could not be reliably assessed by immunoblot (data not shown), but ELISA showed that gene silencing of IL-6Rα reduced the concentration of its cleaved (soluble) form (sIL-6Rα) in cell medium below the detection limit (Fig. 1d).

**Figure 1:**
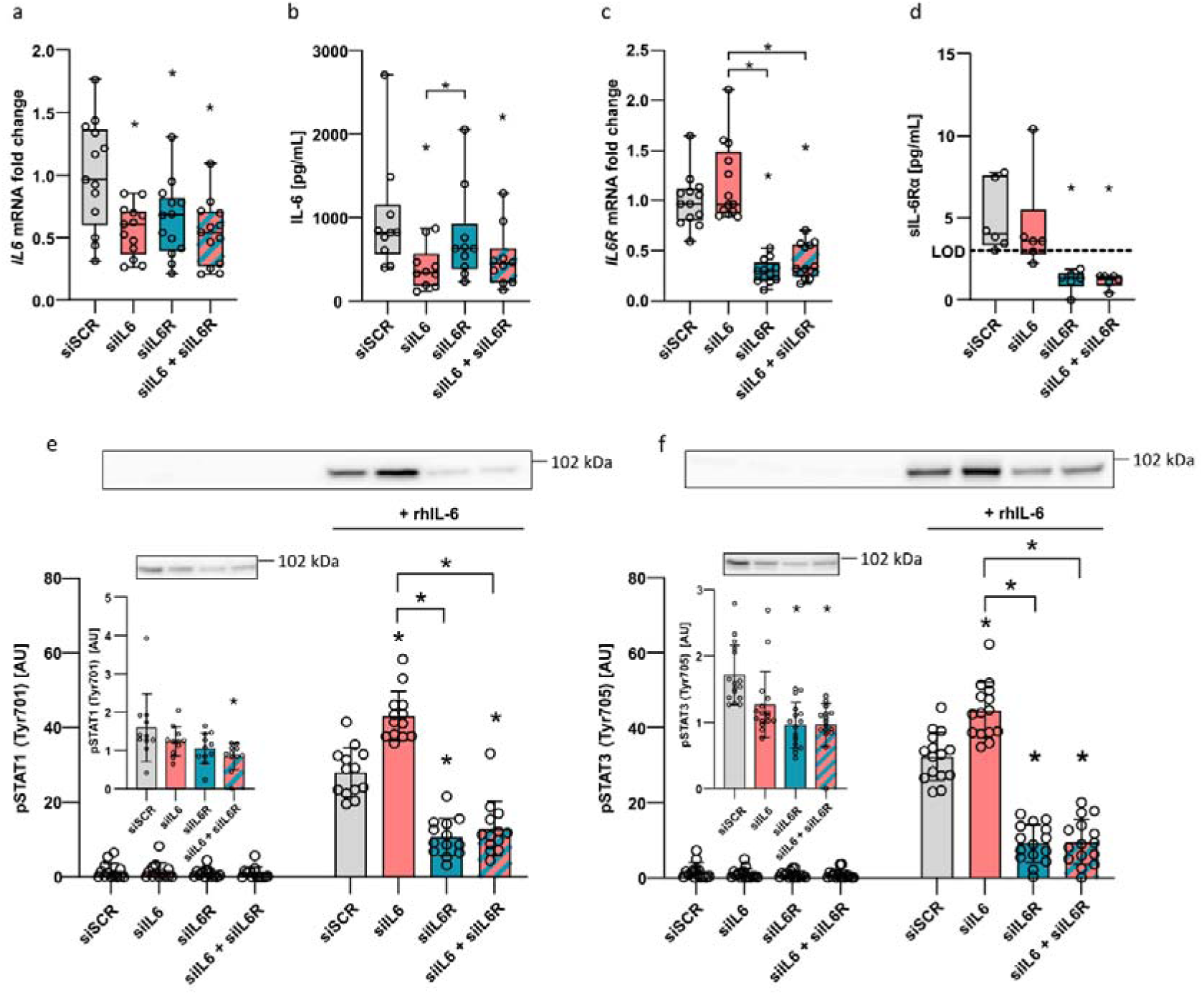
Gene silencing of IL6 and IL6R. Gene expression of IL6 (Fig. 1a) and IL6R (Fig. 1c) in human primary myoblasts after silencing of IL6 (siIL6), IL6R (siIL6R) or the combination of both (siIL6+siIL6R). Negative control cells were transfected by a non-targeting scrambled siRNA (siSCR). IL-6 (Fig. 1b) and soluble IL-6Rα (Fig. 1d) concentration in the conditioned media collected from siSCR, siIL6, siIL6R and siIL6+siIL6R myoblasts on the third day after gene silencing. Fig. 1e, f: pSTAT1 (Tyr701) and pSTAT3 (Tyr705) levels after silencing of either or both IL6 and IL6R in basal conditions (left panel) and after a 30-min exposure to 50 ng/mL rhIL-6 (right panel). Inset graphs represent separately analysed unstimulated pSTAT1/3 levels. qPCR results are normalised to the average siSCR value. qPCR and ELISA results are presented as median, interquartile range, and minimum to maximum range. Immunoblotting results are presented as mean and SD. N=6-16, 3-4 independent experiments. (*)p<0.05, compared to siSCR within the same condition (basal or rhIL-6-stimulated), or as indicated by the brackets. AU, arbitrary units. LOD, limit of detection.

The activity of the JAK/STAT signalling pathway was assessed by measuring phosphorylation of STAT1 at tyrosine 701 (pSTAT1^Tyr701^) and STAT3 at tyrosine 705 (pSTAT3^Tyr705^). In the absence of rhIL-6, phosphorylation of STAT1^Tyr701^ (Fig. 1e, left panel) and STAT3^Tyr705^ (Fig. 1f, left panel) were reduced in siIL6R and siIL6+siIL6R myoblasts. In the presence of 50 ng/mL rhIL-6 (30 min), phosphorylation of STAT1^Tyr701^ was 53% higher (p=0.0007, Fig. 1e, right panel) and STAT3^Tyr705^ 38% higher (p<0.0001, Fig. 1f, right panel) in siIL6 than control (siSCR) myoblasts. Phosphorylation of STAT1^Tyr701^ and STAT3^Tyr705^ was increased by rhIL-6 also in siIL-6R and siIL6+siIL6R but was markedly suppressed compared to siSCR or siIL6 myoblasts (Fig. 1 e, f).

### Gene silencing of IL-6 enhances rhIL-6-induced STAT3^Tyr705^ phosphorylation in a time-dependent manner

We treated siSCR and siIL6 myoblasts with 50 ng/mL rhIL-6 for 5–60 min to determine whether gene silencing of IL-6 affects the kinetics of STAT3^Tyr705^ phosphorylation during exposure to rhIL-6 (Fig. 2). The rhIL-6-induced increase in phosphorylation of STAT3^Tyr705^ was significantly higher in siIL6 than siSCR myoblasts at 10, 15, and 30 min (p<0.0001, Fig. 2a). The area under the curve (AUC) was 1.5-fold larger in siIL6 than in siSCR myoblasts (p<0.0001, Fig. 2b). Conversely, phosphorylation of STAT3 at Ser727 (Fig. 2c), which is needed for full activation of STAT3 (Wen et al., 1995), and total STAT3 levels (Fig. 2d) were not significantly altered by any of the treatments. As cytokines activate mitogen-activated protein kinases (MAPK) in some cell types (Daeipour et al., 1993; Fahmi et al., 2013; Ishikawa et al., 2003; Raingeaud et al., 1995), we also measured phosphorylation of ERK1/2^Thr202/Tyr204^ (Fig. 2e) and p38-MAPK^Thr180/Tyr182^ (Fig. 2f), whose levels remained unaltered after IL-6 silencing or rhIL-6 treatment.

**Figure 2:**
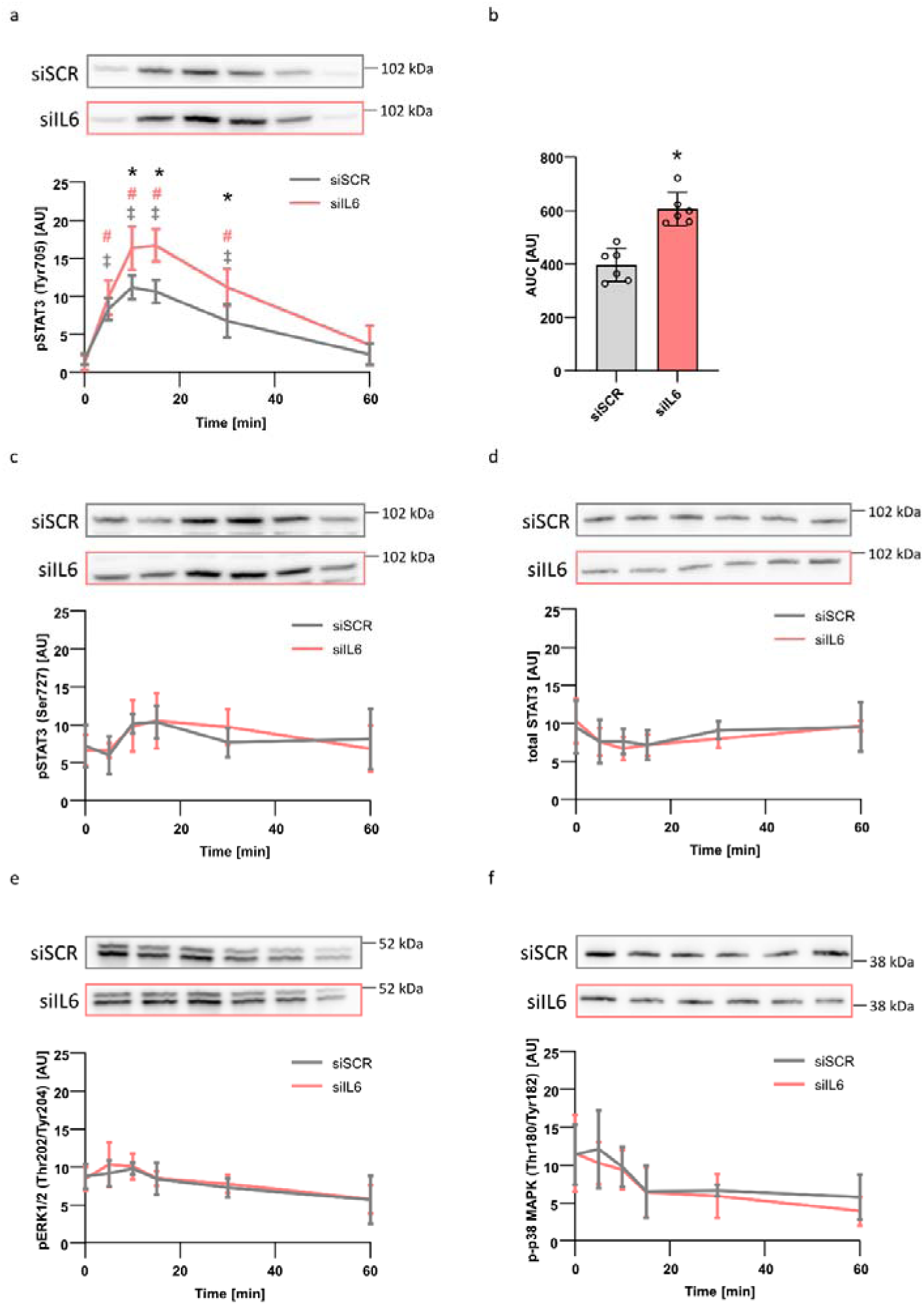
Time-course of rhIL-6 stimulation. siSCR and siIL6 myoblasts were exposed to 50 ng/mL rhIL-6 for 5, 10, 15, 30 or 60 min. Control cells (0) were exposed to an equivalent volume of solvent (0.1% BSA/PBS) for 60 min. pSTAT3 (Tyr705) levels were determined by immunoblotting (Fig. 2a). Area under the curve (AUC) was calculated for each curve and compared between siSCR and siIL6 myoblasts (Fig. 2b). pSTAT3 (Ser727) (Fig. 2c), total STAT3 (Fig. 2d), pERK1/2 (Thr202/Tyr204) (Fig. 2e) and p-p38 MAPK (Thr180/Tyr182) (Fig. 2f) levels were determined by immunoblotting. The results are presented as mean and SD. N=5, two independent experiments. (*)p<0.05 siIL6 vs siSCR. (‡)p<0.05 siSCR vs basal (0) siSCR. (#)p<0.05 siIL6 vs basal (0) siIL6. AU, arbitrary units.

### Gene silencing of IL-6 enhances rhIL-6-induced STAT3^Tyr705^ phosphorylation in a concentration-dependent manner

We treated siSCR and siIL6 myoblasts with 3–300 ng/mL rhIL-6 for 12.5 min (Fig. 3), which was the approximate time of maximal phosphorylation of STAT3 in the time course experiments. Phosphorylation of STAT3^Tyr705^ was higher in siIL6 than in siSCR myoblasts (p<0.0001, Fig. 3a) after exposure to rhIL-6 concentrations of 30 ng/mL or higher. The extrapolated maximal effect (E_max_), which reflects IL-6 responsiveness, was ∼1.5-fold higher in siIL6 myoblasts (p=0.02, Fig. 3b). The concentration at half-maximal response (EC_50_), which reflects IL-6 sensitivity, was 14.7 ± 2 ng/mL in siSCR myoblasts and 20.3 ± 7 ng/mL in siIL6 myoblasts, but the difference did not reach statistical significance (Fig. 3c). The levels of pSTAT3^Ser727^ (Fig. 3d), total STAT3 (Fig. 3e), pERK1/2^Thr202/Tyr204^ (Fig. 3f), and p-p38 MAPK^Thr180/Tyr182^ (Fig. 3g) remained similar across treatments.

**Figure 3:**
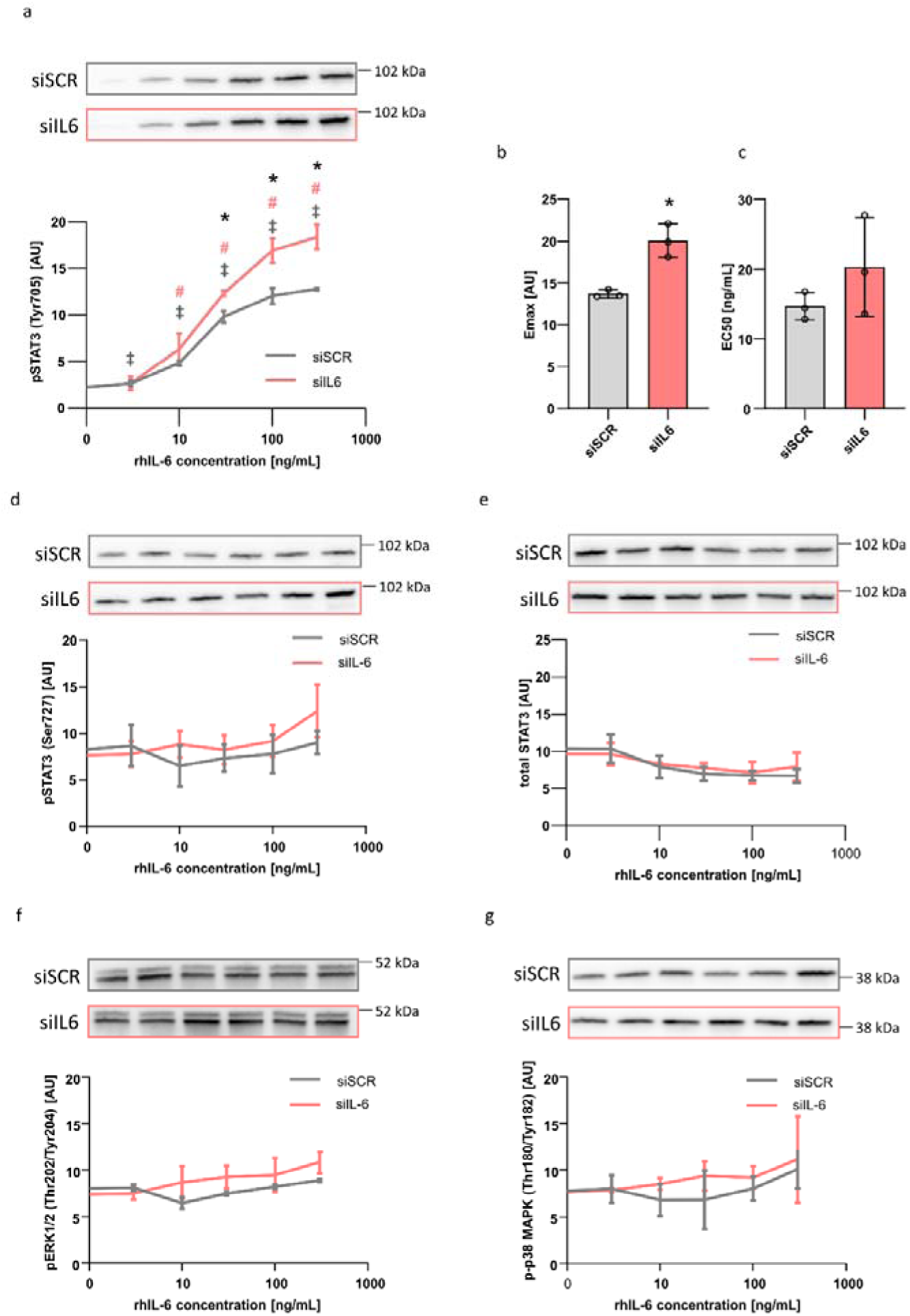
The response of siSCR and siIL6 myoblasts to different concentrations of rhIL-6. Cells were exposed to 3, 10, 30, 100 and 300 ng/mL rhIL-6 for 12.5 min. Control cells (0) were exposed to an equivalent volume of vehicle (0.1% BSA/PBS). pSTAT3 (Tyr705) levels were determined by immunoblotting (Fig. 3a). The maximal response (Emax, Fig. 3b) and the concentration at half-maximal response (EC50, Fig. 3c) were determined from the individual concentration-response curves. pSTAT3 (Ser727) (Fig. 3d), total STAT3 (Fig. 3e), pERK1/2 (Thr202/Tyr204) (Fig. 3f) and p-p38 MAPK (Thr180/Tyr182) (Fig. 3g) levels were determined by immunoblotting. The results are presented as mean and SD. N=3, one experiment. (*)p<0.05 siIL6 vs siSCR. (‡)p<0.05 siSCR vs basal (0) siSCR. (#)p<0.05 siIL6 vs basal (0) siIL6. AU, arbitrary units.

### Gene silencing of STAT3 enhances rhIL-6-induced STAT1^Tyr701^ phosphorylation

As gene silencing of IL-6 enhanced rhIL-6-induced activation of the JAK/STAT pathway, we next determined whether partial suppression of the JAK/STAT pathway has a similar effect. We silenced STAT1 and/or STAT3 (Fig. 4a, b) in myoblasts and then stimulated them with 50 ng/mL rhIL-6 for 30 min. Gene silencing of STAT1 had no effect on STAT3 levels or its phosphorylation (Fig. 4b, d). Conversely, gene silencing of STAT3 increased STAT1 levels (p<0.0001) in unstimulated myoblasts (Fig. 4a) and enhanced rhIL-6-induced phosphorylation of STAT1^Tyr701^ (p=0.0006, Fig. 4c). Taken together, our results indicated that rhIL-6-induced STAT1 phosphorylation can be increased both by gene silencing of IL-6 or STAT3.

**Figure 4:**
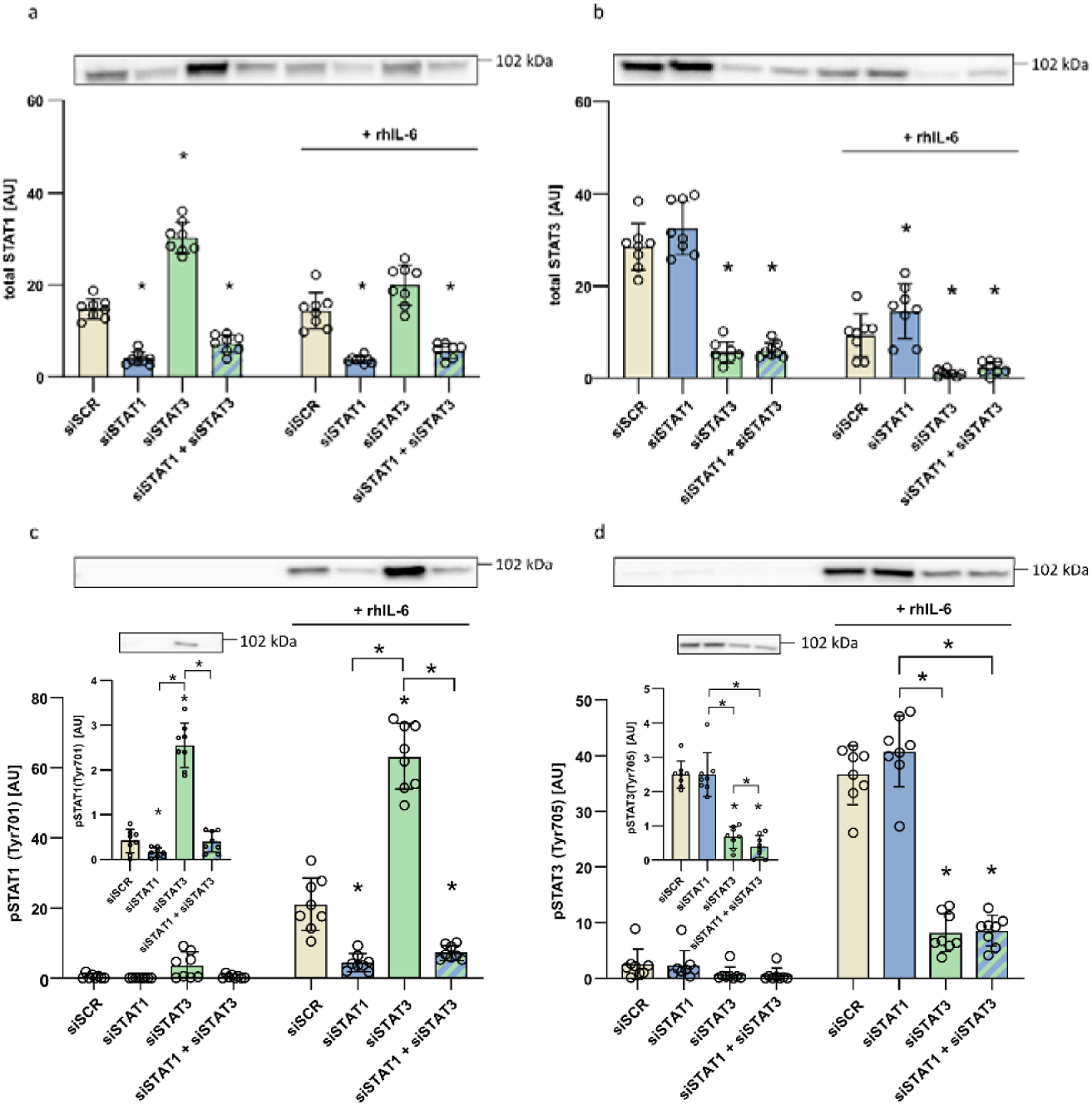
Total STAT1 (Fig. 4a) and STAT3 protein levels (Fig. 4b) after silencing in basal conditions (left panel) and after a 30 min exposure to 50 ng/mL rhIL-6 (right panel). Protein abundance of pSTAT1 (Tyr701) (Fig. 4c), and pSTAT3 (Tyr705) (Fig. 4d), basal (left panel) or stimulated by 50 ng/mL rhIL-6 for 30 min (right panel), analysed by immunoblotting 72 h after siRNA transfection. Inset graphs represent separately analysed unstimulated pSTAT1/3 levels. Results are presented as mean with SD. N=8-12, 4 independent experiments. (*)p<0.05 compared to siSCR within the same condition (basal or rhIL-6-stimulated), or as indicated by the brackets. AU, arbitrary units.

### Effect of IL-6 silencing on the expression of the components of the IL-6R/JAK/STAT pathway

Gene silencing of STAT3 increased the abundance of STAT1 and enhanced rhIL-6-induced STAT1^Tyr701^ phosphorylation. We therefore examined whether altered expression of the key components of the IL-6R/JAK/STAT pathway could provide a mechanism by which gene silencing of IL-6 enhanced rhIL-6 action. The mRNA and protein levels of IL-6Rβ (also known as IL6ST and gp130) (Fig. 5a, b) were unaltered by gene silencing of IL-6 and IL6-Rα. The expression levels of *JAK1* (Fig. 5c), *JAK2* (Fig. 5d), *STAT1* (Fig. 5e), and *STAT3* (Fig. 5f) were unaltered by gene silencing of IL-6. Expression of suppressor of cytokine signalling 3 (*SOCS3*), a negative regulator of the JAK/STAT pathway (Croker et al., 2003; Starr et al., 1997), was also unaltered (Fig. 5g). These results did not support the notion that gene silencing of IL-6 enhanced rhIL-6 action by altering the expression of key components of the IL-6R/JAK/STAT pathway.

**Figure 5:**
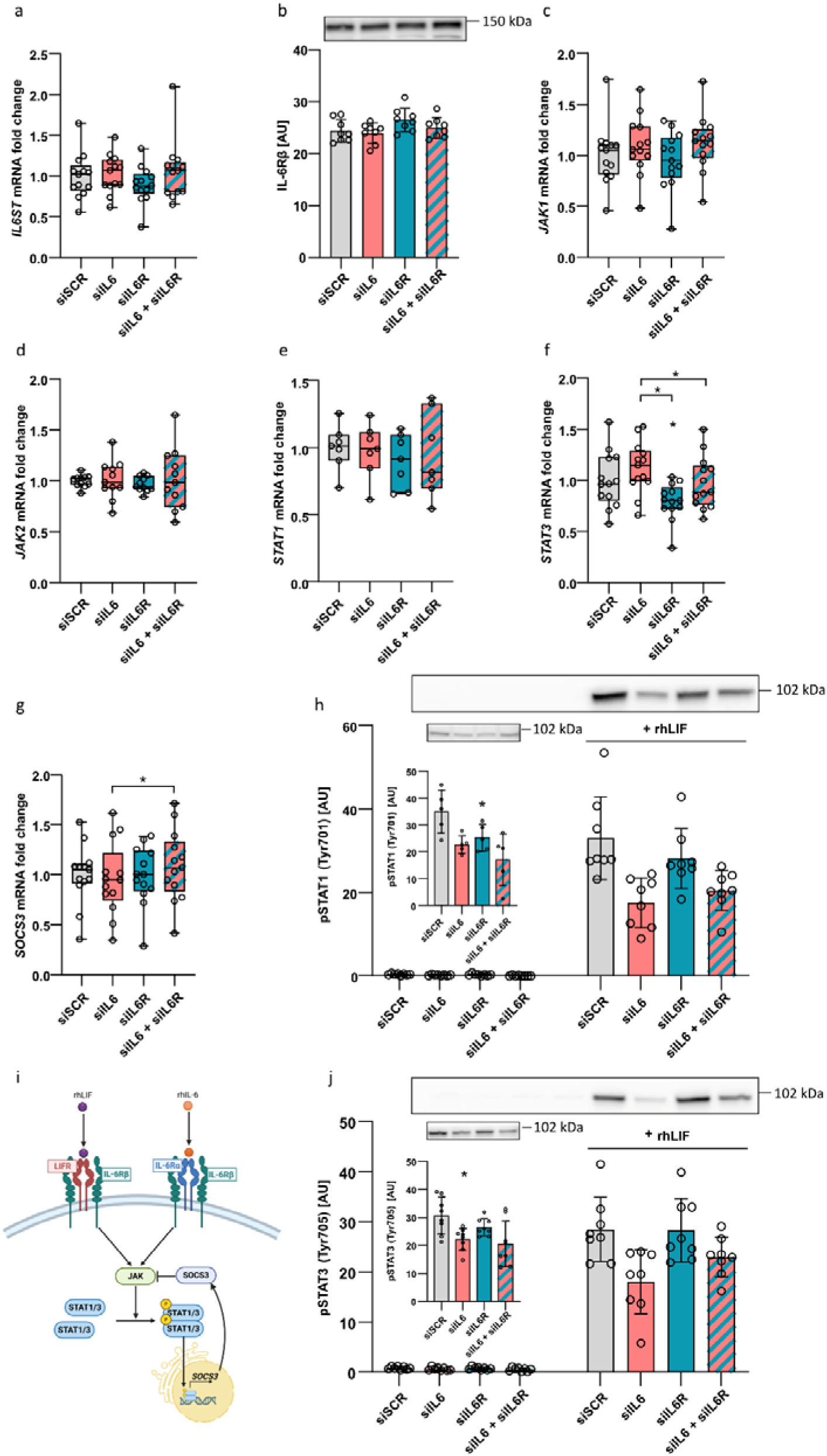
Gene expression of IL6ST (Fig. 5a), analysed by qPCR 48 h after siRNA transfection, and protein levels of IL6ST/IL-6Rβ (Fig. 5b), analysed by immunoblotting 72 h after siRNA transfection. Gene expression of JAK1 (Fig. 5c), JAK2 (Fig. 5d), STAT1 (Fig. 5e), STAT3 (Fig. 5f) and SOCS3 (Fig. 5g) in siIL6, siIL6R and siIL6+siIL6R myoblasts, analysed by qPCR 48 h after siRNA transfection. Fig. 5h, j: pSTAT1 (Tyr701) and pSTAT3 (Tyr705) levels after silencing of either or both IL6 and IL6R in basal conditions (left panel) and after a 15-min exposure to 10 ng/mL rhLIF (right panel). Inset graphs represent separately analysed unstimulated pSTAT1/3 levels. Fig. 5i: Schematic presentation of IL-6 and LIF signalling (rhLIF, recombinant human leukaemia inhibitory factor; rhIL-6, recombinant human IL-6; LIFR, LIF receptor; IL-6Rα, IL-6 receptor subunit α; IL-6Rβ, IL-6 receptor subunit β; JAK, Janus kinase; STAT1/3, signal transducer and activator of transcription 1/3; SOCS3, suppressor of cytokine signalling 3). Fig. 5i was created with BioRender. qPCR results are normalised to the average siSCR value. qPCR results are presented as median, interquartile range, and minimum to maximum range. Immunoblotting results are presented as mean and SD. N=7–13, 2–4 independent experiments. (*)p<0.05 compared to siSCR within the same condition (basal or rhIL-6-stimulated), or as indicated by the brackets. AU, arbitrary units.

### Gene silencing of IL-6 does not enhance rhLIF-induced STAT1^Tyr701^ and STAT3^Tyr705^ phosphorylation

Leukaemia inhibitory factor (LIF) activates the JAK/STAT pathway via a complex comprising LIF receptor (LIFR) and IL-6Rβ (Bower et al., 1995; Gearing et al., 1991) (Fig. 5i). As LIF partially shares the signalling machinery with IL-6 (Heinrich et al., 2003), we examined whether gene silencing of IL-6 enhances actions of recombinant human LIF (rhLIF). The phosphorylation of STAT1^Tyr701^ (Fig. 5h) and STAT3^Tyr705^ (Fig. 5j) was increased by rhLIF, but in contrast to rhIL-6, rhLIF-induced phosphorylation of STAT1 and STAT3 tended to be the lowest in siIL6 myoblasts. This result indicates that gene silencing of IL-6 does not enhance cytokine-induced activation of the JAK/STAT pathway in general.

### Pretreatment with neutralizing anti-IL-6 antibody enhances rhIL-6-induced STAT1^Tyr701^ and STAT3^Tyr705^ phosphorylation

Based on gene silencing experiments we hypothesized that the presence of endogenous IL-6 that is intrinsically expressed and secreted into the medium might suppress the response to the rhIL-6 treatment. To block IL-6 that was secreted in the medium, we used a commercially available neutralising antibody against IL-6 (NAbIL6). We first tested its efficacy by exposing myoblasts to different NAbIL6 concentrations and 5 ng/mL rhIL-6. The rhIL-6-induced phosphorylation of STAT1^Tyr701^ (Fig. 6a) and STAT3^Tyr705^ (Fig. 6b) was completely blocked by 100 ng/mL NAbIL6, which corresponds to an approximate 2.8-fold molar excess of NAbIL6 to rhIL-6. As mean concentrations of endogenous IL-6 in cell medium reached ∼1 ng/mL in our experiments (Fig. 1b), we decided to use 40 ng/mL NAbIL6 for further treatments.

**Figure 6:**
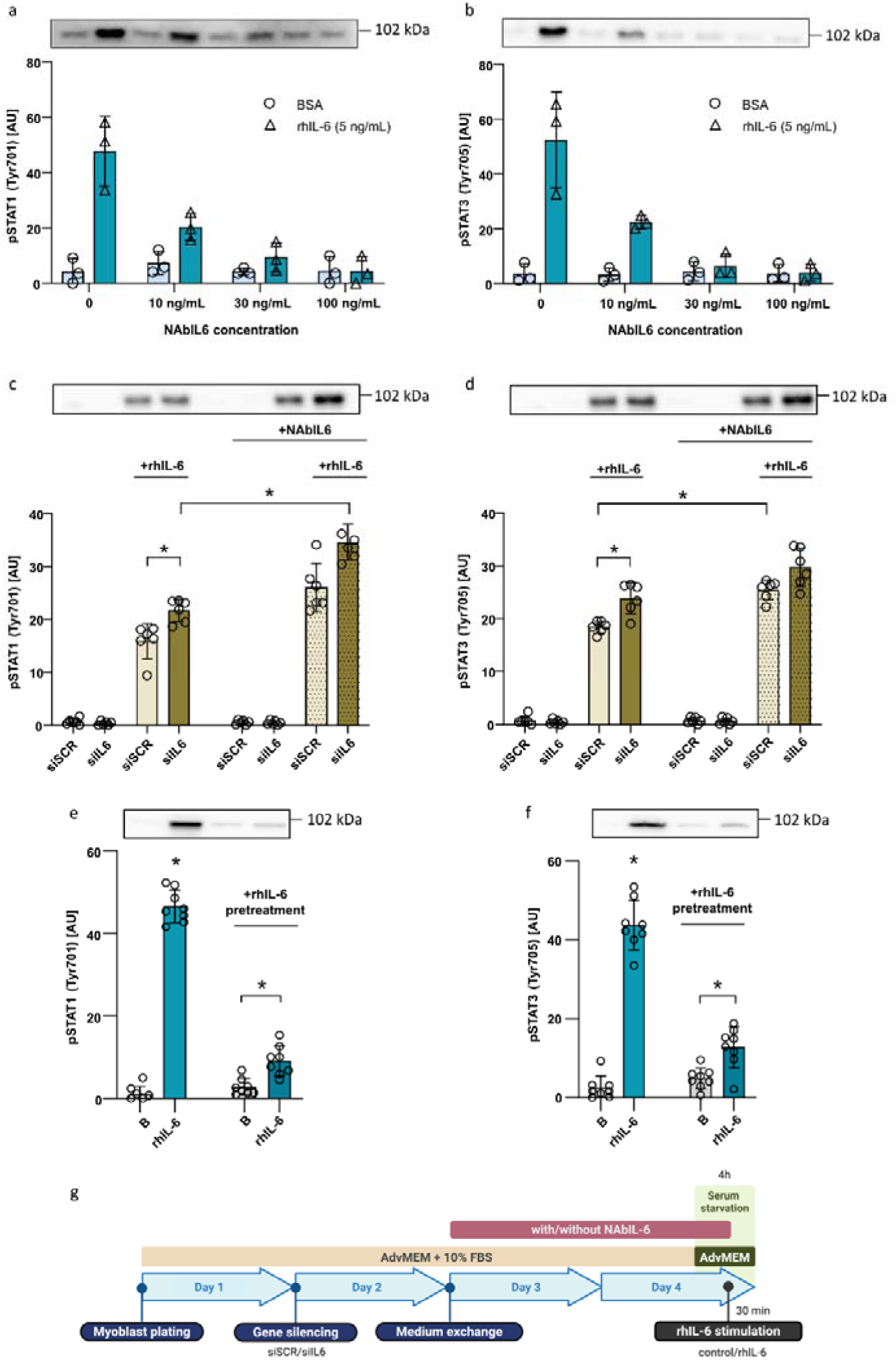
Levels of pSTAT1 (Tyr701) (Fig. 6a) and pSTAT3 (Tyr705) (Fig. 6b) after a 30-min exposure of myoblasts to 50 ng/mL rhIL-6 or 0.1 % BSA/PBS as negative control, mixed with different concentrations of an anti-IL-6 neutralising antibody (NAbIL6). Experimental combinations of IL6 silencing, rhIL-6 stimulation, and/or presence of 40 ng/mL NAbIL6 were performed, and levels of pSTAT1 (Tyr701) (Fig. 6c) and pSTAT3 (Tyr705) (Fig. 6d) were measured by immunoblotting. Levels of pSTAT1 (Tyr701) (Fig. 6e) and pSTAT3 (Tyr705) (Fig. 6f) in myoblasts exposed to 50 ng/mL rhIL-6 for 12.5 min, and pretreated without or with 5 ng/mL rhIL-6. The experimental procedure for Fig. 6c, d is summarized in Fig. 6g (created with BioRender). Data are presented as mean and SD. N=3, one independent experiment (Fig. 6a, b), N=6, 2 independent experiments (Fig. 6c, d). N=8, two independent experiments (Fig. 6e, f). (*)p<0.05 compared to siSCR within the same condition (basal or rhIL-6-stimulated), or as indicated by the brackets. AU, arbitrary units.

To neutralize endogenous IL-6 before the treatment with rhIL-6, siSCR and siIL6 myoblasts were cultured in growth medium with or without NAbIL6. After pretreatment with NAbIL6, myoblasts were switched to a fresh medium without NAbIL6 and treated with 50 ng/mL rhIL-6 for 30 min (Fig. 6g). As assessed by the two-way ANOVA, rhIL-6-induced phosphorylation of STAT1^Tyr701^ (p<0.0001, Fig. 6c) and STAT3^Tyr705^ (p<0.0001, Fig. 6d) were higher overall in the presence of NAbIL6 than in its absence. Analysis of inter-group differences showed that pretreatment with NAbIL6 increased rhIL-6-induced phosphorylation of STAT1^Tyr701^ in siIL6 myoblasts (Fig. 6c) and rhIL-6-induced phosphorylation of STAT3^Tyr705^ in siSCR myoblasts (Fig. 6d). These results suggest that neutralization of endogenous IL-6 with NAbIL6 promotes rhIL-6-induced activation of the JAK/STAT pathway.

### Pretreatment with rhIL-6 blunts rhIL-6-induced STAT3^Tyr705^ and STAT1^Tyr701^ phosphorylation

As pretreatment with NAbIL6 increased the response to 50 ng/mL rhIL-6, we hypothesized that pretreatment with low concentration of rhIL-6 would suppress the activation of the JAK/STAT pathway upon stimulation with high concentration of rhIL-6. To evaluate this hypothesis, myoblasts were pretreated with 5 ng/mL rhIL-6 for 24 h, which was followed by treatment with 50 ng/mL rhIL-6 for 12.5 min (Fig. 6e, f). Pretreatment with 5 ng/mL rhIL-6 did not prevent but markedly blunted the increase in phosphorylation of STAT1^Tyr701^ (Fig. 3e) and STAT3^Tyr705^ (Fig. 3f) after 12.5-min treatment with 50 ng/mL rhIL-6.

### Gene silencing of IL-6 suppresses glucose and lipid metabolism

Control and siIL6 myoblasts were exposed to 1 or 10 ng/mL rhIL-6 for 4 or 24 h, followed by incubation with radioactive-labelled glucose or oleic acid (Fig. 7). In control myoblasts, the 4-h treatment with rhIL-6 increased glucose uptake (p=0.0001, Fig. 7a), complete glucose oxidation (p<0.0001, Fig. 7c), fractional glucose oxidation (p=0.03–0.05, Fig. 7e), complete oleic acid oxidation (p=0.005–0.0001, Fig. 7d), and fractional oleic acid oxidation (p=0.03, Fig. 7f), while the 24-h rhIL-6 treatment increased complete oleic acid oxidation (Fig. 7d) without affecting glucose metabolism. In contrast, gene silencing of IL-6 reduced glucose uptake (p<0.0001, Fig. 7a), complete glucose oxidation (p<0.0001, Fig. 7c), oleic acid uptake (Fig. 7b, p<0.0001) and complete oleic acid oxidation (p<0.0001, Fig. 7d). In siIL6 myoblasts, rhIL-6 increased complete (Fig. 7d) and/or fractional (Fig. 7f) oleic acid oxidation but had no effect on glucose metabolism (Fig. 7a, c, e). In conclusion, rhIL-6 upregulated multiple metabolic processes, whereas IL-6 silencing inhibited these same processes, an effect that rhIL-6 was unable to fully rescue.

**Figure 7:**
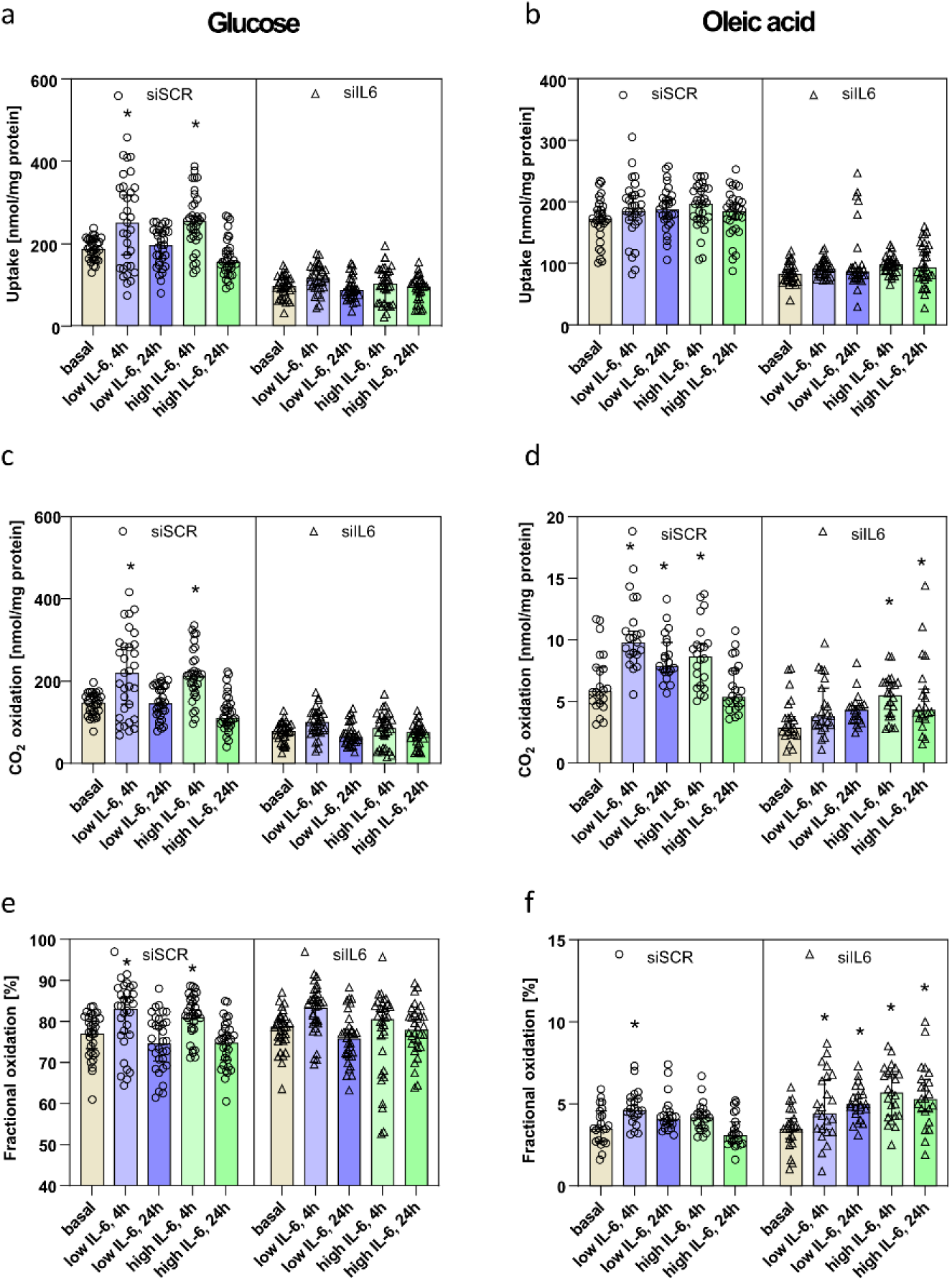
Uptake, complete oxidation and fractional oxidation of glucose (Fig. 7a, c, e) and oleic acid (Fig. 7b, d, f) in control (siSCR) and IL6-silenced (siIL6) myoblasts. Myoblasts were exposed to low (1 ng/mL) or high (10 ng/mL) concentration of rhIL-6 for 4 or 24 h, followed by incubation with radioactive-labelled glucose or oleic acid. Substrate uptake was calculated as the sum of complete oxidation (^14^CO_2_ production) and cell-associated radioactivity, while fractional oxidation was calculated as the ratio of ^14^CO_2_ to total substrate uptake. The effect of silencing was evaluated by a mixed-model analysis, and the effects of treatment were individually compared to the respective basal value using Dunnett‘s test. N=3-4, 2 independent experiments. (*)p<0.05 compared to basal within the same condition (siSCR or siIL6).

## DISCUSSION

In this study, we examined whether intrinsic expression of IL-6 in cultured human myoblasts affected their ability to respond to rhIL-6. We found that gene silencing of IL-6 enhanced rhIL-6-induced, but not rhLIF-induced, phosphorylation of STAT1^Tyr701^ and STAT3^Tyr705^.

Pretreatment with IL-6 neutralizing antibody also enhanced rhIL-6-induced phosphorylation of STAT1^Tyr701^ and STAT3^Tyr705^ in myoblasts, while pretreatment with low concentration of rhIL-6 suppressed it. In contrast to the enhanced signalling responses, gene silencing of IL-6 suppressed basal and rhIL-6-induced glucose and oleic acid metabolism. Interestingly, metabolic effects of rhIL-6, especially on glucose metabolism, were most pronounced during the 4-hour treatment with low rhIL-6 in siSCR myoblasts, while 24-hour treatment with high rhIL-6 significantly increased oleic acid oxidation in siIL6 myoblasts, although it had no effect under control conditions.

Unstimulated myoblasts exhibited relatively low levels of phosphorylated STAT1^Tyr701^and STAT3^Tyr705^ compared to those during treatment with rhIL-6. Nevertheless, gene silencing of IL-6 and/or IL-6Rα reduced their phosphorylation levels, demonstrating that intrinsic expression and secretion of IL-6 lead to activation of the IL-6R/JAK/STAT pathway. This indicates that cultured myoblasts were exposed to continuous autocrine and/or paracrine IL-6 stimulation due to its intrinsic expression. Although this stimulation was relatively modest, as estimated from the STAT1 and STAT3 phosphorylation levels, it appears to be functionally significant in regulating IL-6 responsiveness and energy metabolism in myoblasts. Indeed, both gene silencing of IL-6 and neutralization of intrinsically produced IL-6 with NAbIL6 increased the responsiveness of the JAK/STAT pathway upon rhIL-6 stimulation. These results suggest that myoblast responsiveness to extrinsic IL-6 is inversely proportional to the level of intrinsic IL-6 signalling. If a similar mechanism exists *in vivo*, upregulation of local IL-6 signalling could make myoblasts less receptive for the systemic, circulating/extrinsic IL-6, whereas downregulation of local IL-6 signalling could make them more receptive to the circulating/extrinsic IL-6 stimulation.

Gene silencing of IL-6 suppressed glucose and oleic acid metabolism, demonstrating that intrinsic secretion of IL-6, which leads to continuous low-grade activation of the JAK/STAT pathway in cultured myoblasts, is important for basal energy substrate metabolism of myoblasts. The siIL6 myoblasts were incapable of increasing glucose uptake and oxidation upon stimulation with rhIL-6 and while fatty acid oxidation was increased, the response to rhIL-6 was blunted and oxidation remained below its basal level in control (siSCR) myoblasts. Thus, an increased capability to activate the JAK/STAT pathway in siIL6 myoblasts did not translate into enhanced metabolic responses during stimulation with rhIL-6. The suppressed metabolic responses in siIL6 myoblasts are consistent with observations that IL-6 increases glucose uptake in cultured human (Al-Khalili et al., 2006) and L6 (Carey et al., 2006) myotubes, muscle lipolysis (Wolsk et al., 2010) and fatty acid uptake in humans (Kistner et al., 2025; Trinh et al., 2025), as well as fatty acid oxidation in L6 myotubes (Carey et al., 2006; Petersen et al., 2005), human myotubes (Al-Khalili et al., 2006) and in humans (Petersen et al., 2005; van Hall et al., 2003), reflecting the role of IL-6 as an allocator of energy substrates (Kistner et al., 2022). Taken together, these findings indicate that siIL6 myoblasts are not only less capable of maintaining basal metabolic activity but are also less metabolically adaptable to acute stimulation with rhIL-6.

Apart from the canonical JAK/STAT signalling pathway, IL-6 is an activator of several others, such as the PI3K and MAPK pathways (Eulenfeld et al., 2012; Hirano et al., 1997). In skeletal muscle cells, MAPK pathways are promyogenic and interconnected with IL-6 signalling (Baeza-Raja & Munoz-Canoves, 2004). The activation of p38 MAPK or ERK1/2 is associated with increased IL-6 expression in L6 myotubes (Green et al., 2011), C2C12 myotubes (Ito et al., 2018; Luo et al., 2003), C2C12 myoblasts (Baeza-Raja & Munoz-Canoves, 2004) and human skeletal muscle *in vivo* (Chan et al., 2004). Conversely, IL-6 has been shown to activate p38 MAPK or ERK1/2 signalling pathways in murine models (Manjavachi et al., 2010) and C2C12 myotubes (Brown et al., 2017; Fix et al., 2019; Weigert et al., 2007). In our cell model, the ERK1/2 and p38 MAPK signalling pathways, which have been reported to affect STAT3 phosphorylation at both tyrosine and serine residues (Eulenfeld et al., 2012; Jain et al., 1998; Xu et al., 2003), did not respond to rhIL-6 treatment or IL-6 silencing. While we did not explore the reasons for this apparent discrepancy, it is possible that it reflects functional differences between primary human skeletal muscle cells and skeletal muscle cell lines or differences in media composition, which have all been shown to have a major impact on experimental outcomes (Abdelmoez et al., 2020; Dolinar et al., 2018; Pirkmajer & Chibalin, 2011).

Myoblasts exhibited intrinsic secretory activity, which resulted in IL-6 concentrations in the cell culture medium of approximately 1 ng/mL. Although rhIL-6 was used in 50-fold higher concentration (50 ng/mL), pretreatment of myoblasts with NAbIL6, an anti-IL-6 antibody, enhanced rhIL-6-stimulated phosphorylation of STAT1^Tyr701^ and STAT3^Tyr705^. As concentration of NAbIL6 was sufficient to block 5 ng/mL rhIL-6, we can assume that intrinsic IL-6 was completely suppressed during the 44-h pretreatment. Based on this assumption we concluded that the presence of IL-6 even at low concentrations has a suppressive effect on the ability of myoblasts to respond to rhIL-6. This conclusion is supported by the observation that a 24-h pretreatment with 5 ng/mL rhIL-6 reduced an increase in phosphorylation of STAT1^Tyr701^ and STAT3^Tyr705^ upon stimulation with 50 ng/mL rhIL-6. Clearly, exposure to endogenous or exogenous IL-6 reduces response of cultured myoblasts to further stimulation with IL-6, even if IL-6 is used in much higher concentrations.

An inverse relationship between concentration of IL-6 and the abundance of IL-6Rα in myoblasts would provide an elegant mechanism by which gene silencing of IL-6 and neutralization of endogenous IL-6 with NAbIL6 increased response to rhIL-6. Interestingly, chronic exercise training results in downregulation of IL-6 and upregulation of IL-6Rα under basal conditions (Fischer et al., 2004). Moreover, a study with primary human myoblasts showed that a 24-h treatment with 10 pg/mL IL-6 upregulates IL-6Rα, whereas treatment with 10 ng/mL IL-6 or 48-h treatment with 10 pg/mL IL-6 downregulates it (Steyn et al., 2019), underscoring that IL-6 suppresses IL6-Rα expression in a concentration- and time-dependent manner. However, while gene silencing of IL-6 led to a trend towards upregulation of IL-6Rα mRNA in our experiments, we could not reliably detect IL-6Rα by immunoblotting. Gene silencing of IL-6Rα markedly suppressed the response to rhIL-6, which shows that the receptor was functional despite technical difficulties to detect it at the protein level. Our results therefore neither directly support nor refute the possibility that gene silencing of IL-6 increased the response to rhIL-6 by upregulating IL-6Rα in myoblasts.

Gene silencing of IL-6 did not increase the rhLIF-induced phosphorylation of STAT1^Tyr701^ and STAT3^Tyr705^, although LIF and IL-6 share part of signalling machinery, including IL-6Rβ (also known as gp130 or IL6ST), the signal transducing subunit of the LIF and IL-6 receptor complexes (Rose-John, 2018). Gene silencing of IL-6 also did not alter protein abundance of IL-6Rβ or mRNA expression of key components of the JAK/STAT pathway, including *JAK1*, *JAK2*, *STAT1*, *STAT3*, or *SOCS3*, which may explain why the response to rhLIF was not enhanced. No apparent change in the common receptor subunit or key downstream components of the common signalling pathway indirectly suggests that gene silencing of IL-6 altered signalling machinery that is specific to IL-6. If we consider that response to rhIL-6 was dependent on the expression of IL-6Rα, that gene silencing of IL-6 tended to upregulate IL-6Rα mRNA, and that a previous study showed that expression of IL-6Rα depends on concentration of IL-6 (Steyn et al., 2019), increased abundance of IL-6Rα in siIL6 myoblasts remains a realistic, although unproven, possibility.

Gene silencing of STAT3 increased basal and rhIL-6-induced STAT1^Tyr701^ levels, which shows that a partial suppression of the JAK/STAT pathway can in principle enhance the response to stimulation with rhIL-6. Consistent with our result, STAT3 deficiency promoted STAT1 phosphorylation or transcriptional activity in immune cells and mouse embryonic fibroblasts (Hirahara et al., 2015; Ho & Ivashkiv, 2006; Schiavone et al., 2011). However, while STAT3 exerted control over STAT1, gene silencing of STAT1 had no effect on STAT3 in myoblasts, demonstrating these transcription factors are not functionally equivalent. Indeed, although STAT1 and STAT3 share overlapping transcriptional targets, they also regulate distinct gene sets (Qing & Stark, 2004).

In conclusion, our findings demonstrate that a reduction in intrinsic IL-6 expression in human myoblasts increases responsiveness of the JAK/STAT pathway to stimulation with rhIL-6, which possibly represents a mechanism for regulating IL-6 signalling. In addition, they show that IL-6 deficiency suppresses both glucose and fatty acid oxidation in myoblasts, supporting the notion that IL-6 is a regulator of energy metabolism in skeletal muscle cells. Collectively, our results imply that intrinsic IL-6 acts as an autocrine/paracrine signal that restrains activation of the JAK/STAT pathway by extrinsic (circulating) IL-6, but acts synergistically with it to promote myoblast energy metabolism.

## Supporting information

qPCR raw data

Immunoblotting raw data

## Author contributions

Conceived and designed research: SP, KPP, ACR, ETK, AS; analyzed data: AS, KM, BŽBG, KD, UNM, KL, SP; performed experiments: AS, KM, BŽBG, KD, UNM; interpreted results of experiments: AS, ACR, ETK, KL, KPP, SP; prepared figures: AS, BŽBG, KD; drafted manuscript: AS, SP; edited and revised manuscript: KM, BŽBG, KD, UNM, ACR, ETK, KL, KPP; approved final version: AS, KM, BŽBG, KD, UNM, ACR, ETK, KL, KPP, SP.

## Funding

This study was supported with funding from the Slovenian Research and Innovation Agency (Grants P3-0043, P3-0314, J7-3153, J7-60125, Bilateral Programme BI-NO/25-27-004, and the Young Researchers’ Programme).

## Acknowledgements

The authors would like to acknowledge Ksenja Babič Benedik for the preparation of primary human skeletal muscle cell cultures, and Hege Gilbø Bakke, Hilde Nilsen and Åse-Karine Fjeldheim for their contribution to the study. The authors also wish to thank Professor Alexander V. Chibalin, Professor Marc Gilbert, and Blaž Kociper, MD, for thorough reading of the manuscript.

## Use of artificial intelligence

A completed, fully written version of the manuscript, which had been prepared without the use of AI, was subjected to assessment by the Nature research assistant and InstaText with the purpose of evaluating English grammar and clarity of the text. Suggestions by these tools were taken into consideration, but all writing was done by the authors.

## Notes

### Competing Interest Statement

The authors have declared no competing interest.

