## Supplementary material for "Intrinsic IL-6 expression reduces rhIL-6-induced JAK/STAT activation and promotes glucose and oleic acid oxidation in cultured human myoblasts": Immunoblotting raw data

### Immunoblotting

#### Experimental procedure

**Lysis buffer**: Laemmli buffer (62.5 mM Tris-HCl (pH 6.8), 10% (w/v) glycerol, 2% (w/v) sodium dodecyl sulfate (SDS), 5% (v/v) 2-mercaptoethanol, 0.002% (w/v) bromophenol blue)

**Gel:** Criterion XT 4-12% Bis-Tris Precast Gel (#3450123, #3450124, Bio-Rad, Hercules, CA, USA)

**Electrophoresis:** SDS-PAGE in XT MES Running Buffer (#1610789, Bio-Rad, Hercules, CA, USA) at a constant voltage 200 V

**Membrane:** Immobilon-P Membrane, polyvinylidene fluoride (PVDF) (#IPVH85R, Merck Millipore, Burlington, MA, USA)

**Transfer**: in transfer buffer (31 mM Tris, 0.24 M glycine, 10% (v/v) methanol, and 0.01% (w/v) SDS) at a constant voltage 100 V

**Molecular weight marker:** Amersham ECL Full-Range Molecular Weight Marker (#RPN800E, Cytiva, Marlborough, MA, USA)

**Sample loading and transfer control:** Ponceau S (0.1% (w/v) in 5% (v/v) acetic acid) staining

**Substrate:** Immobilon Crescendo Western HRP Substrate (#WBLUR0500, Merck Millipore, Burlington, MA, USA)

**Imaging system:** Fusion FX (Vilber, Marne-la-Vallée, France)

**Antibodies:**

| | | Primary Antibody | | | | Secondary Antibody |
|---|---|---|---|---|---|---|
| **Target** | **Size (kDa)** | **Supplier** | **Catalog No.** | **Research Resource Identifier (RRID)** | **Dilution** | **Dilution** |
| **Phospho-STAT3 (Tyr705)** | 79, 86 | Cell Signaling Technology | 9145 | AB_2491009 | 1:1000 | 1:10000 |
| **Phospho-STAT3 (Ser727)** | 86 | Cell Signaling Technology | 9134 | AB_331589 | 1:1000 | 1:15000 |
| **Phospho-STAT1 (Tyr701)** | 84, 91 | Cell Signaling Technology | 9167 | AB_561284 | 1:1000 | 1:25000 |
| **STAT1** | 84, 91 | Cell Signaling Technology | 9172 | AB_2198300 | 1:1000 | 1:25000 |
| **STAT3** | 79, 86 | Cell Signaling Technology | 4904 | AB_331269 | 1:1000 | 1:15000 |
| **Phospho-p38 MAPK (Thr180/Tyr182)** | 43 | Cell Signaling Technology | 4511 | AB_2139682 | 1:1000 | 1:20000 |
| **Phospho-p44/42 MAPK (ERK1/2) (Thr202/Tyr204)** | 44, 42 | Cell Signaling Technology | 4370 | AB_2315112 | 1:2000 | 1:15000 |
| **GP130*** | 130 | Cell Signaling Technology | 3732 | AB_2125953 | 1:1000 | 1:10000 |
| **IL-6Rα/CD126** | 70-100 | Cell Signaling Technology | 18935 | AB_3741500 | 1:500 | 1:5000 |
| **SOCS3** | 28 | Cell Signaling Technology | 52113 | AB_2799408 | 1:500 | 1:5000 |

Red frames indicate all analyzed bands. Blue frames indicate bands shown in the Figures.

#### Figure 1e, f: Silencing *IL6/IL6Rα/IL6 + IL6R* + rhIL-6 stimulation

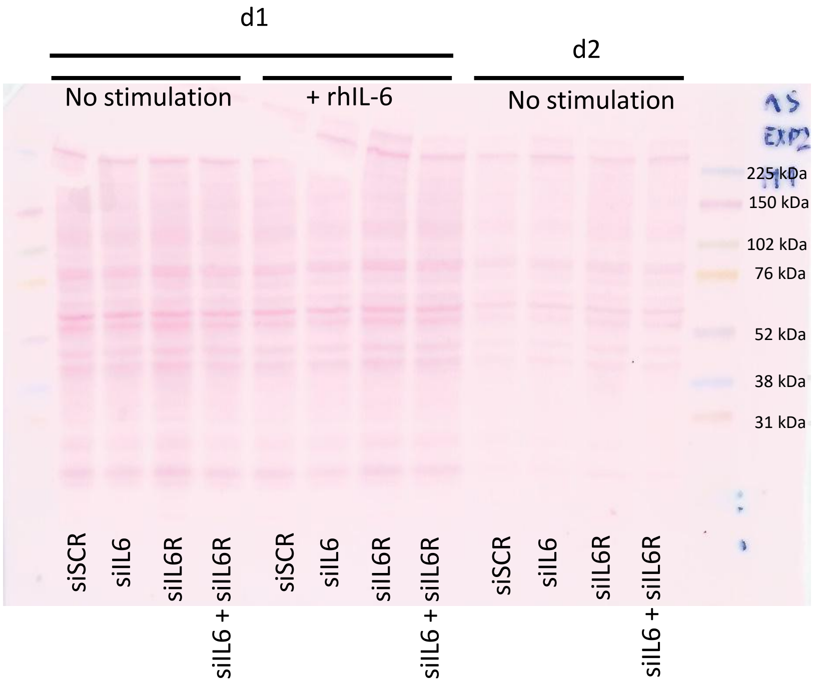


**pSTAT1 (Tyr701)**

- **Basal**
- **Basal + rhIL-6**

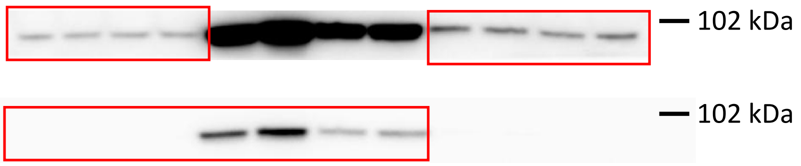


**pSTAT3 (Tyr705)**

- **Basal**
- **Basal + rhIL-6**

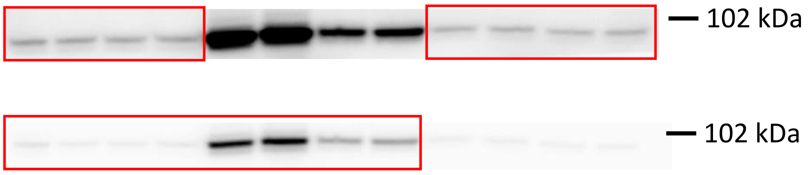

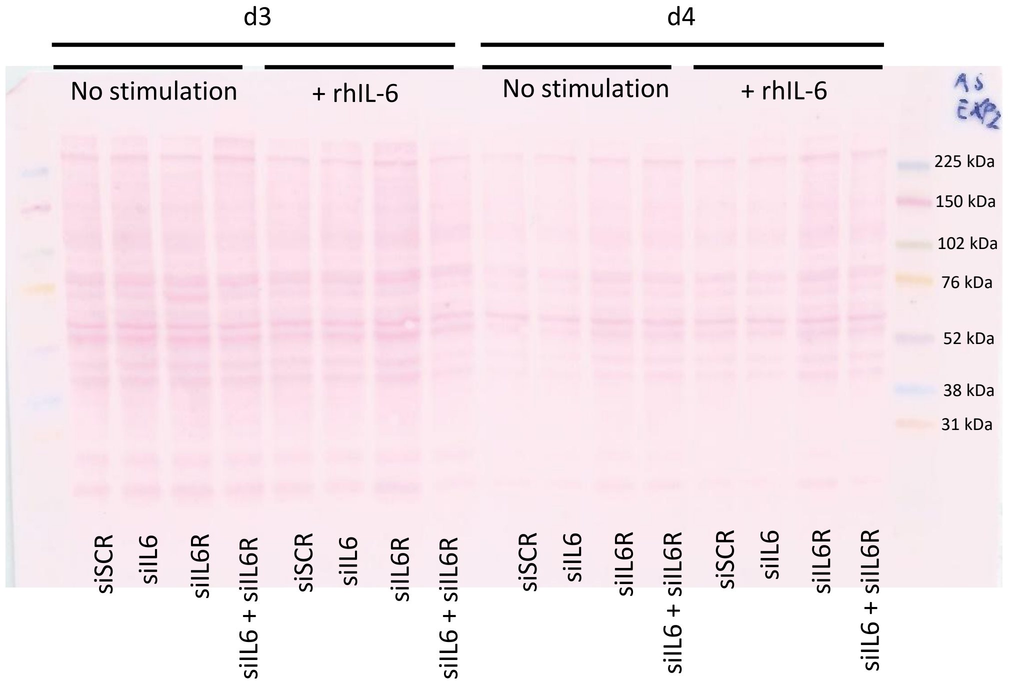


**pSTAT1 (Tyr701)**

- **Basal**

- **Basal + rhIL-6**

**pSTAT3 (Tyr705)**

- **Basal**

- **Basal + rhIL-6**

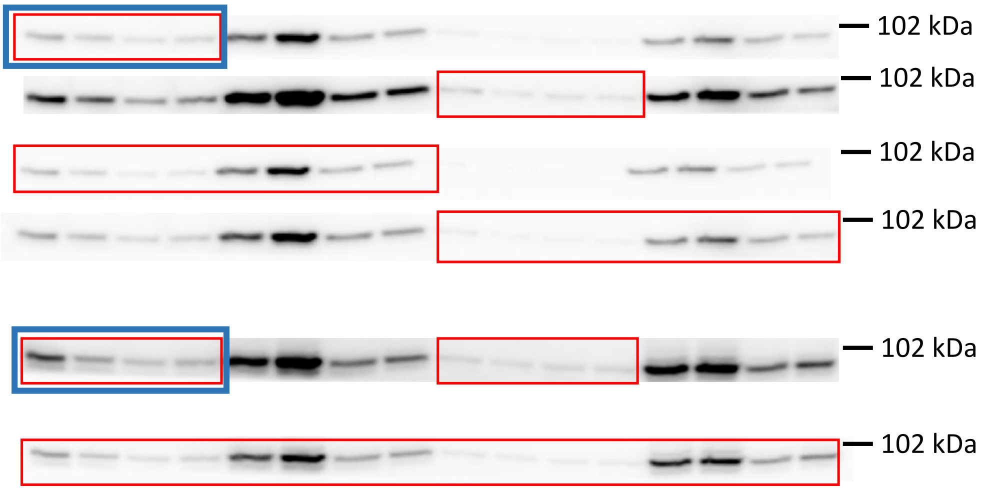

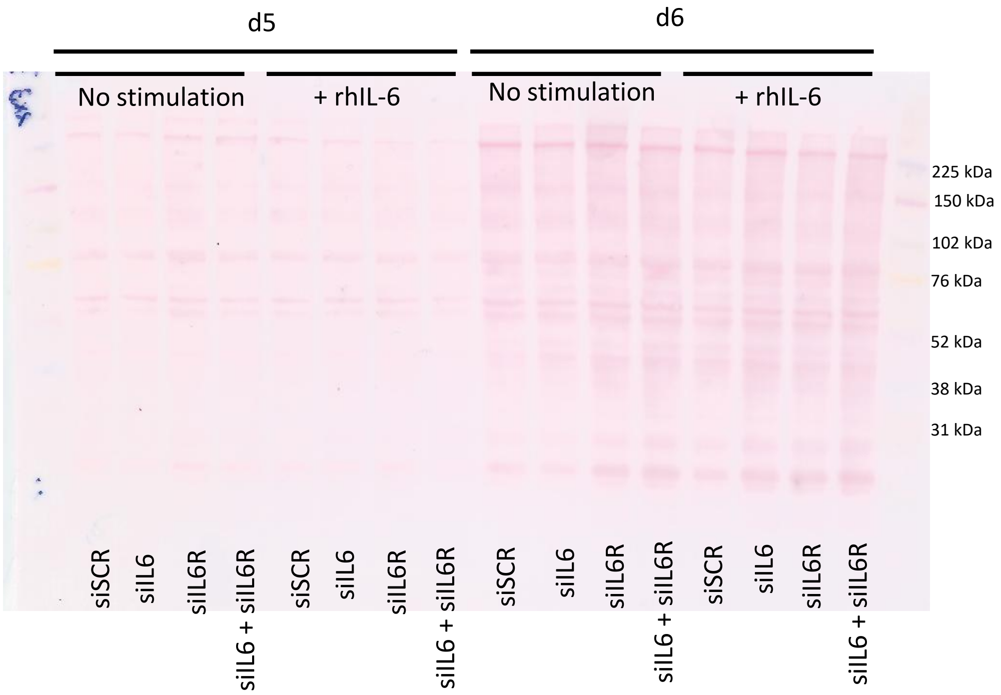
d5
d6
No stimulation
+ rhIL-6
No stimulation
+ rhIL-6
225 kDa
150 kDa
102 kDa
76 kDa
52 kDa
38 kDa
31 kDa
siSCR
siIL6
siIL6R
siIL6 + siIL6R
siSCR
siIL6
siIL6R
siIL6 + siIL6R
siSCR
siIL6
siIL6R
siIL6 + siIL6R
siSCR
siIL6
siIL6R
siIL6 + siIL6R

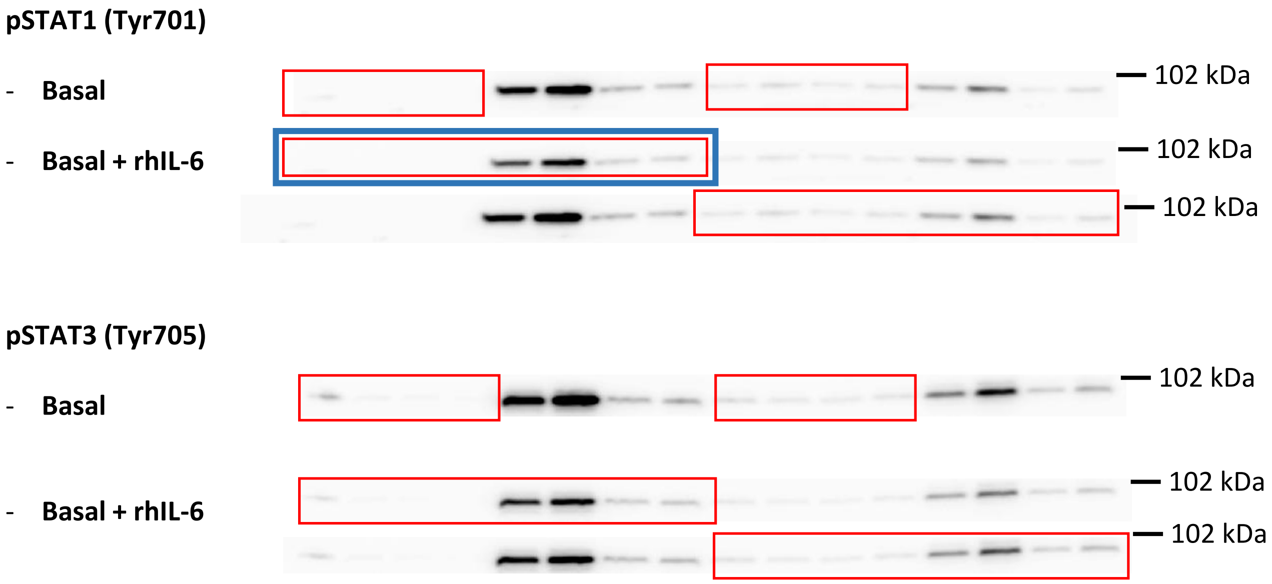
pSTAT1 (Tyr701)
- Basal
- Basal + rhIL-6
102 kDa
102 kDa
102 kDa
pSTAT3 (Tyr705)
- Basal
- Basal + rhIL-6
102 kDa
102 kDa
102 kDa

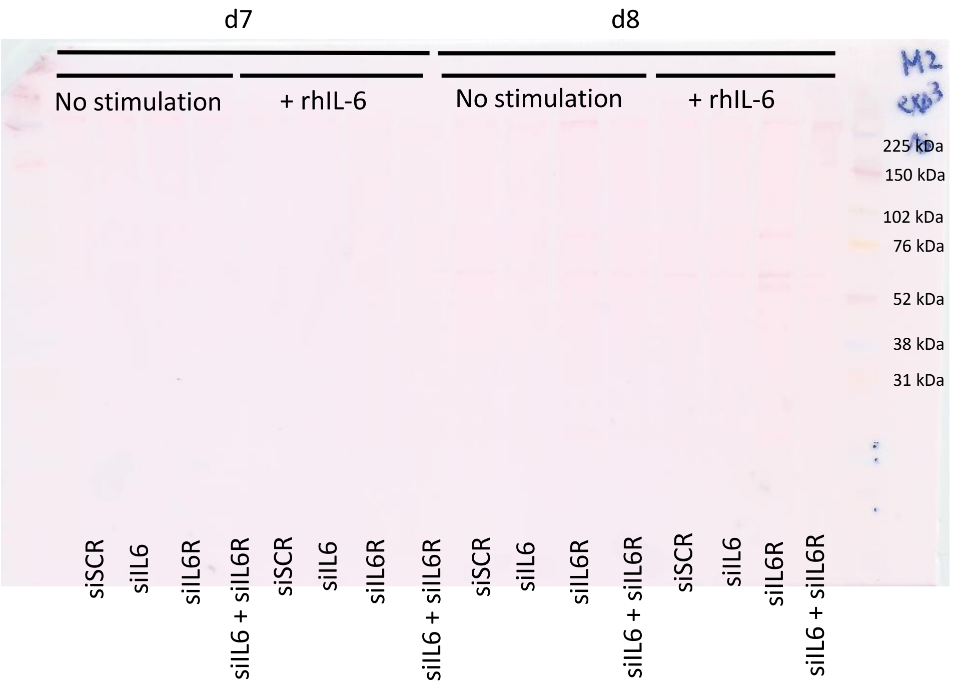


**pSTAT1 (Tyr701)**

- **Basal**

Not detectable.

- **Basal + rhIL-6**

Excluded due to a technical artefact.

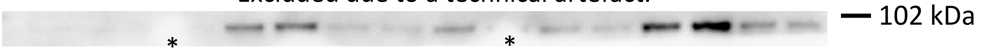


**pSTAT3 (Tyr705)**

- **Basal**
- **Basal + rhIL-6**

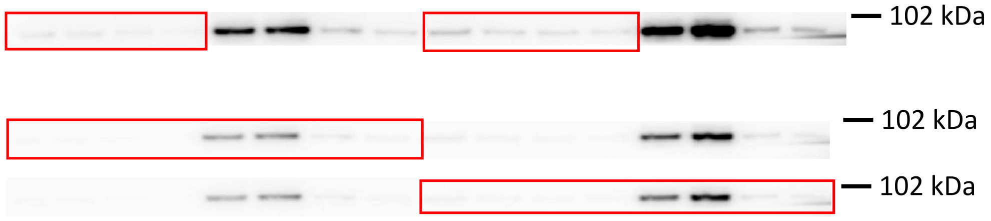

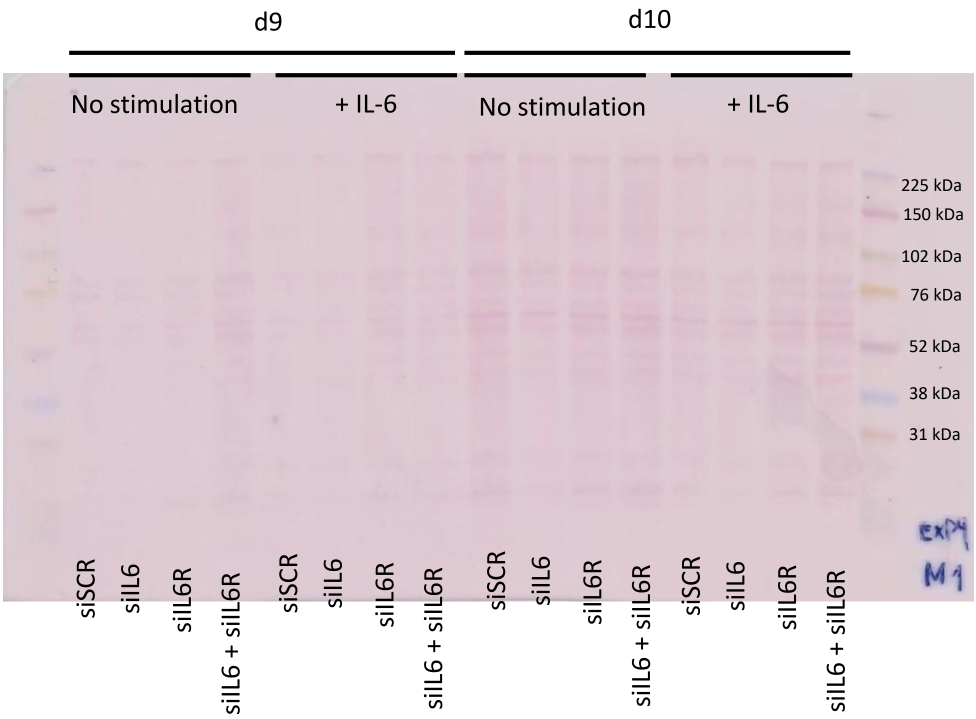
d9
d10
No stimulation
+ IL-6
No stimulation
+ IL-6
225 kDa
150 kDa
102 kDa
76 kDa
52 kDa
38 kDa
31 kDa
ExP4
M1
siSCR
siIL6
siIL6R
siIL6 + siIL6R
siSCR
siIL6
siIL6R
siIL6 + siIL6R
siSCR
siIL6
siIL6R
siIL6 + siIL6R
siSCR
siIL6
siIL6R
siIL6 + siIL6R

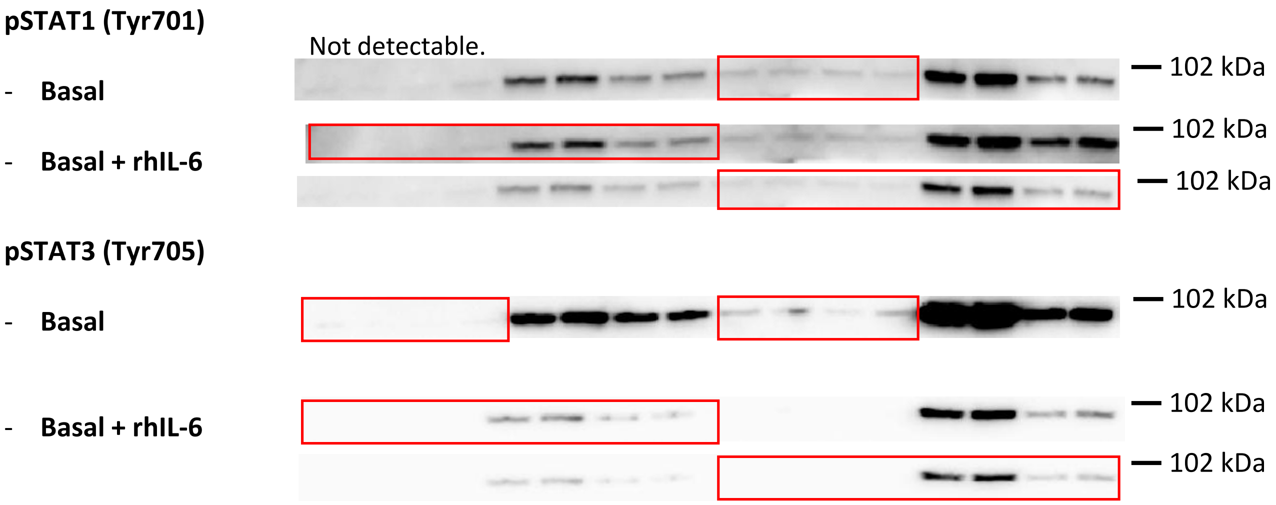
pSTAT1 (Tyr701)
- Basal
- Basal + rhIL-6
Not detectable.
102 kDa
102 kDa
102 kDa
pSTAT3 (Tyr705)
- Basal
- Basal + rhIL-6
102 kDa
102 kDa
102 kDa

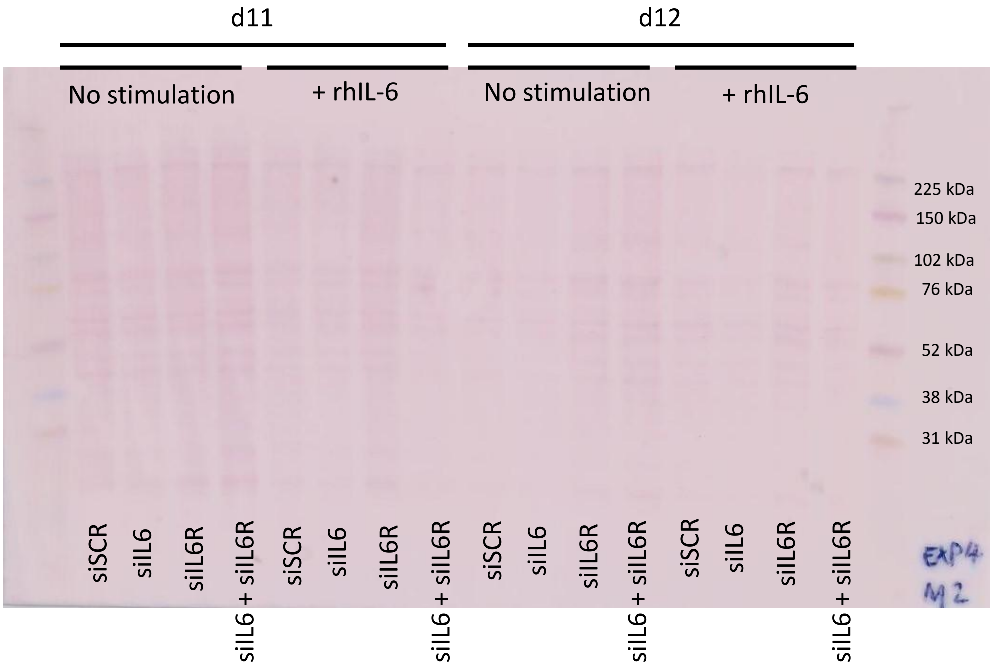


**pSTAT1 (Tyr701)**

- **Basal**
- **Basal + rhIL-6**

**pSTAT3 (Tyr705)**

- **Basal**
- **Basal + rhIL-6**

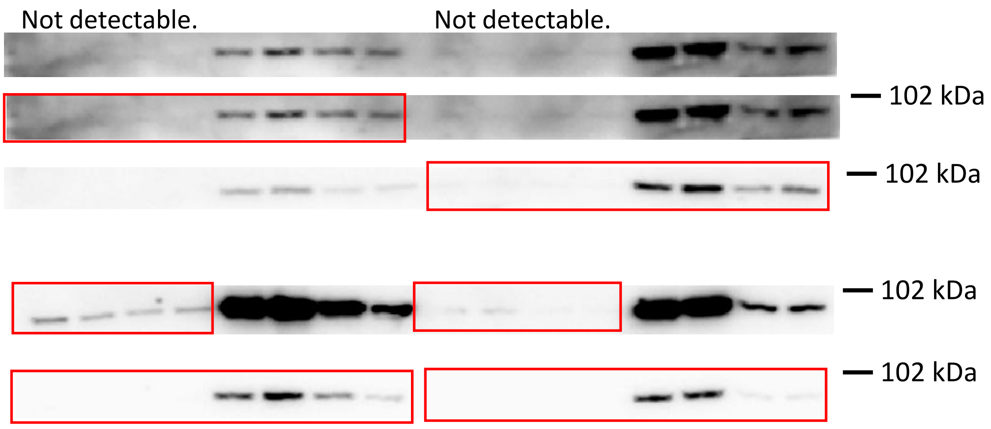

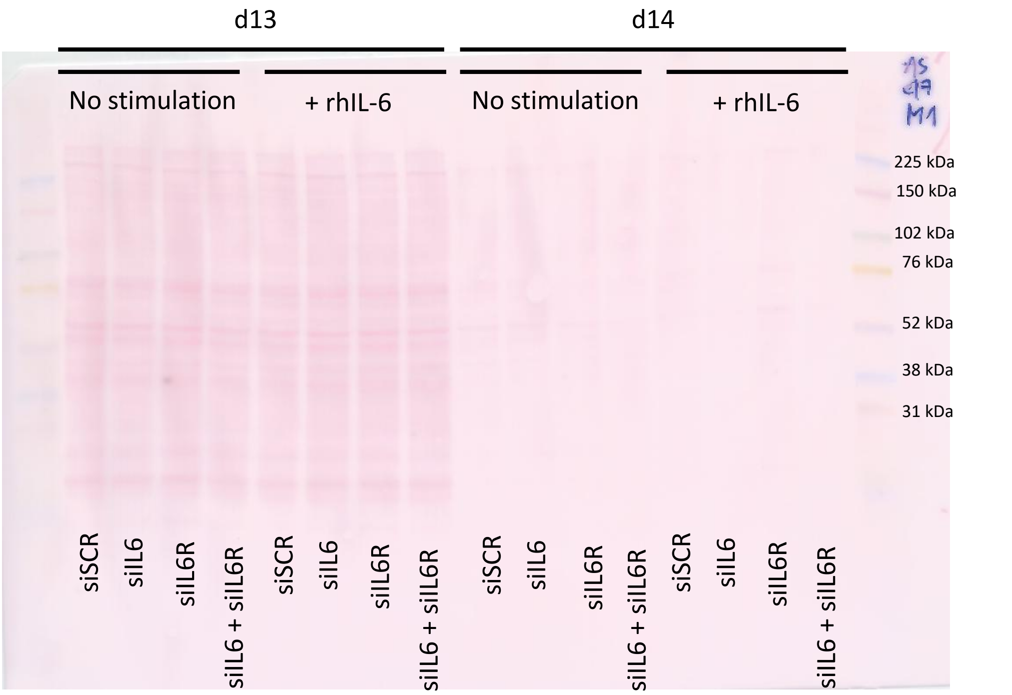


**pSTAT1 (Tyr701)**

- **Basal**

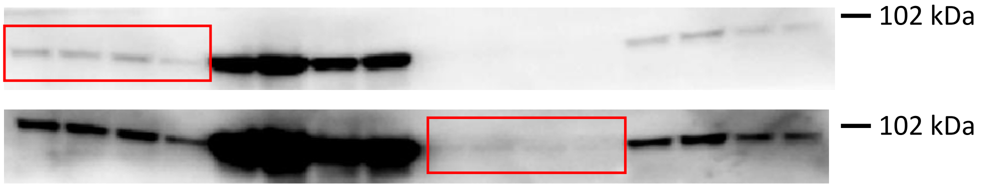


- **Basal + rhIL-6**

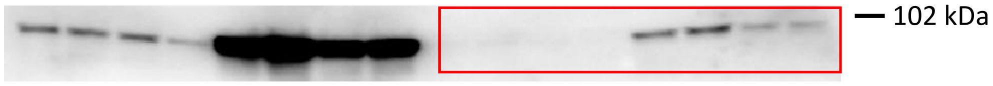


**pSTAT3 (Tyr705)**

- **Basal**

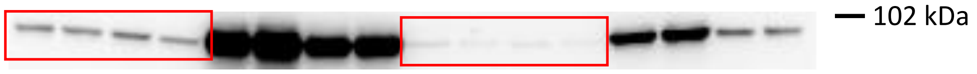


- **Basal + rhIL-6**

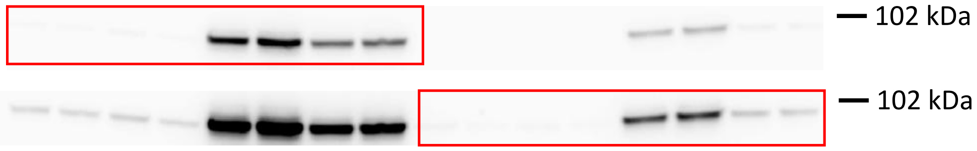

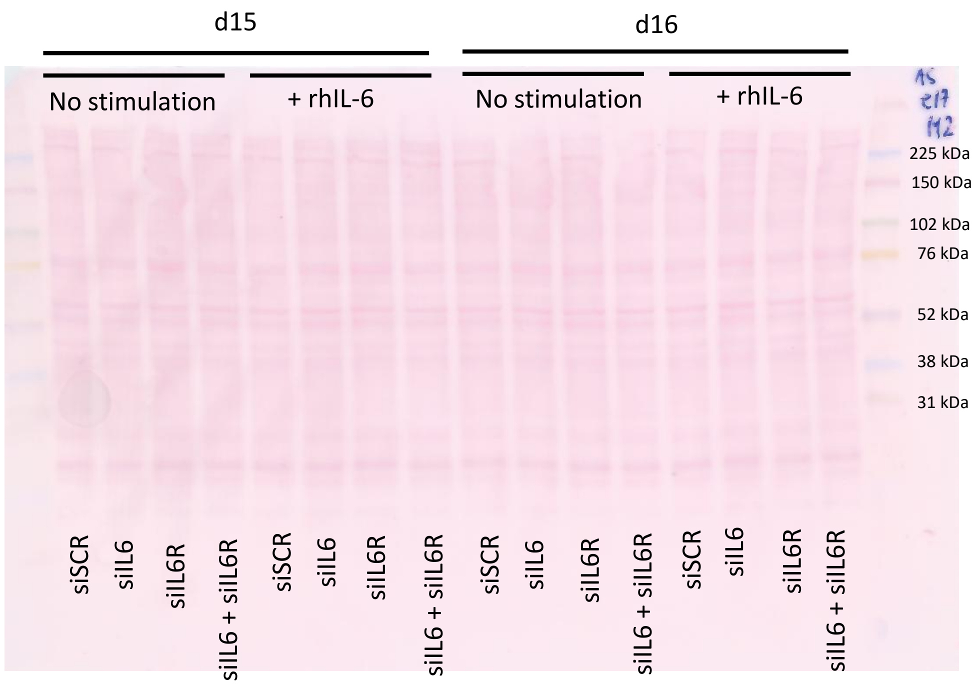


**pSTAT1 (Tyr701)**

- **Basal**
- **Basal + rhIL-6**

**pSTAT3 (Tyr705)**

- **Basal**
- **Basal + rhIL-6**

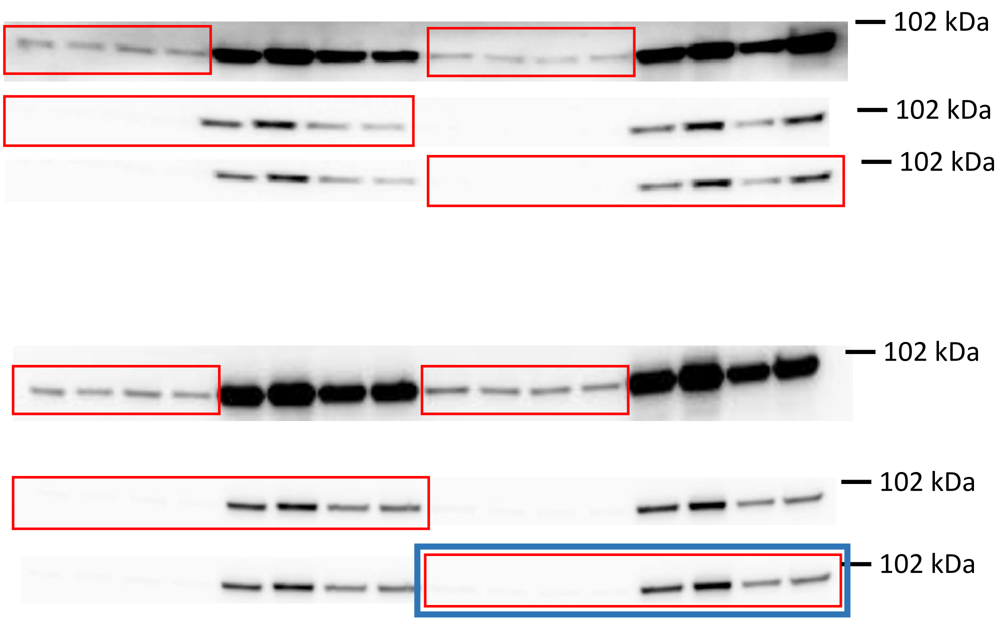

**Figure 2a, c – f: Time-course experiments**

d1

siSCR siIL6

IL-6 [min]

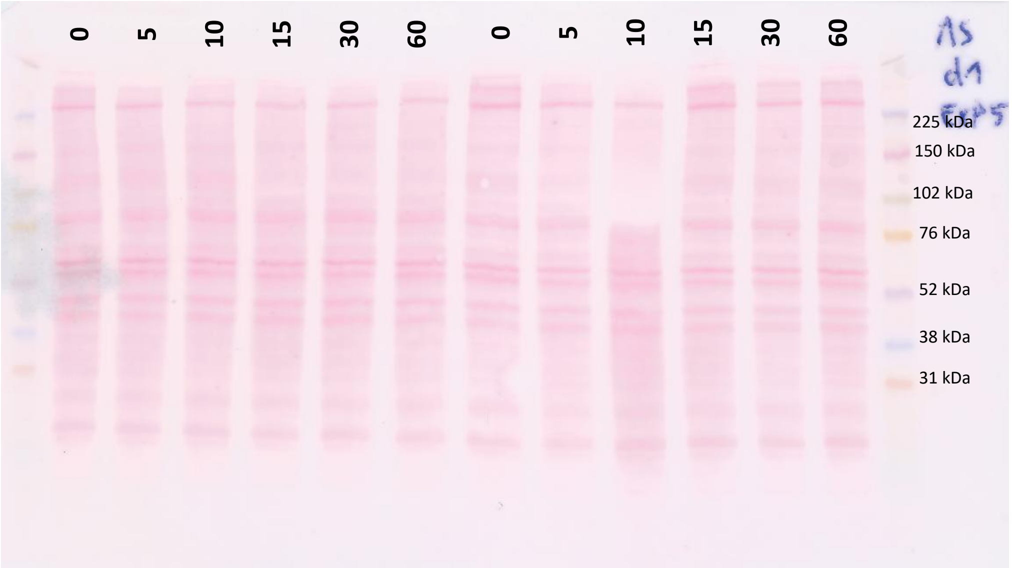


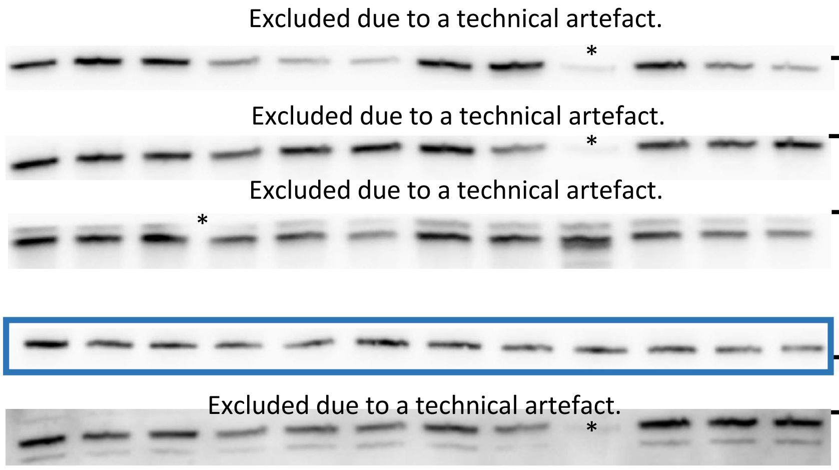


pSTAT3 (Tyr705) — 102 kDa

tSTAT3 — 102 kDa

pERK1/2 (Thr202/Tyr204) — 52 kDa

Pp38 MAPK (Thr202/Tyr204) — 38 kDa

pSTAT3 (Ser727) — 102 kDa

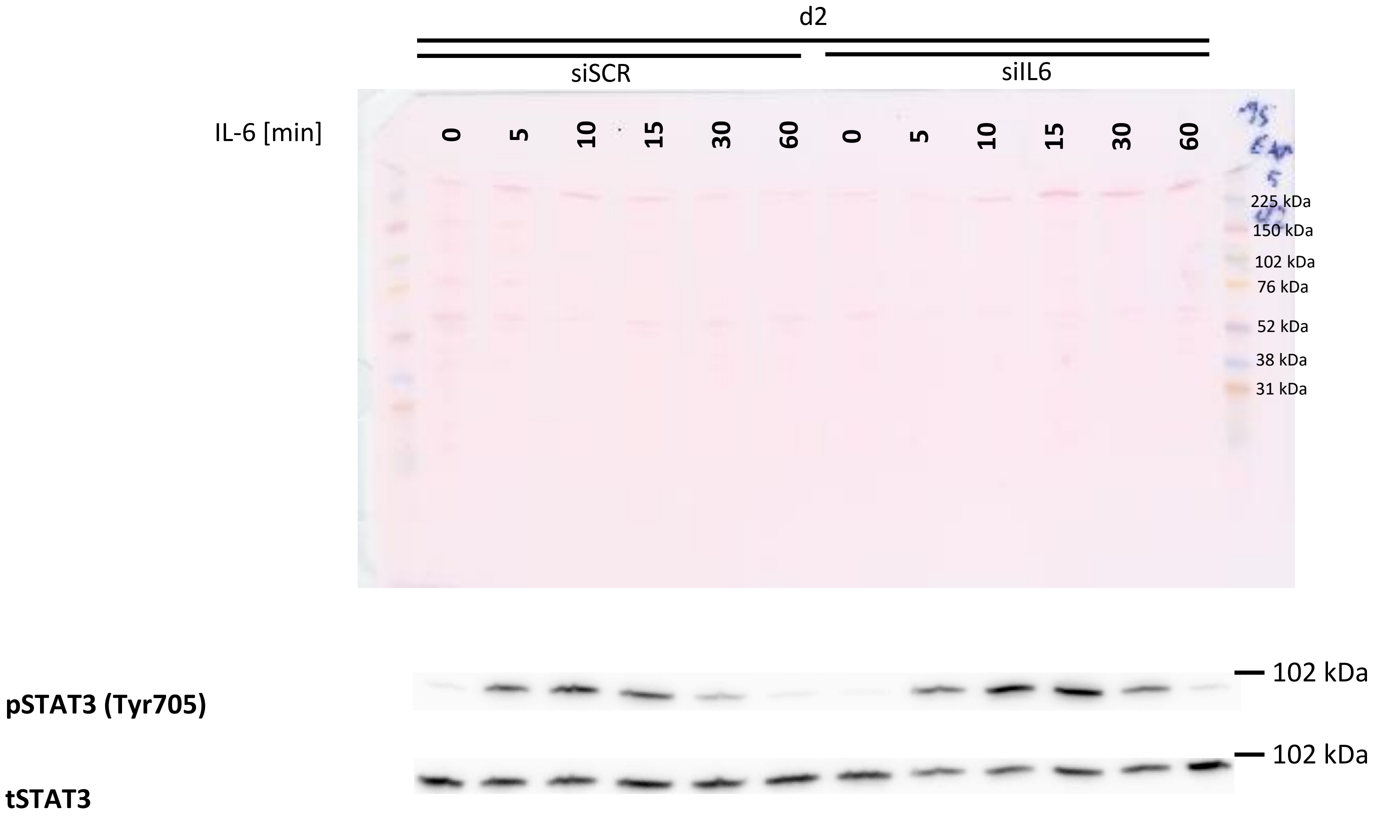
d2
siSCR
siIL6
IL-6 [min]
0 5 10 15 30 60 0 5 10 15 30 60
225 kDa
150 kDa
102 kDa
76 kDa
52 kDa
38 kDa
31 kDa
pSTAT3 (Tyr705)
102 kDa
tSTAT3
102 kDa

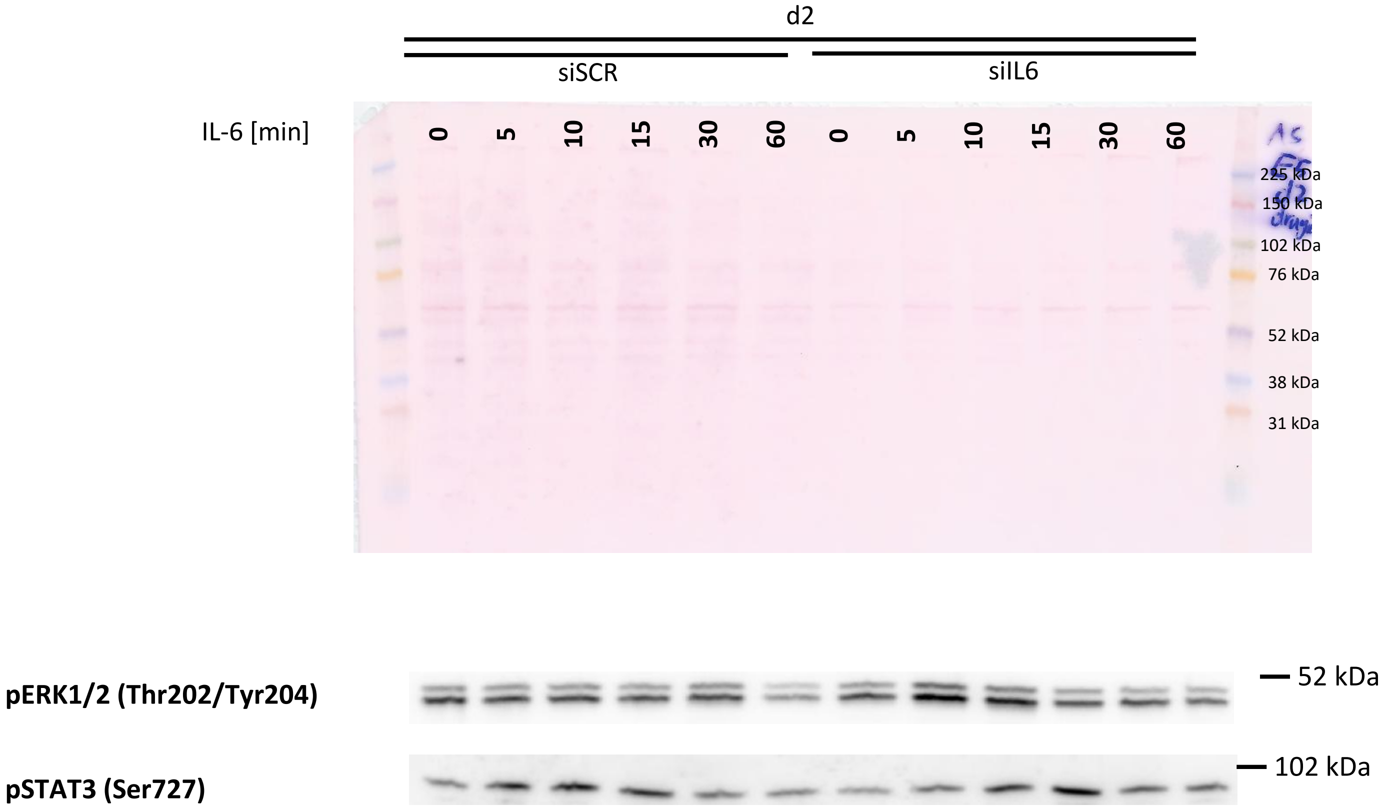
d2
siSCR
siIL6
IL-6 [min]
0 5 10 15 30 60 0 5 10 15 30 60
225 kDa
150 kDa
102 kDa
76 kDa
52 kDa
38 kDa
31 kDa
pERK1/2 (Thr202/Tyr204)
52 kDa
pSTAT3 (Ser727)
102 kDa

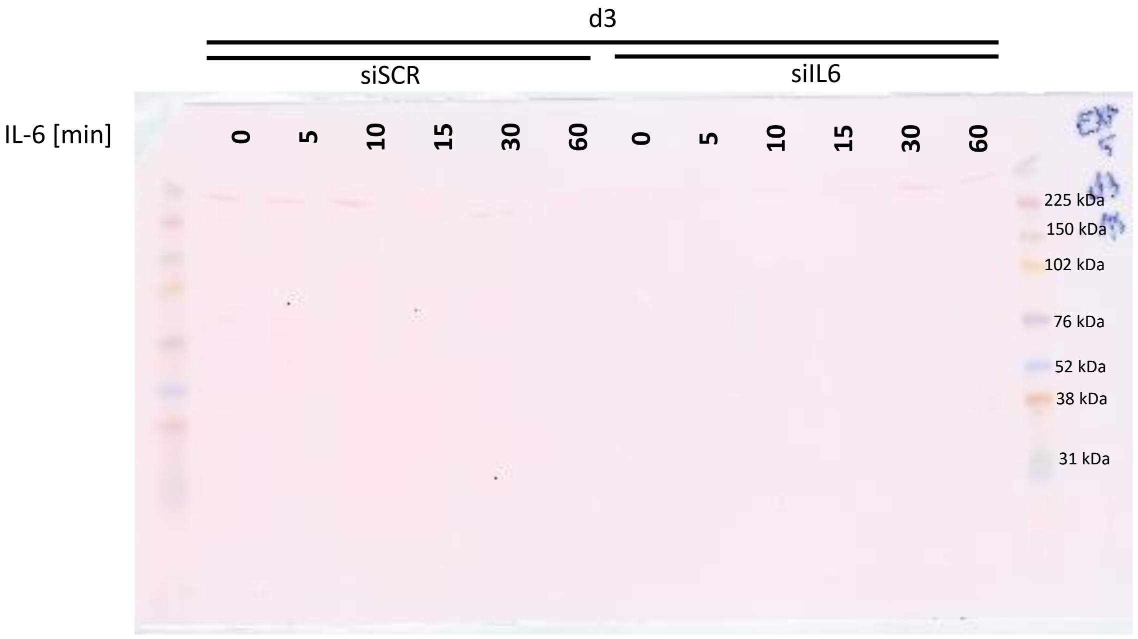

d3
siSCR
siIL6
IL-6 [min]
0
5
10
15
30
60
0
5
10
15
30
60
225 kDa
150 kDa
102 kDa
76 kDa
52 kDa
38 kDa
31 kDa


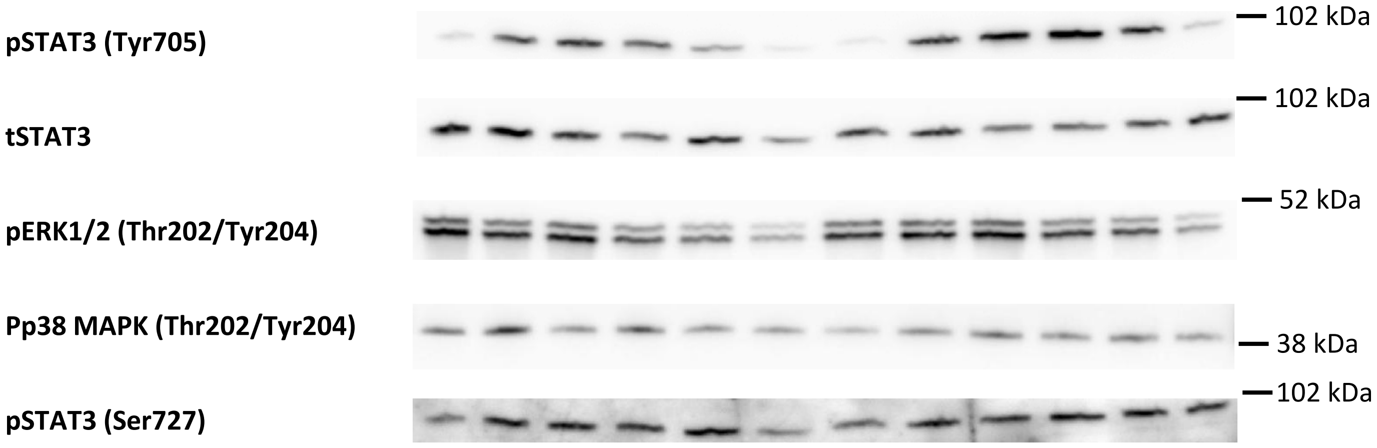

pSTAT3 (Tyr705)
102 kDa
tSTAT3
102 kDa
pERK1/2 (Thr202/Tyr204)
52 kDa
Pp38 MAPK (Thr202/Tyr204)
38 kDa
pSTAT3 (Ser727)
102 kDa

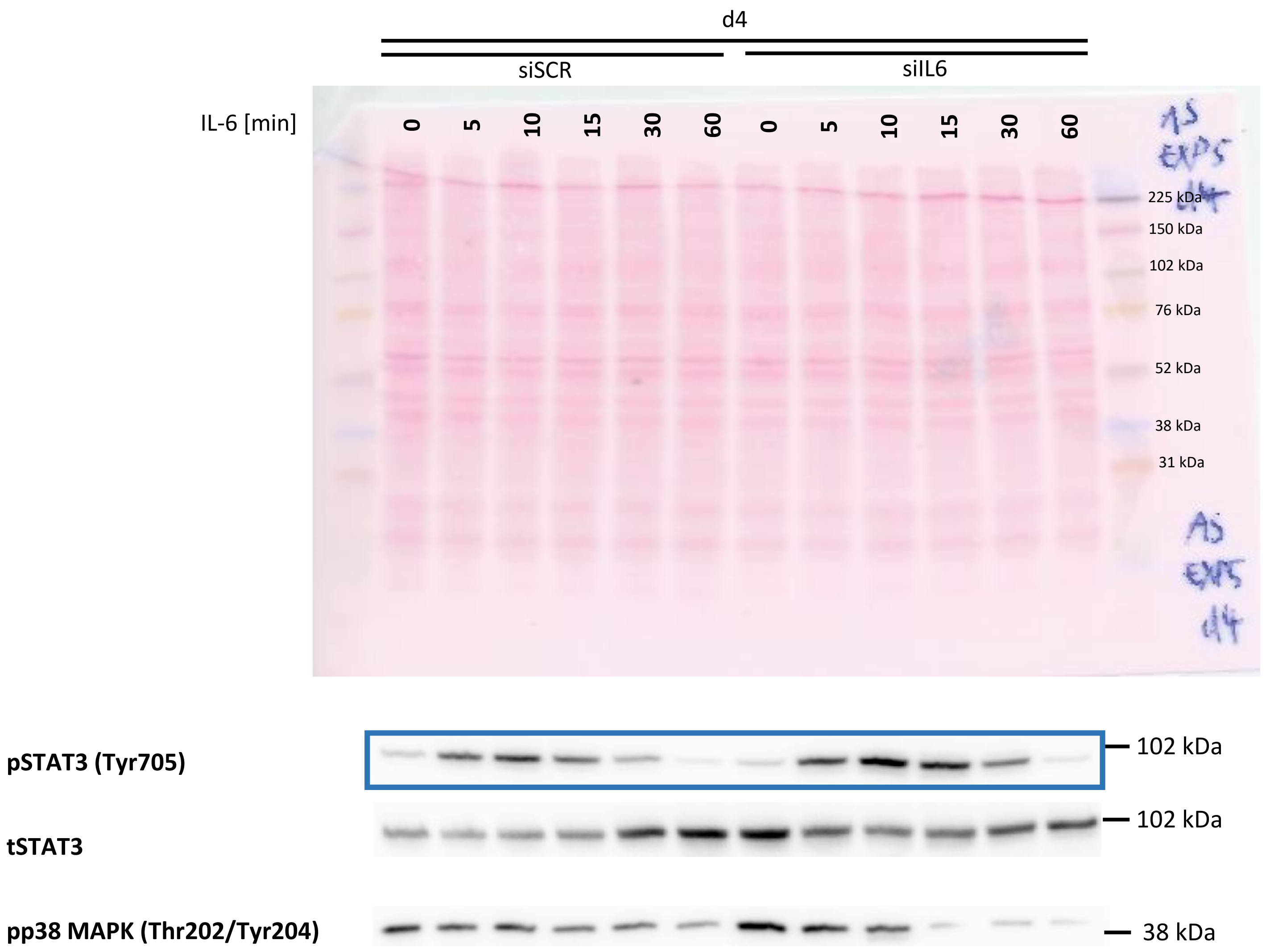
d4
siSCR
siIL6
IL-6 [min]
0 5 10 15 30 60 0 5 10 15 30 60
225 kDa
150 kDa
102 kDa
76 kDa
52 kDa
38 kDa
31 kDa
pSTAT3 (Tyr705)
102 kDa
tSTAT3
102 kDa
pp38 MAPK (Thr202/Tyr204)
38 kDa

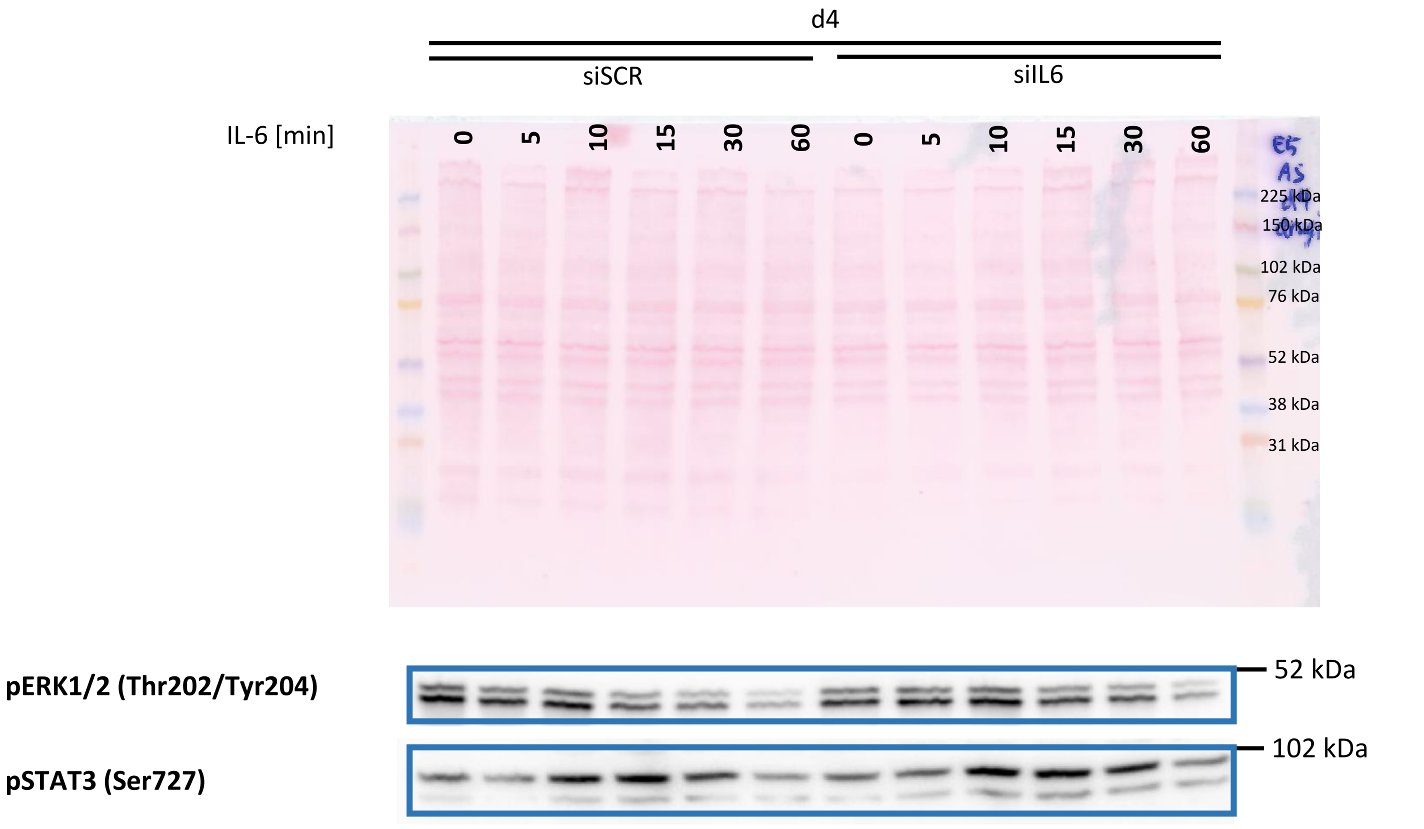
d4
siSCR
siIL6
IL-6 [min]
0 5 10 15 30 60 0 5 10 15 30 60
225 kDa
150 kDa
102 kDa
76 kDa
52 kDa
38 kDa
31 kDa
pERK1/2 (Thr202/Tyr204)
52 kDa
pSTAT3 (Ser727)
102 kDa

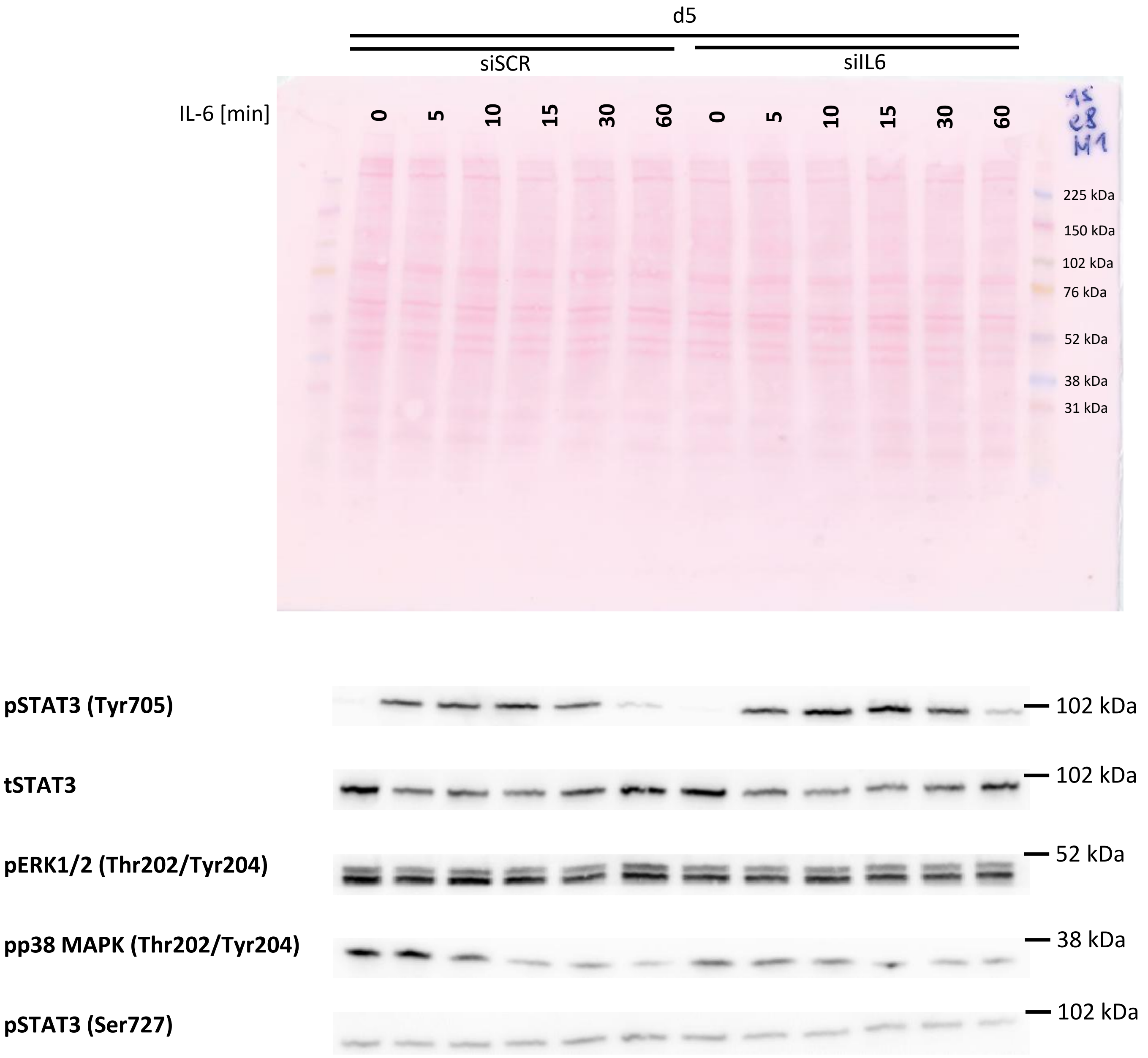
d5
siSCR
siIL6
IL-6 [min]
0
5
10
15
30
60
0
5
10
15
30
60
225 kDa
150 kDa
102 kDa
76 kDa
52 kDa
38 kDa
31 kDa
pSTAT3 (Tyr705)
102 kDa
tSTAT3
102 kDa
pERK1/2 (Thr202/Tyr204)
52 kDa
pp38 MAPK (Thr202/Tyr204)
38 kDa
pSTAT3 (Ser727)
102 kDa

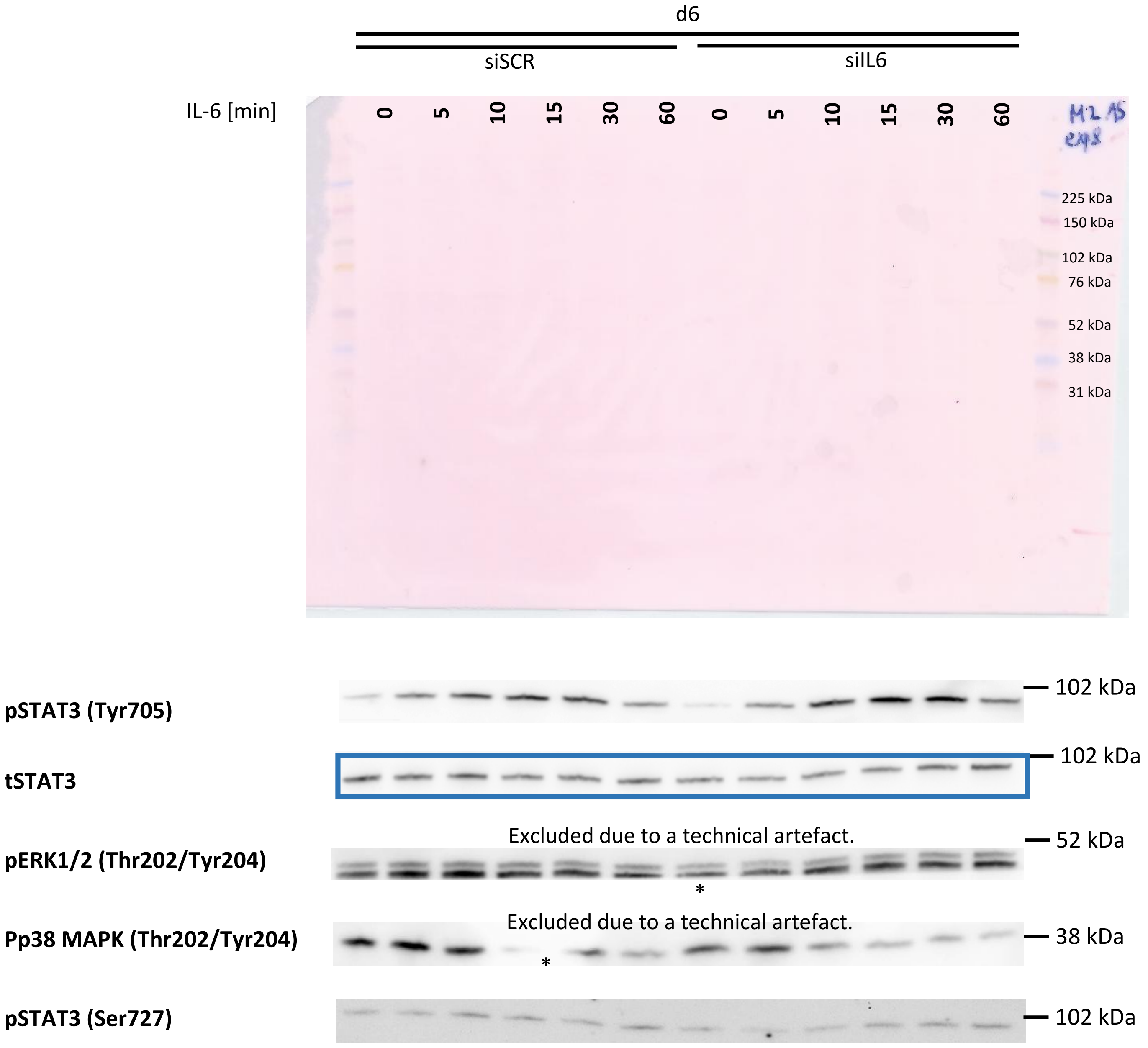
d6
siSCR
siIL6
IL-6 [min]
0
5
10
15
30
60
0
5
10
15
30
60
225 kDa
150 kDa
102 kDa
76 kDa
52 kDa
38 kDa
31 kDa
pSTAT3 (Tyr705)
102 kDa
tSTAT3
102 kDa
Excluded due to a technical artefact.
pERK1/2 (Thr202/Tyr204)
52 kDa
*
Excluded due to a technical artefact.
Pp38 MAPK (Thr202/Tyr204)
38 kDa
*
pSTAT3 (Ser727)
102 kDa

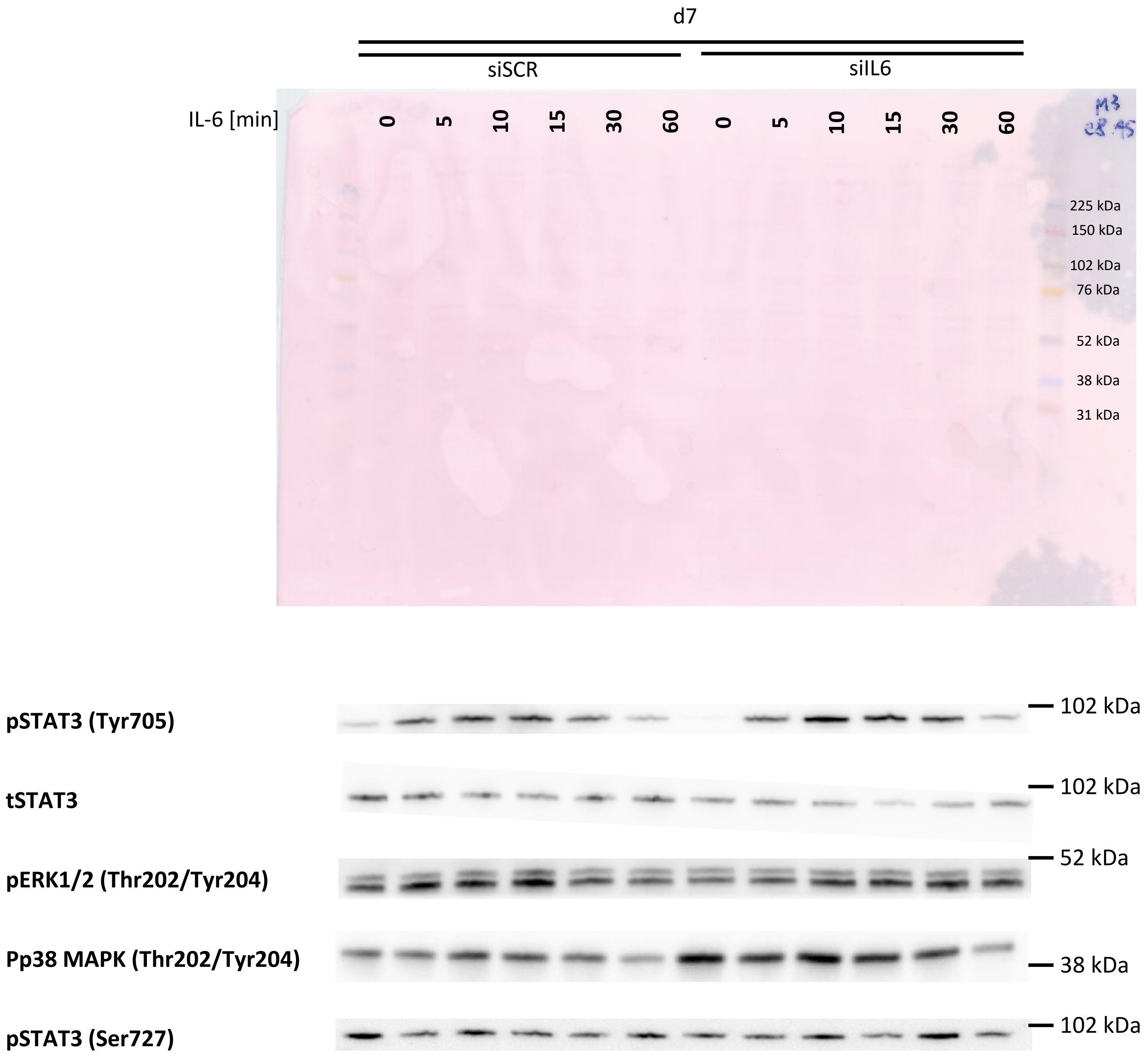
d7
siSCR
siIL6
IL-6 [min]
0 5 10 15 30 60 0 5 10 15 30 60
225 kDa
150 kDa
102 kDa
76 kDa
52 kDa
38 kDa
31 kDa
pSTAT3 (Tyr705)
102 kDa
tSTAT3
102 kDa
pERK1/2 (Thr202/Tyr204)
52 kDa
Pp38 MAPK (Thr202/Tyr204)
38 kDa
pSTAT3 (Ser727)
102 kDa

**Figure 3a, d – g: Dose-response experiments**

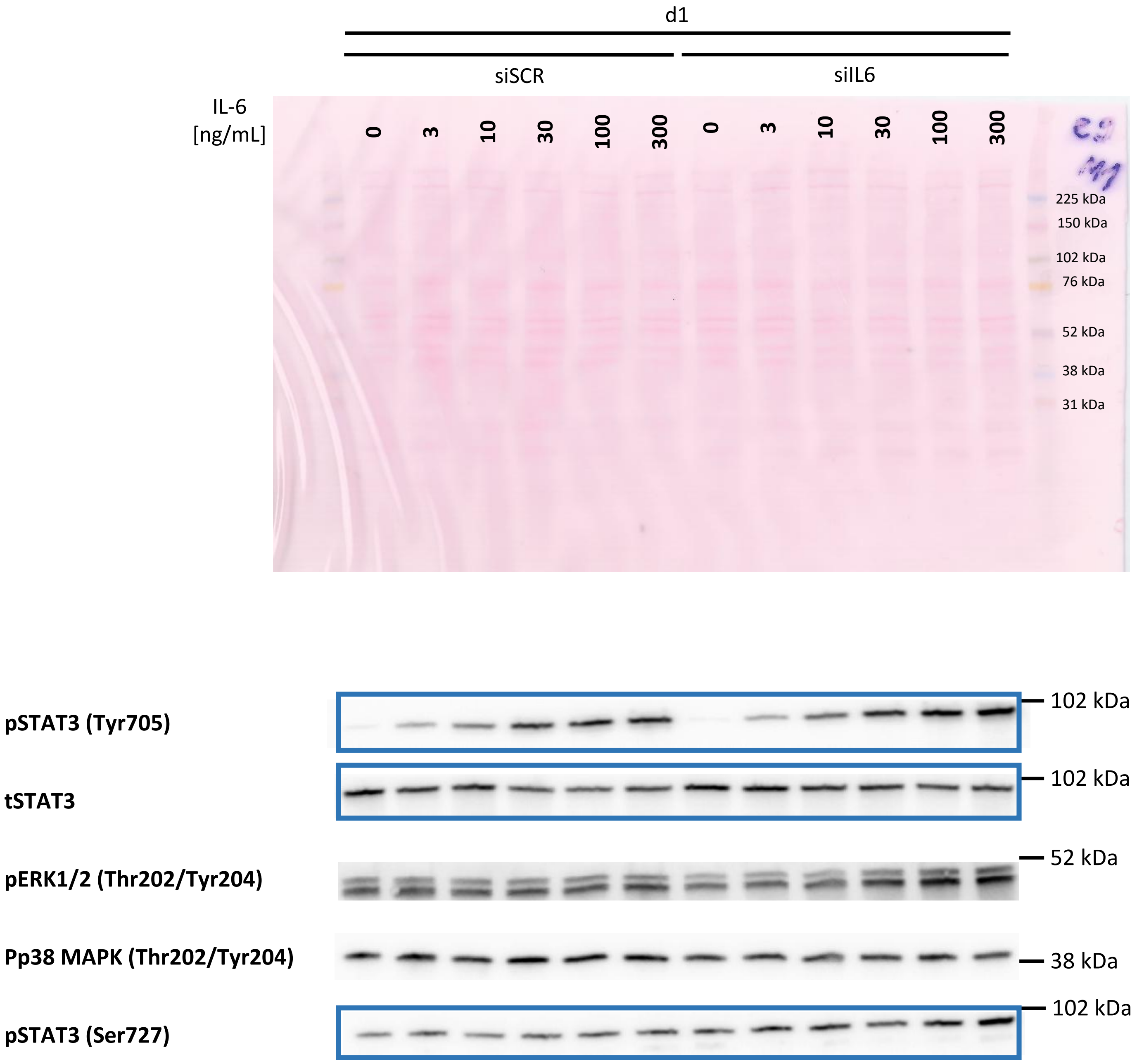

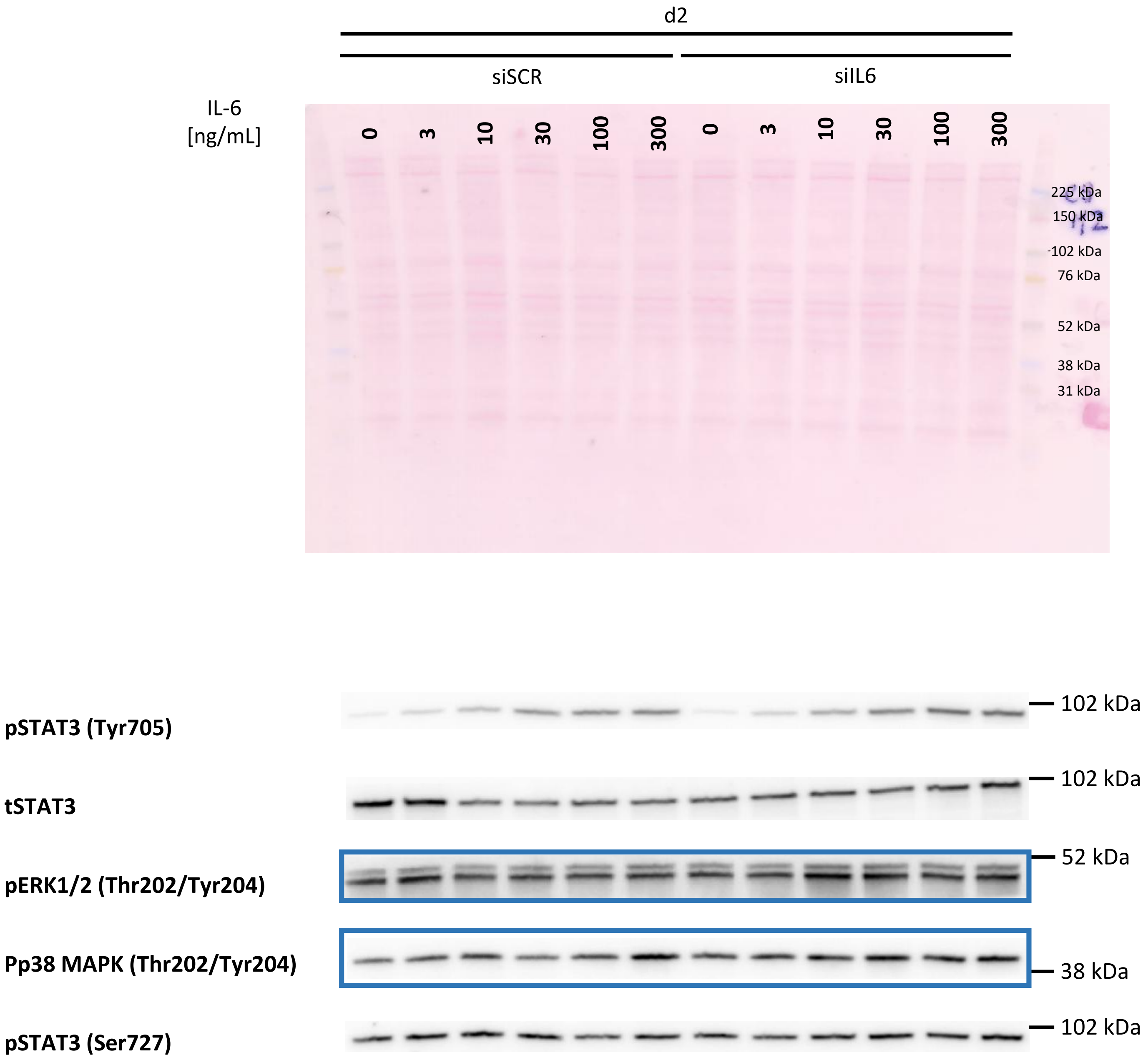
d2
siSCR
siIL6
IL-6 [ng/mL]
0
3
10
30
100
300
0
3
10
30
100
300
225 kDa
150 kDa
102 kDa
76 kDa
52 kDa
38 kDa
31 kDa
pSTAT3 (Tyr705)
102 kDa
tSTAT3
102 kDa
pERK1/2 (Thr202/Tyr204)
52 kDa
Pp38 MAPK (Thr202/Tyr204)
38 kDa
pSTAT3 (Ser727)
102 kDa

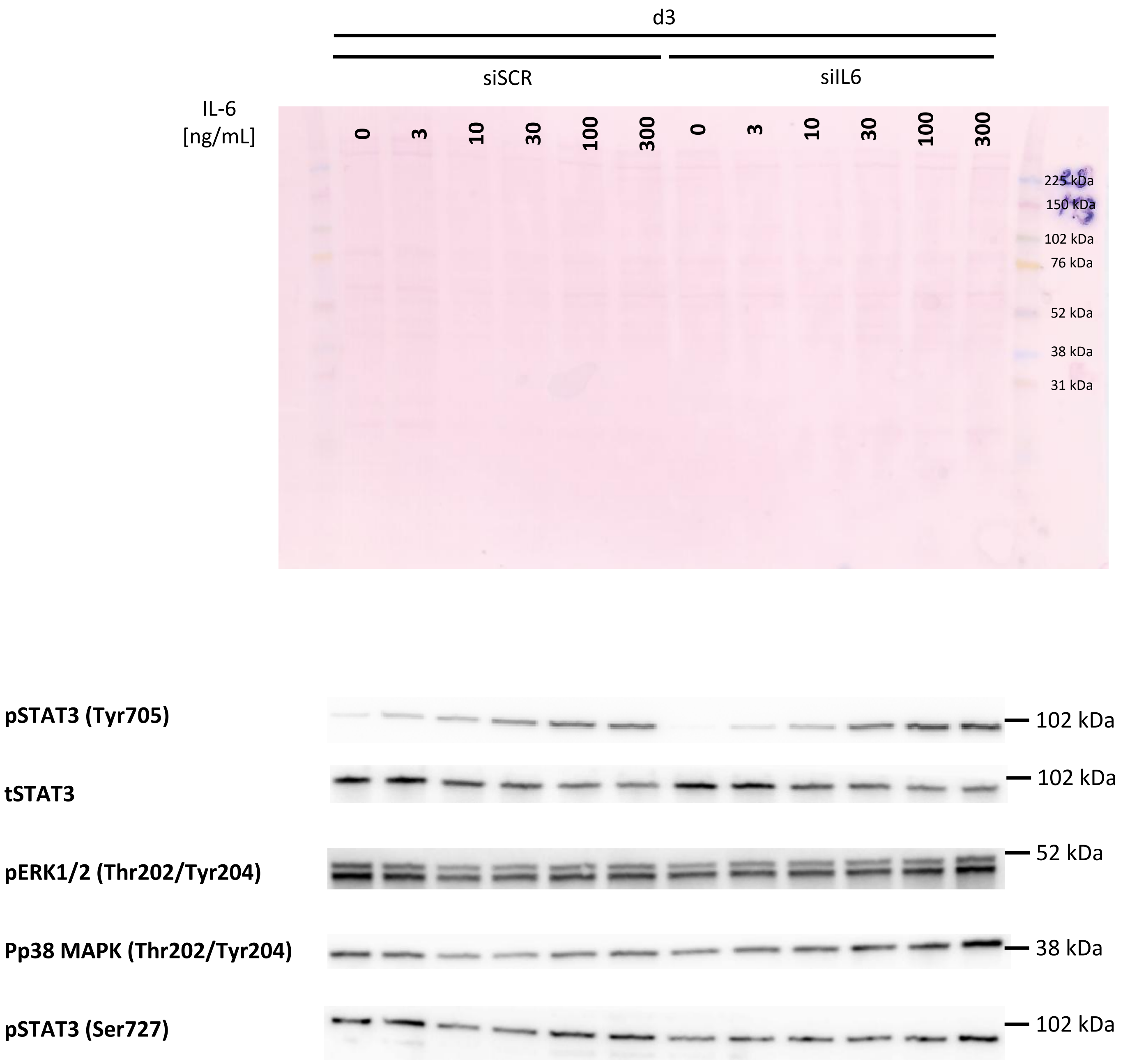
d3
siSCR
siIL6
IL-6 [ng/mL]
0
3
10
30
100
300
0
3
10
30
100
300
225 kDa
150 kDa
102 kDa
76 kDa
52 kDa
38 kDa
31 kDa
pSTAT3 (Tyr705)
102 kDa
tSTAT3
102 kDa
pERK1/2 (Thr202/Tyr204)
52 kDa
Pp38 MAPK (Thr202/Tyr204)
38 kDa
pSTAT3 (Ser727)
102 kDa

#### Figure 4: *STAT1* and *STAT3* silencing and rhIL-6 stimulation

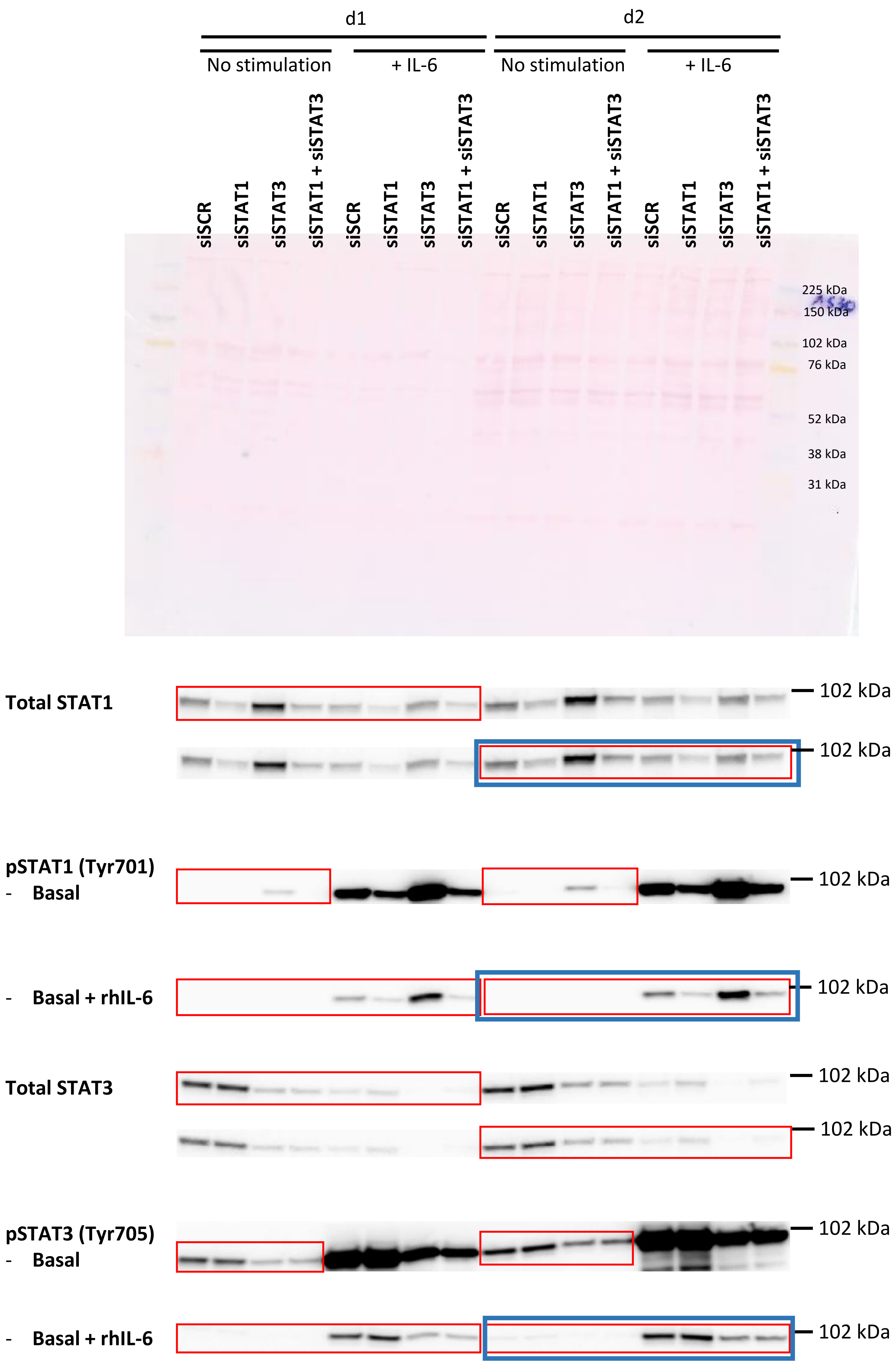

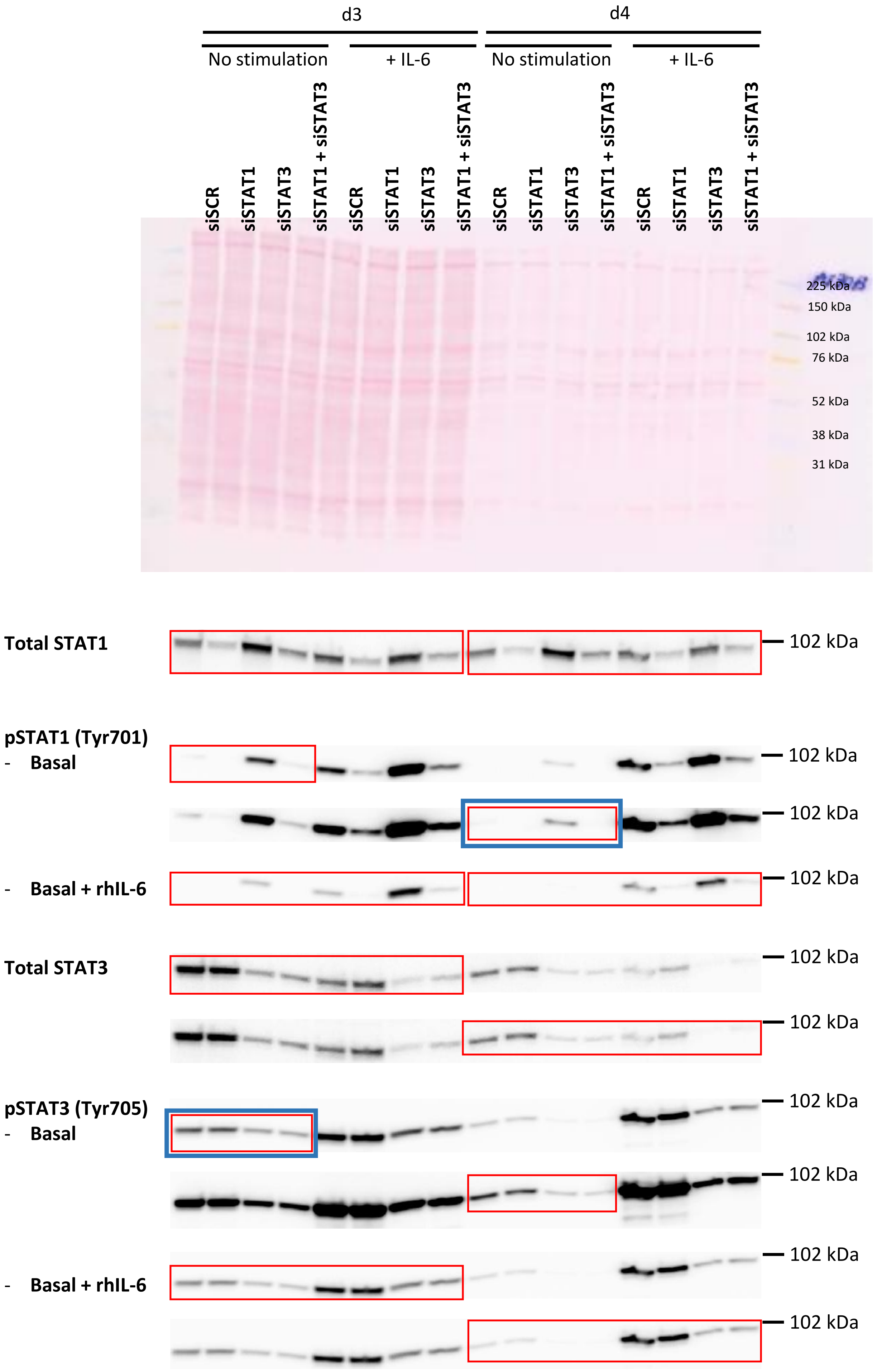

d3
d4
No stimulation
+ IL-6
No stimulation
+ IL-6
siSCR
siSTAT1
siSTAT3
siSTAT1 + siSTAT3
siSCR
siSTAT1
siSTAT3
siSTAT1 + siSTAT3
siSCR
siSTAT1
siSTAT3
siSTAT1 + siSTAT3
siSCR
siSTAT1
siSTAT3
siSTAT1 + siSTAT3
225 kDa
150 kDa
102 kDa
76 kDa
52 kDa
38 kDa
31 kDa
Total STAT1
102 kDa
pSTAT1 (Tyr701)
- Basal
102 kDa
102 kDa
- Basal + rhIL-6
102 kDa
Total STAT3
102 kDa
102 kDa
pSTAT3 (Tyr705)
- Basal
102 kDa
102 kDa
- Basal + rhIL-6
102 kDa
102 kDa

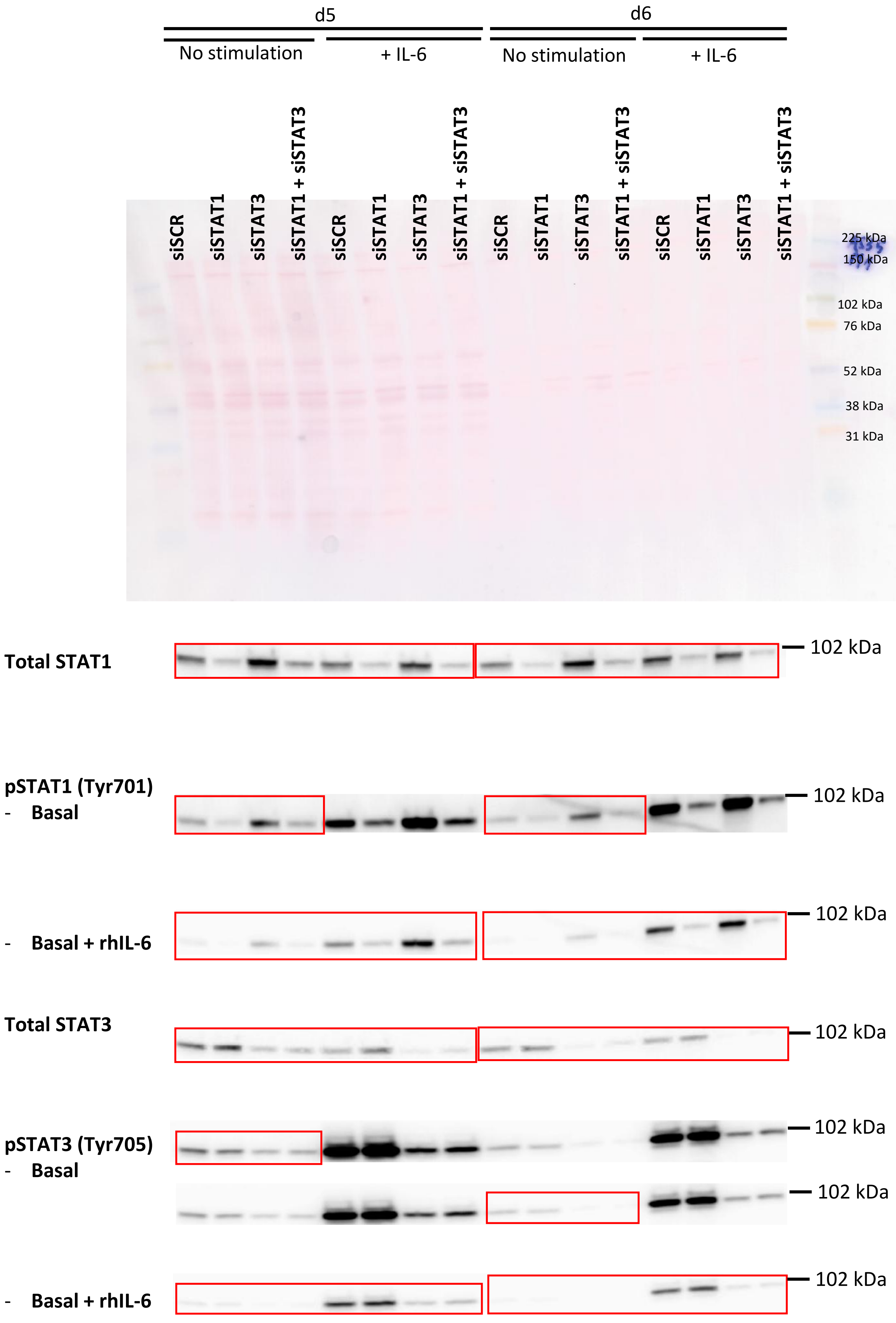
d5
d6
No stimulation
+ IL-6
No stimulation
+ IL-6
siSCR
siSTAT1
siSTAT3
siSTAT1 + siSTAT3
225 kDa
150 kDa
102 kDa
76 kDa
52 kDa
38 kDa
31 kDa
Total STAT1
pSTAT1 (Tyr701)
- Basal
- Basal + rhIL-6
Total STAT3
pSTAT3 (Tyr705)
- Basal
- Basal + rhIL-6
102 kDa

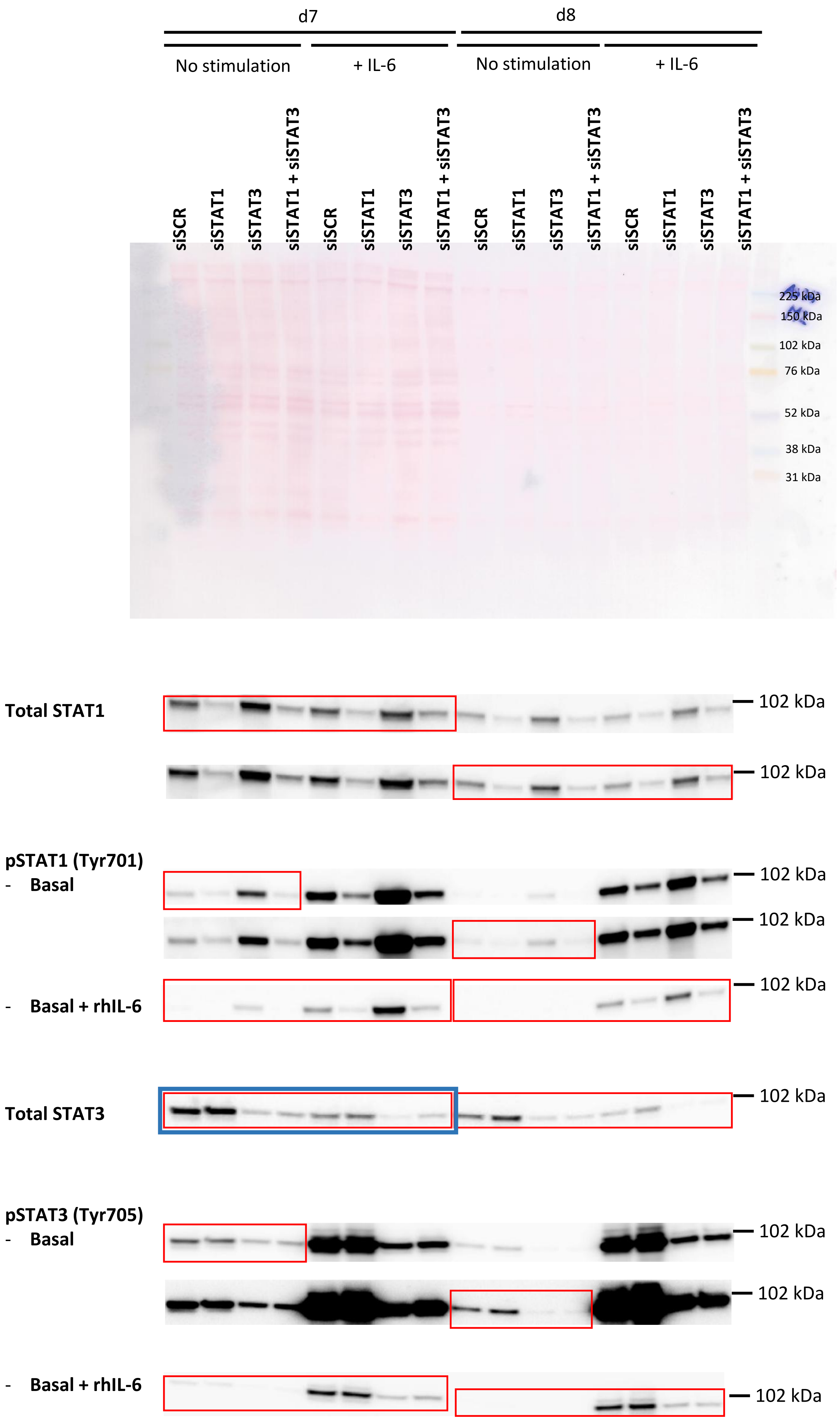
d7
d8
No stimulation
+ IL-6
No stimulation
+ IL-6
siSCR
siSTAT1
siSTAT3
siSTAT1 + siSTAT3
225 kDa
150 kDa
102 kDa
76 kDa
52 kDa
38 kDa
31 kDa
Total STAT1
102 kDa
pSTAT1 (Tyr701)
- Basal
- Basal + rhIL-6
Total STAT3
pSTAT3 (Tyr705)
- Basal
- Basal + rhIL-6

**Figure 5b: Silencing *IL6/IL6R/IL6 + IL6R* + rhIL-6 stimulation**

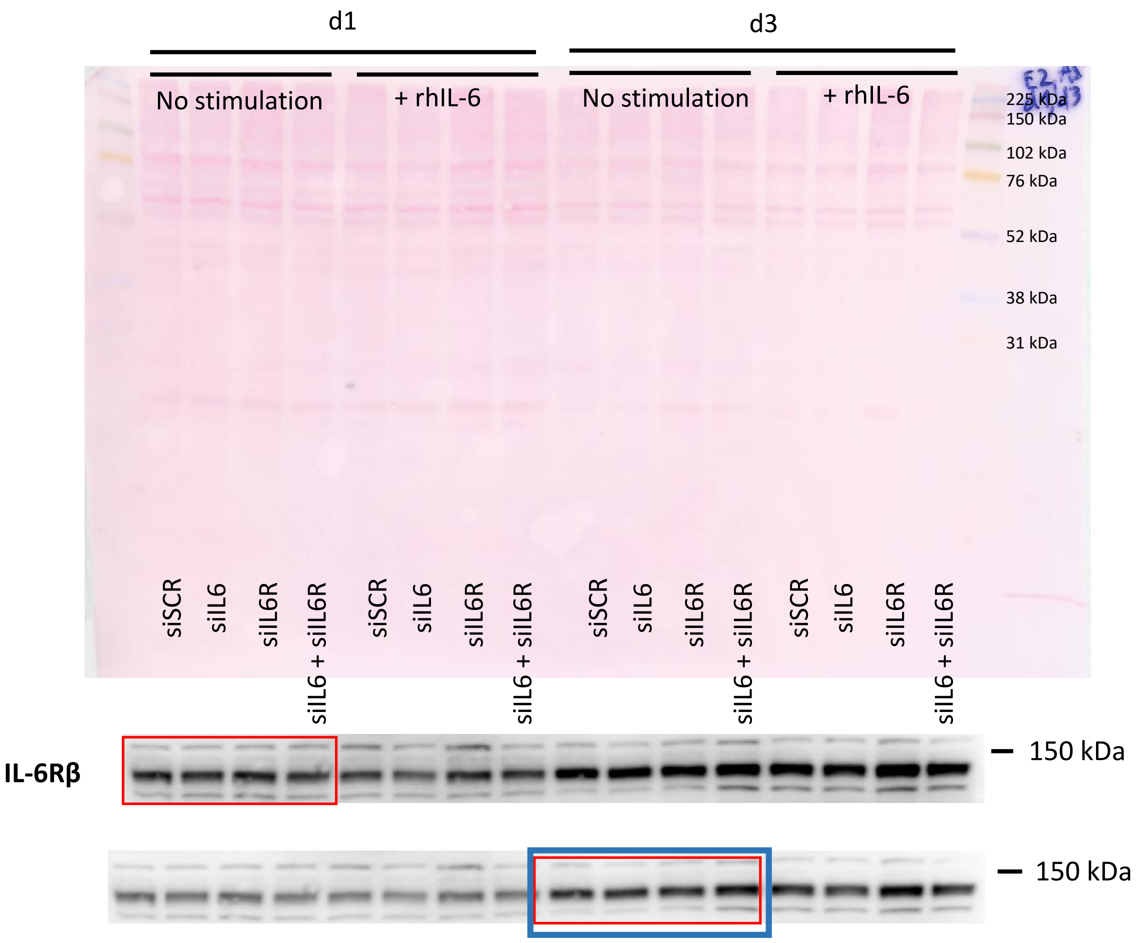


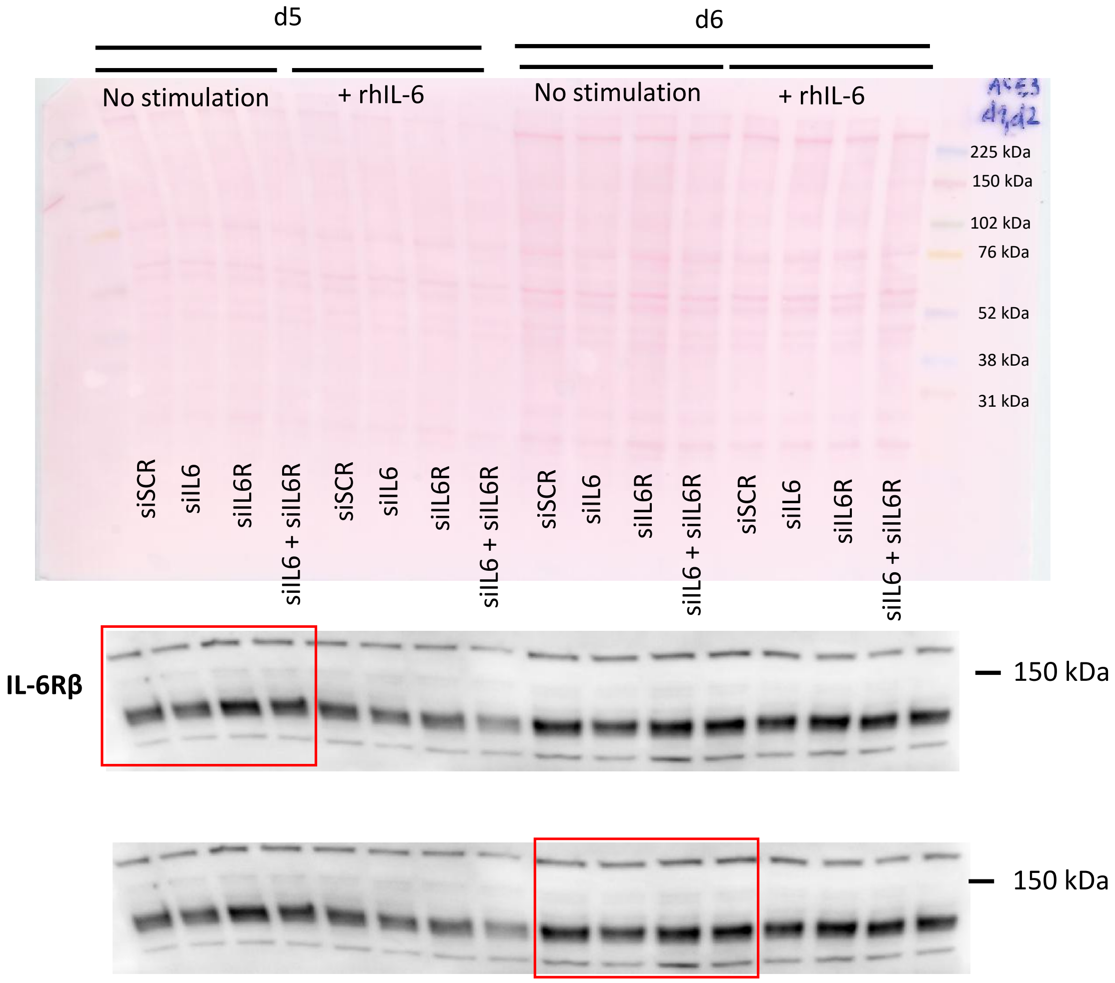

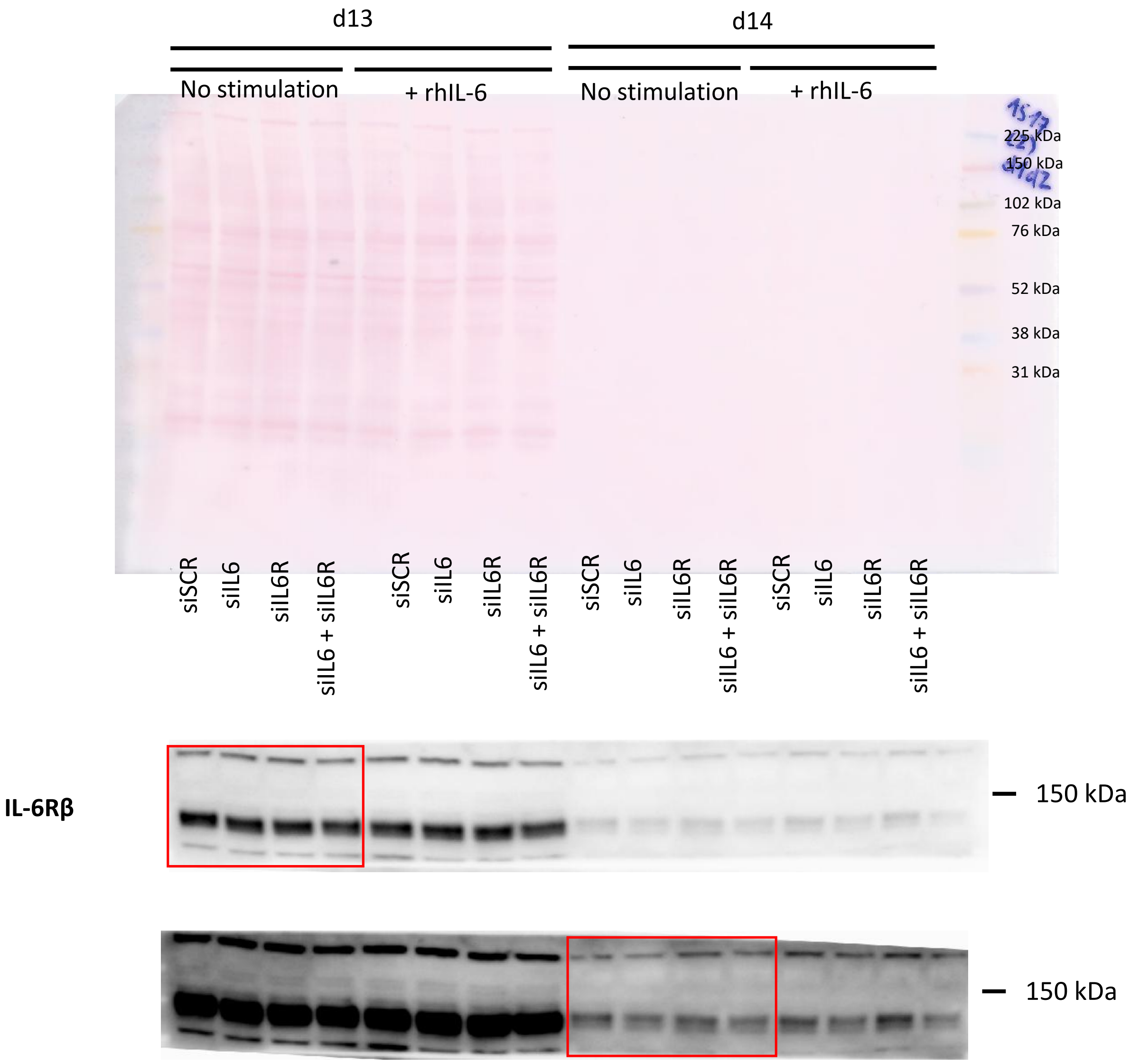
d13
d14
No stimulation
+ rhIL-6
No stimulation
+ rhIL-6
225 kDa
150 kDa
102 kDa
76 kDa
52 kDa
38 kDa
31 kDa
siSCR
siIL6
siIL6R
siIL6 + siIL6R
siSCR
siIL6
siIL6R
siIL6 + siIL6R
siSCR
siIL6
siIL6R
siIL6 + siIL6R
siSCR
siIL6
siIL6R
siIL6 + siIL6R
IL-6Rβ
150 kDa
150 kDa

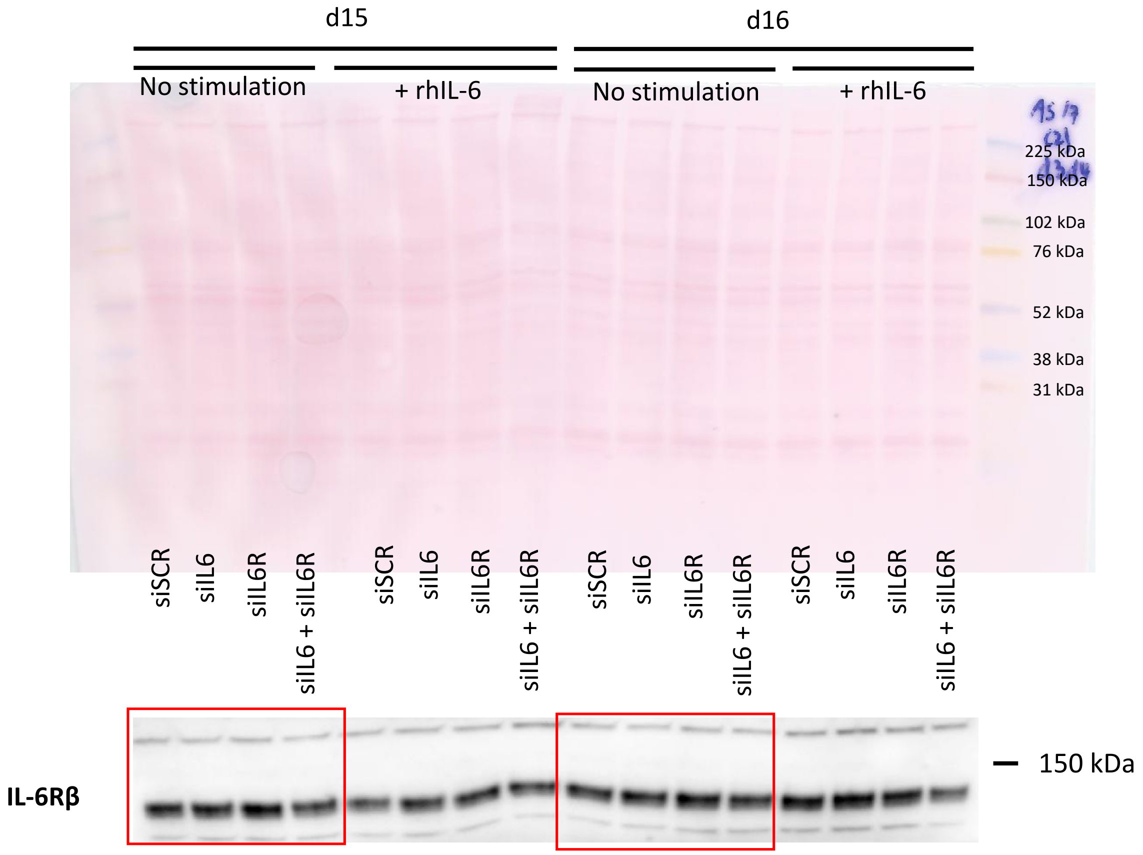
d15
d16
No stimulation
+ rhIL-6
No stimulation
+ rhIL-6
225 kDa
150 kDa
102 kDa
76 kDa
52 kDa
38 kDa
31 kDa
siSCR
siIL6
siIL6R
siIL6 + siIL6R
siSCR
siIL6
siIL6R
siIL6 + siIL6R
siSCR
siIL6
siIL6R
siIL6 + siIL6R
siSCR
siIL6
siIL6R
siIL6 + siIL6R
IL-6Rβ
150 kDa

#### Figure 5h, j: Silencing *IL6/IL6R/IL6 + IL6R* + rhLIF stimulation

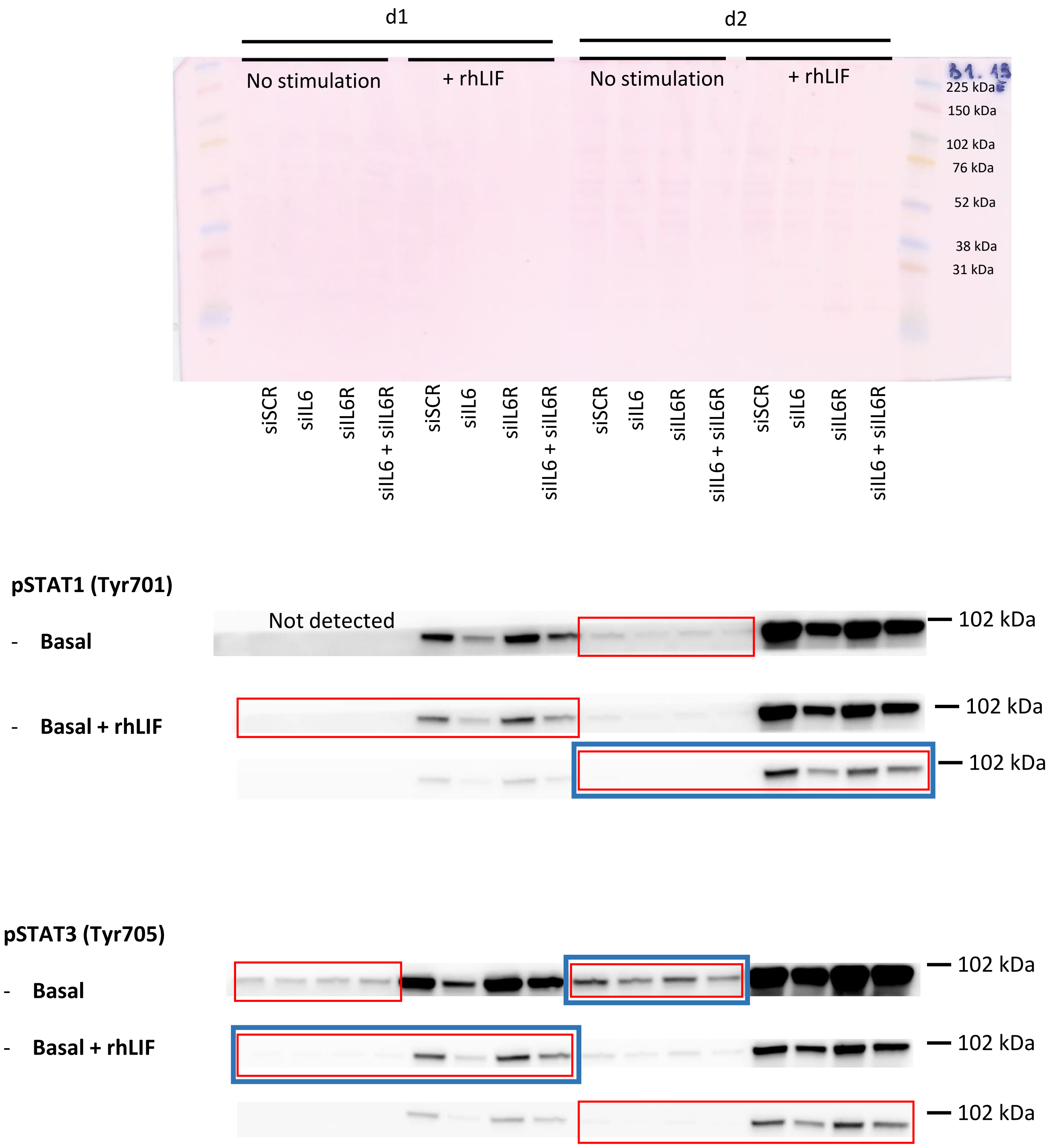

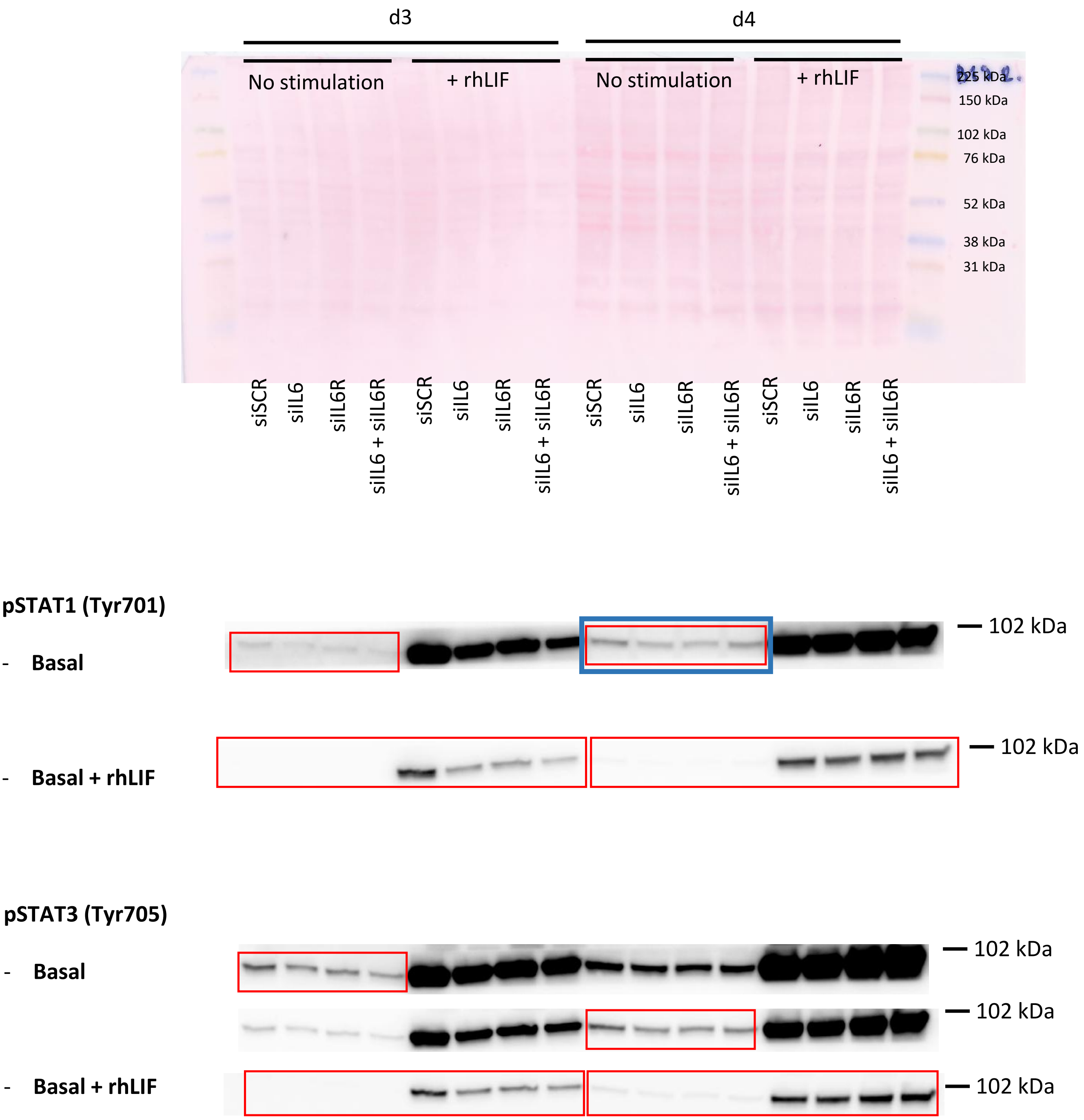
d3
d4
No stimulation
+ rhLIF
No stimulation
+ rhLIF
225 kDa
150 kDa
102 kDa
76 kDa
52 kDa
38 kDa
31 kDa
siSCR
siIL6
siIL6R
siIL6 + siIL6R
siSCR
siIL6
siIL6R
siIL6 + siIL6R
siSCR
siIL6
siIL6R
siIL6 + siIL6R
siSCR
siIL6
siIL6R
siIL6 + siIL6R
pSTAT1 (Tyr701)
- Basal
102 kDa
- Basal + rhLIF
102 kDa
pSTAT3 (Tyr705)
- Basal
102 kDa
102 kDa
- Basal + rhLIF
102 kDa

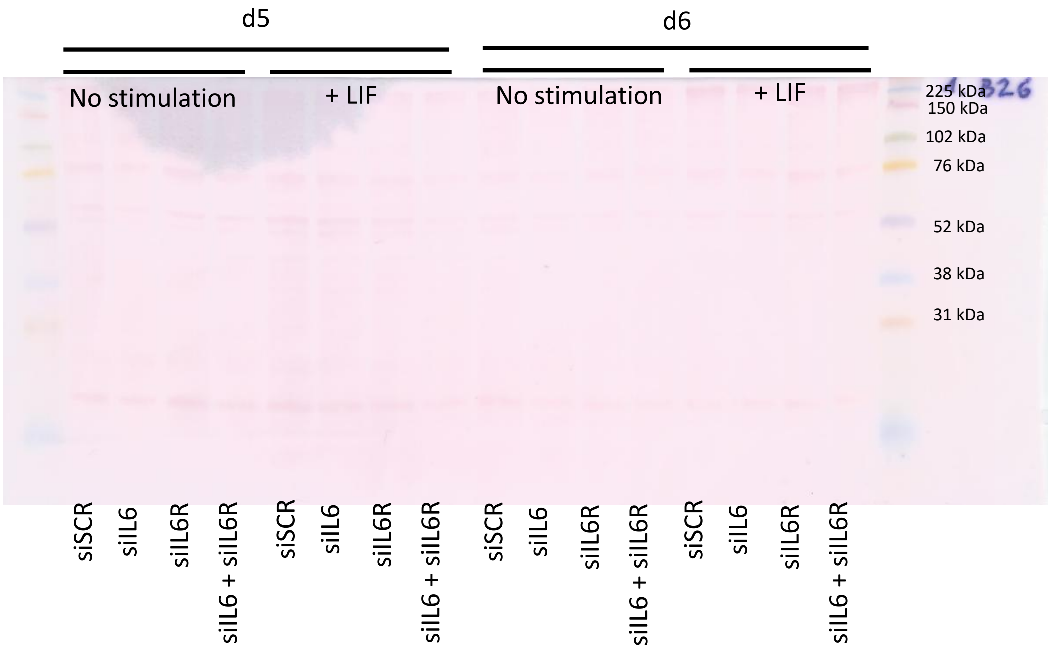
d5
d6
No stimulation
+ LIF
No stimulation
+ LIF
225 kDa
150 kDa
102 kDa
76 kDa
52 kDa
38 kDa
31 kDa
siSCR
siIL6
siIL6R
siIL6 + siIL6R
siSCR
siIL6
siIL6R
siIL6 + siIL6R
siSCR
siIL6
siIL6R
siIL6 + siIL6R
siSCR
siIL6
siIL6R
siIL6 + siIL6R

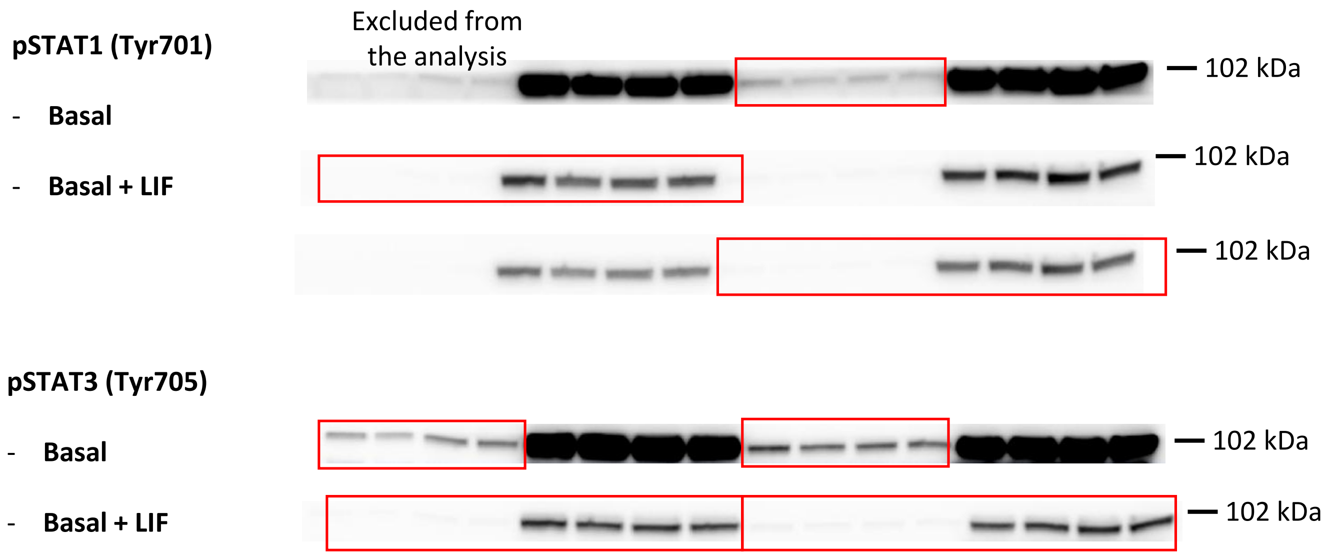
pSTAT1 (Tyr701)
- Basal
- Basal + LIF
Excluded from the analysis
102 kDa
102 kDa
102 kDa
pSTAT3 (Tyr705)
- Basal
- Basal + LIF
102 kDa
102 kDa

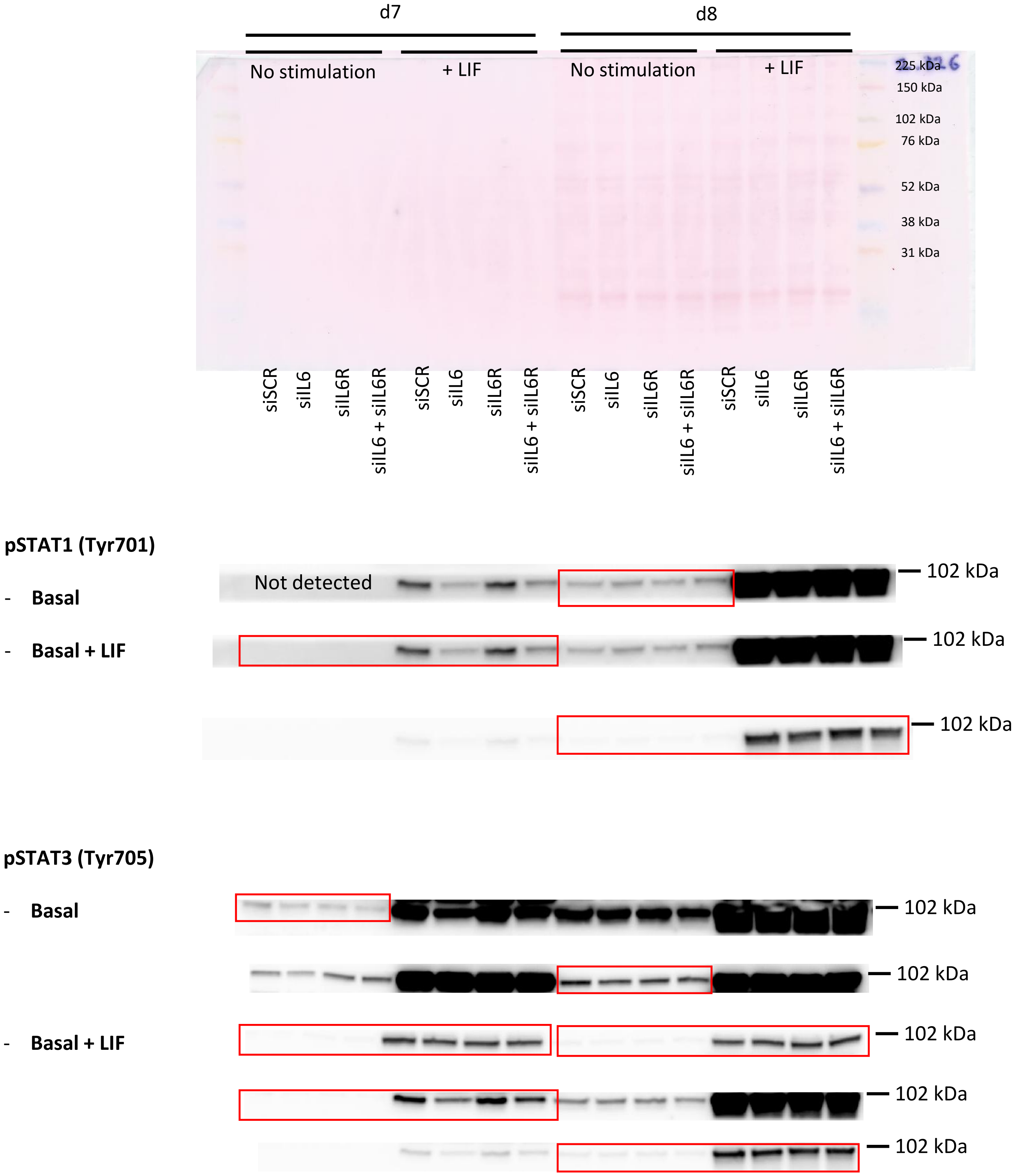
d7
d8
No stimulation
+ LIF
No stimulation
+ LIF
225 kDa
150 kDa
102 kDa
76 kDa
52 kDa
38 kDa
31 kDa
siSCR
siIL6
siIL6R
siIL6 + siIL6R
siSCR
siIL6
siIL6R
siIL6 + siIL6R
siSCR
siIL6
siIL6R
siIL6 + siIL6R
siSCR
siIL6
siIL6R
siIL6 + siIL6R
pSTAT1 (Tyr701)
- Basal
- Basal + LIF
Not detected
102 kDa
102 kDa
102 kDa
pSTAT3 (Tyr705)
- Basal
- Basal + LIF
102 kDa
102 kDa
102 kDa
102 kDa
102 kDa

#### Figure 6a, b: Neutralising antibody experiments

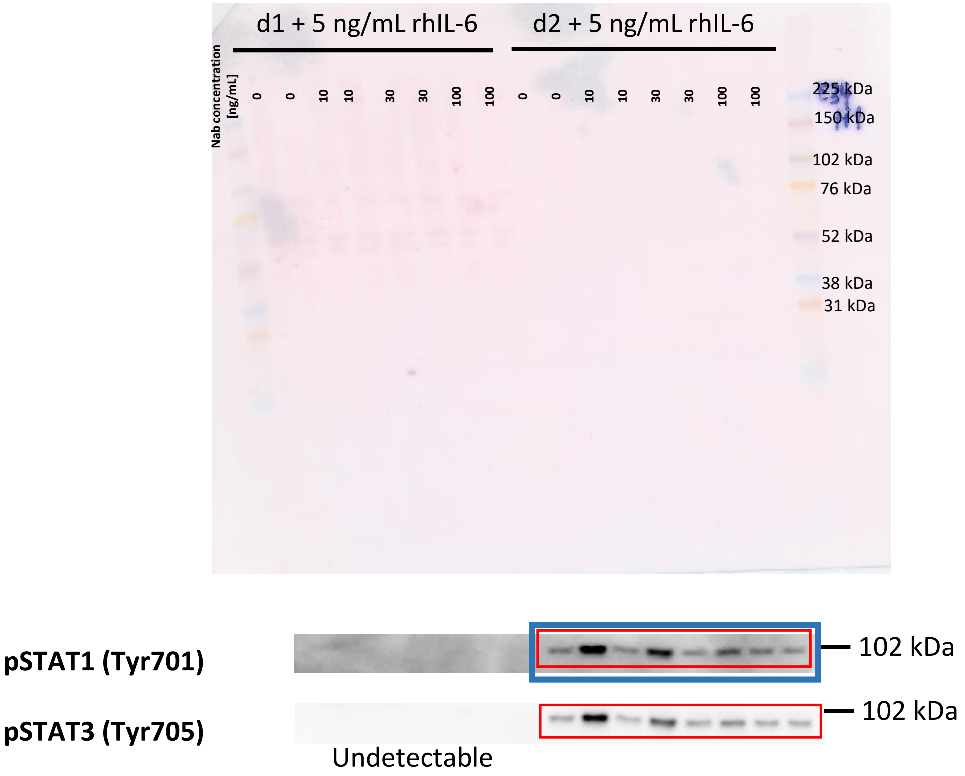


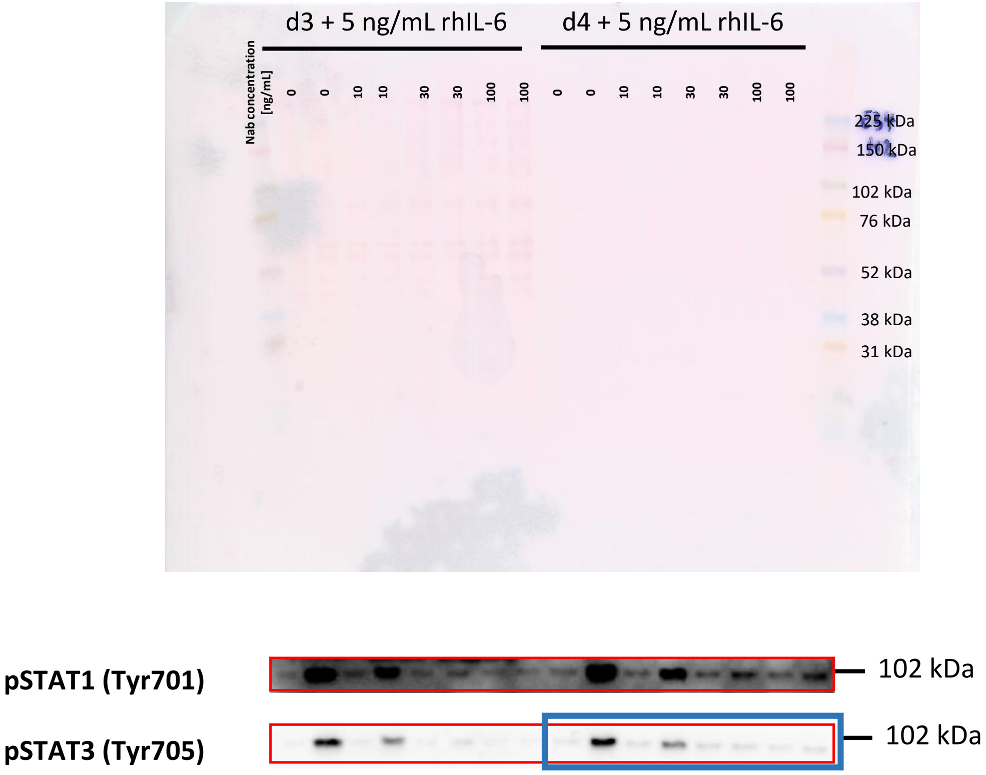

**Figure 6c, d: Neutralising antibody experiments**

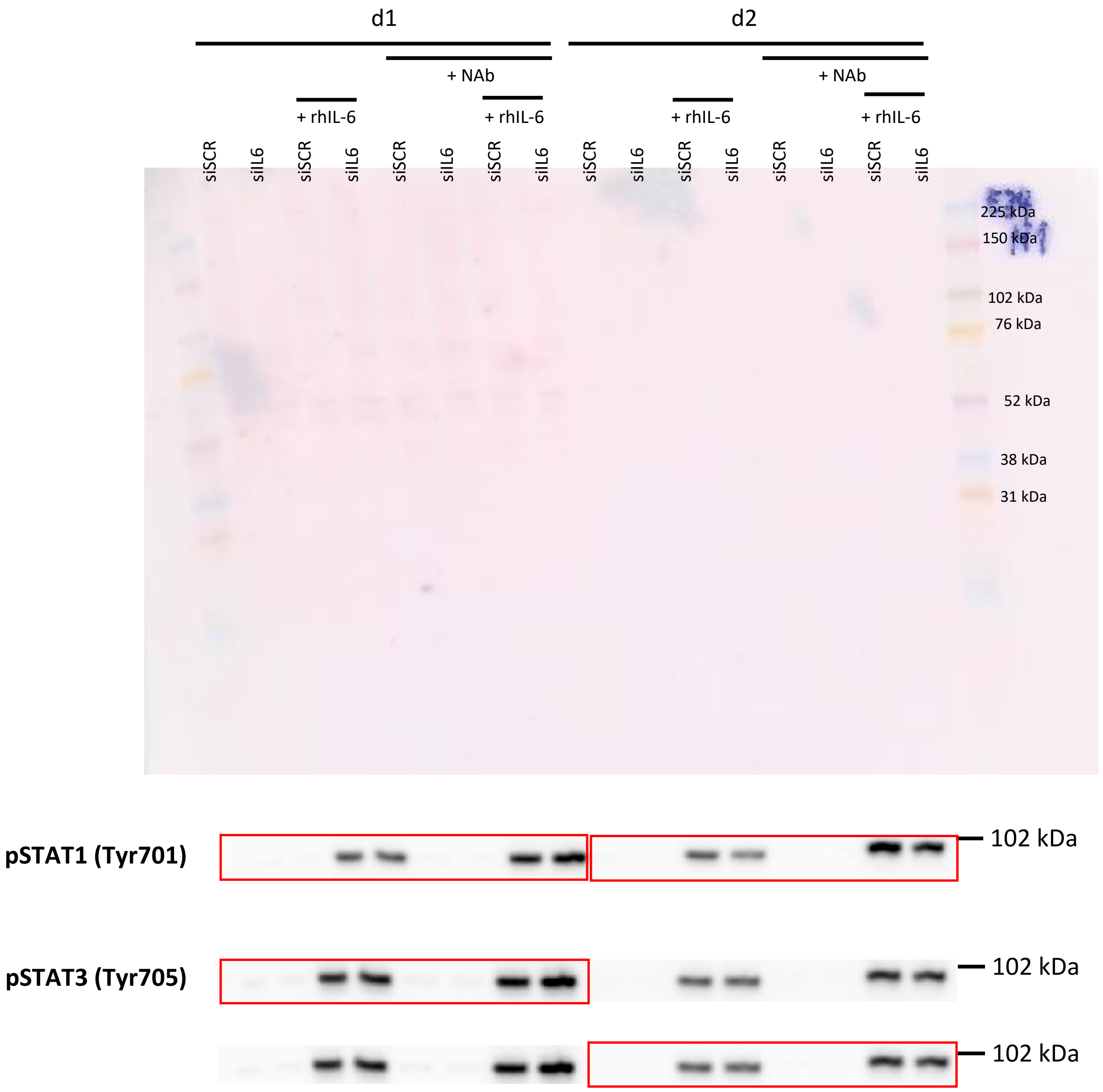

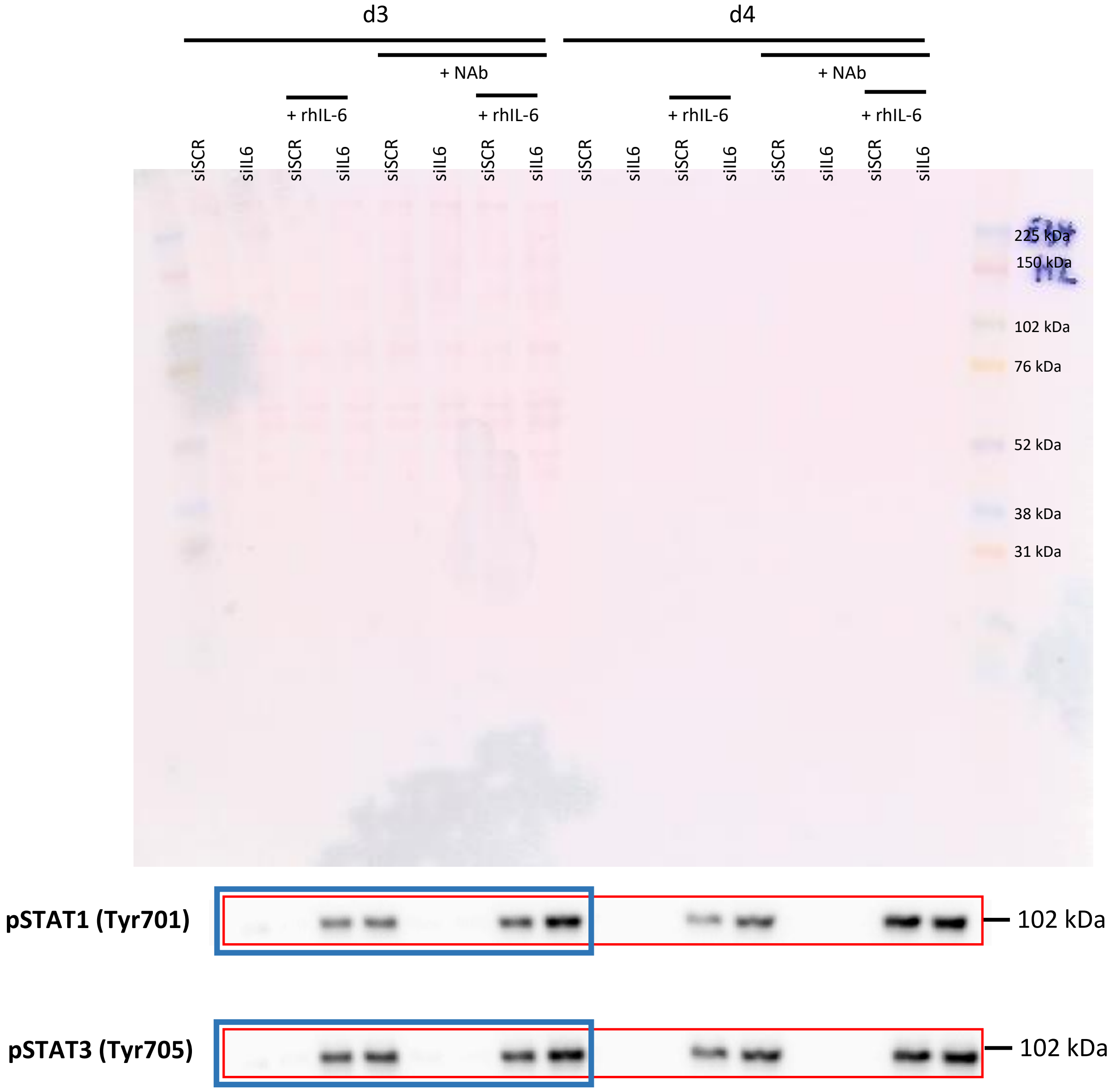
d3
d4
+ NAb
+ NAb
+ rhIL-6
+ rhIL-6
+ rhIL-6
+ rhIL-6
siSCR
siIL6
siSCR
siIL6
siSCR
siIL6
siSCR
siIL6
siSCR
siIL6
siSCR
siIL6
siSCR
siIL6
siSCR
siIL6
225 kDa
150 kDa
102 kDa
76 kDa
52 kDa
38 kDa
31 kDa
pSTAT1 (Tyr701)
102 kDa
pSTAT3 (Tyr705)
102 kDa

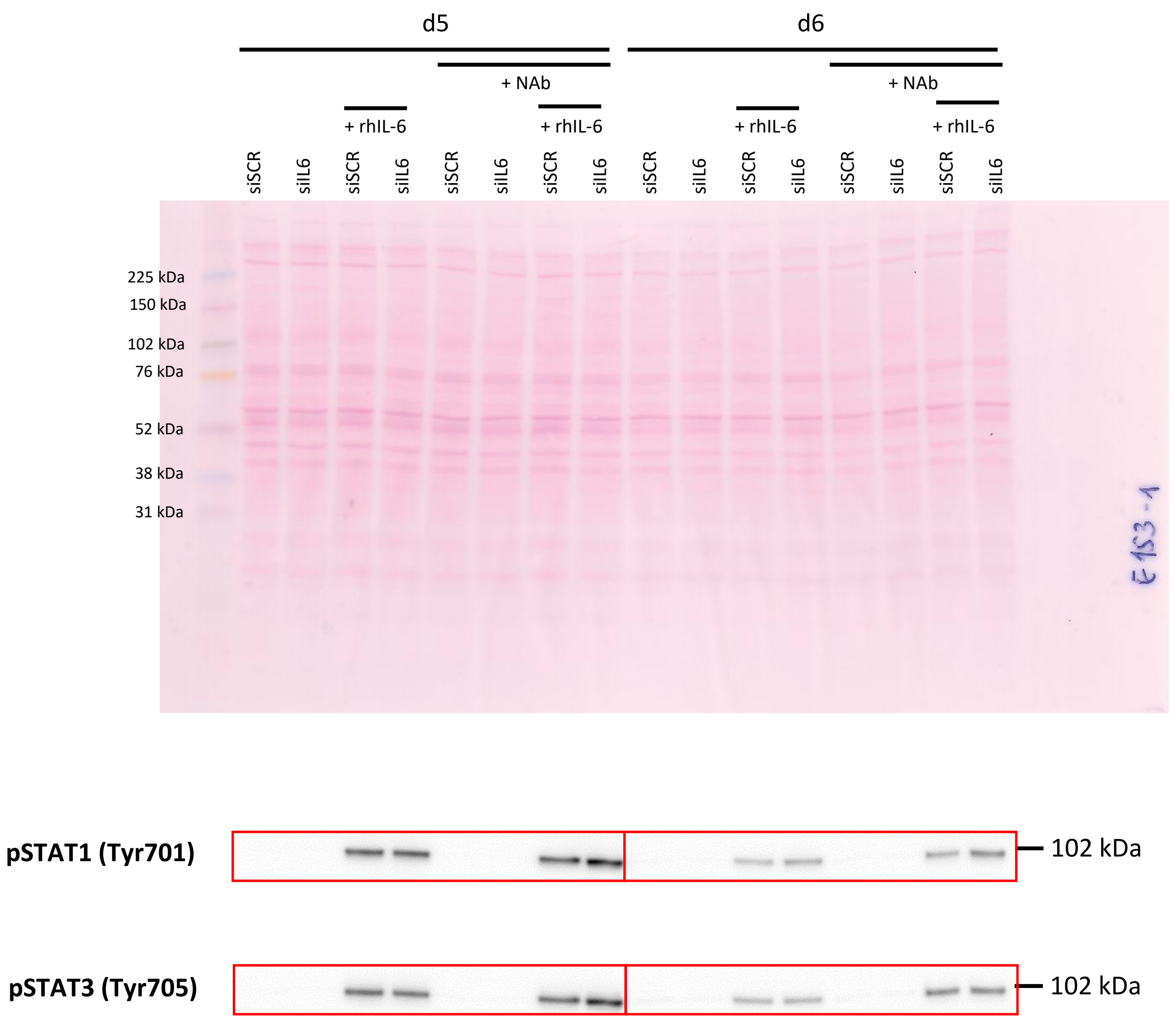
d5
d6
+ NAb
+ NAb
+ rhIL-6
+ rhIL-6
+ rhIL-6
+ rhIL-6
siSCR
siIL6
siSCR
siIL6
siSCR
siIL6
siSCR
siIL6
siSCR
siIL6
siSCR
siIL6
siSCR
siIL6
siSCR
siIL6
225 kDa
150 kDa
102 kDa
76 kDa
52 kDa
38 kDa
31 kDa
E153-1
pSTAT1 (Tyr701)
102 kDa
pSTAT3 (Tyr705)
102 kDa

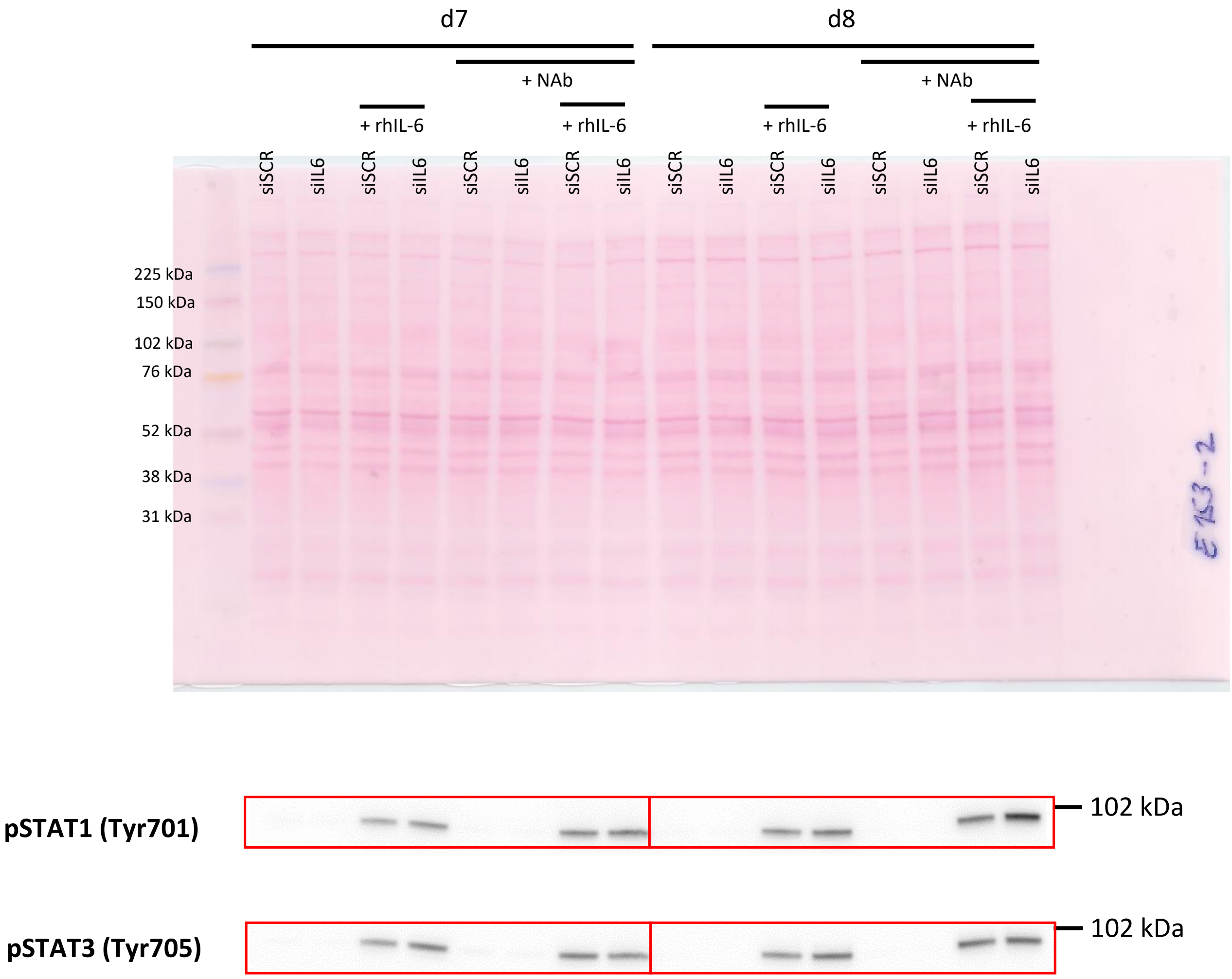
d7
d8
+ NAb
+ NAb
+ rhIL-6
+ rhIL-6
+ rhIL-6
+ rhIL-6
siSCR
siIL6
siSCR
siIL6
siSCR
siIL6
siSCR
siIL6
siSCR
siIL6
siSCR
siIL6
siSCR
siIL6
siSCR
siIL6
225 kDa
150 kDa
102 kDa
76 kDa
52 kDa
38 kDa
31 kDa
pSTAT1 (Tyr701)
102 kDa
pSTAT3 (Tyr705)
102 kDa

#### Figure 6e, f: pretreatment with rhIL-6

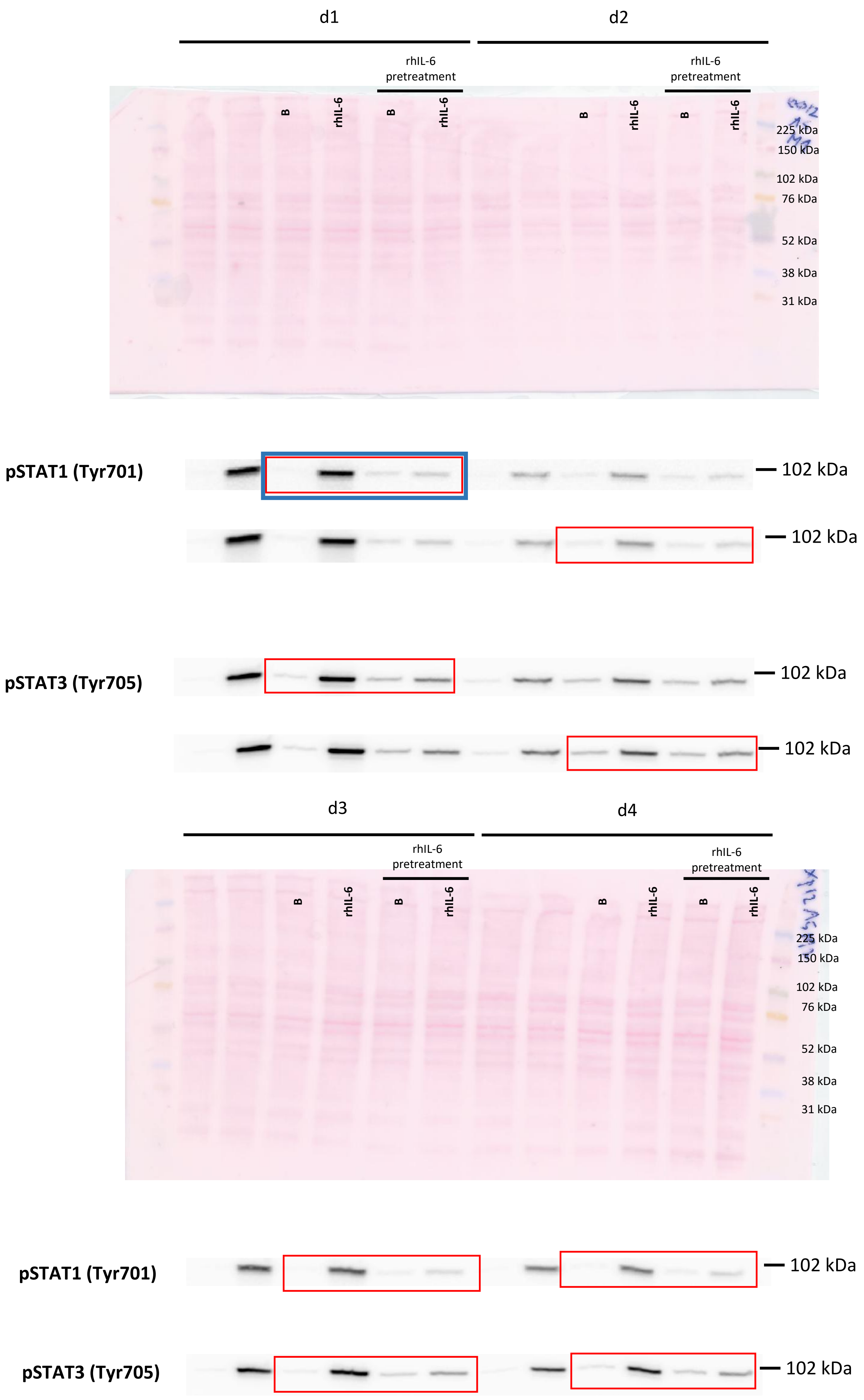

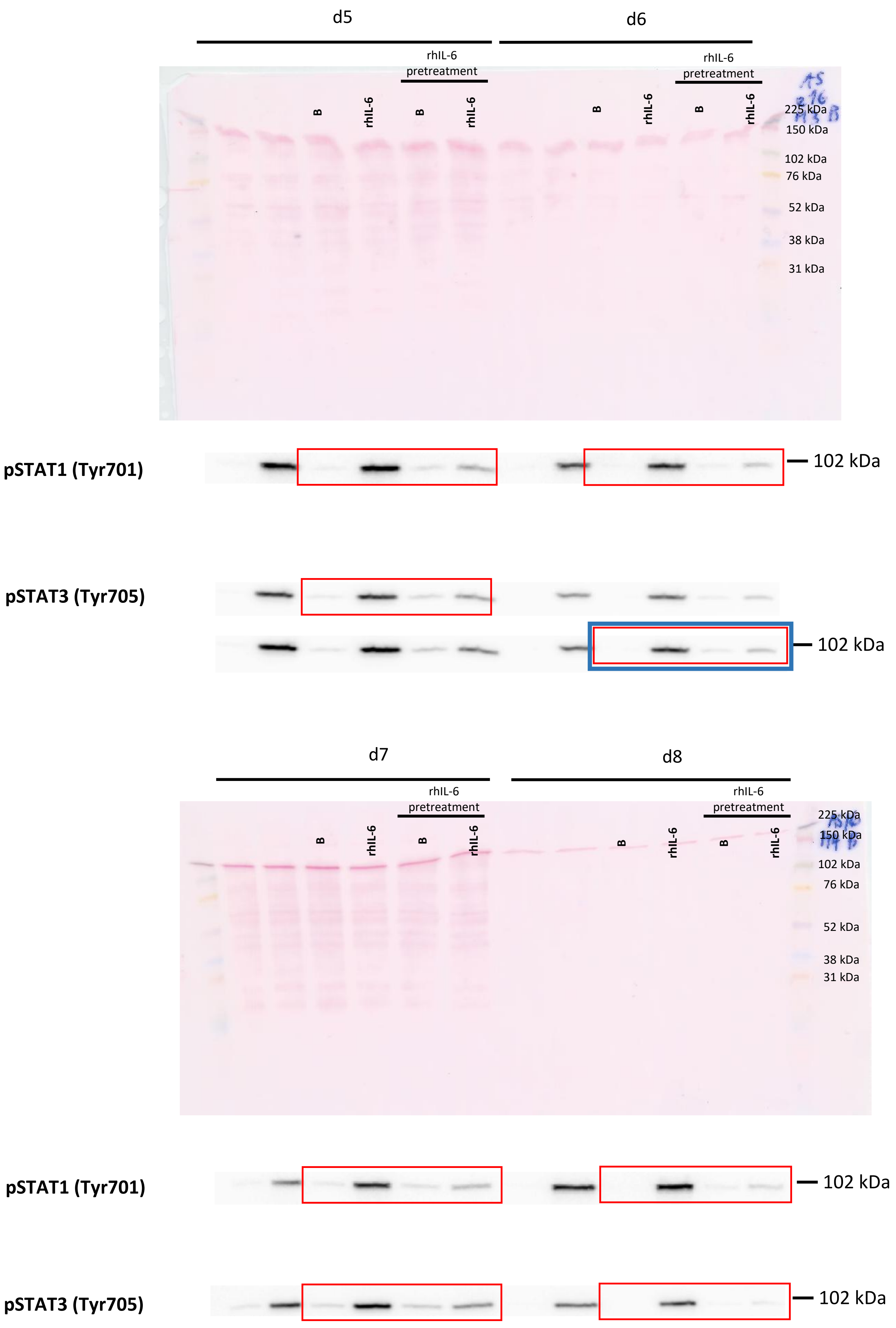

d5
d6
rhIL-6 pretreatment
B
rhIL-6
B
rhIL-6
B
rhIL-6
B
rhIL-6
225 kDa
150 kDa
102 kDa
76 kDa
52 kDa
38 kDa
31 kDa
pSTAT1 (Tyr701)
102 kDa
pSTAT3 (Tyr705)
102 kDa
d7
d8
rhIL-6 pretreatment
B
rhIL-6
B
rhIL-6
B
rhIL-6
B
rhIL-6
225 kDa
150 kDa
102 kDa
76 kDa
52 kDa
38 kDa
31 kDa
pSTAT1 (Tyr701)
102 kDa
pSTAT3 (Tyr705)
102 kDa
